# An updated structural model of the A domain of the *Pseudomonas putida* XylR regulator exposes a distinct interplay with aromatic effectors

**DOI:** 10.1101/2021.01.17.427014

**Authors:** Pavel Dvořák, Carlos Alvarez-Carreño, Sergio Ciordia, Alberto Paradela, Víctor de Lorenzo

## Abstract

A revised model of the aromatic binding A domain of the σ^54^-dependent regulator XylR of *Pseudomonas putida* mt-2 was produced based on the known 3D structures of homologous regulators PoxR, MopR, and DmpR. The resulting frame was instrumental for mapping the large number of mutations known to alter effector specificity, which were then reinterpreted under a dependable spatial reference. Some of these changes involved the predicted aromatic-binding pocket but others occurred in distant locations, including dimerization interfaces and putative zinc-binding site. The effector pocket was buried within the protein structure and accessible from the outside only through a narrow tunnel. The model was experimentally validated by treating the cells *in vivo* and the purified protein *in vitro* with benzyl bromide, which reacts with accessible nucleophilic residues on the protein surface. Proteomic analyses of the thereby tagged peptides confirmed the predicted in/out distribution of residues but also suggested that the fully-folded protein is not accessible by externally added effectors. The data thus suggested that XylR inducers assist the folding and/or the structuring of the A domain in an intramolecular non-repressive form rather than interacting dynamically with the aromatic partner once a fully structured protein is shaped.

**Originality-Significance Statement:** XylR is a transcriptional regulator of *Pseudomonas putida* strain mt-2 which activates the *upper T*OL pathway promoter *Pu* for catabolism of toluene and *m*-xylene upon binding of these aromatic effectors to its N-terminal A domain. While this feature has made XylR a popular platform for the development of whole-cell biosensors for aromatic compounds, the difficulty to crystallize the A domain —let alone the whole-length protein— has made structural comprehension of the effector-regulator binding quite problematic. To overcome this impasse, we have combined homology-based structural predictions of the A domain of XylR with biochemical probing of exposed amino acids on the surface of the protein, both *in vivo* and *in vitro*. The results generally matched the effects of mutations known from previous genetic/phenotypic analyses of the protein. However, the data also suggested an intriguing mechanism of activation of XylR by effectors in which the inducer assists the shaping of the regulator in an active conformation rather than interacting *a posteriori* with an already formed protein *invitro*. This may in fact explain the longstanding failure to purify the protein in an effector-responsive form.

## INTRODUCTION

Operons encoding enzymatic routes typically found in environmental bacteria for biodegradation of aromatic environmental pollutants are often regulated by transcriptional factors (TFs) directly responsive to the pathway substrates themselves or to metabolic intermediates of the catabolic process (Shingler, 2003; Galvão and de Lorenzo, 2006). One conspicuous class of such factors belongs to the so-called NtrC superfamily of bacterial enhancer-binding proteins (EBPs) that act at a distance on cognate promoters in concert with the σ^54^-containing form of RNA polymerase (Weiss *et al.*, 1992; North *et al.*, 1993). The conserved structure of EBPs comprises 3 distinct domains (**Fig. 1**), the N-terminal of which (the so-called A domain) being the one that receives the environmental signal that turns on the protein to become an effective transcriptional activator. In a subset of EBPs, the A domain inhibits the interaction of the central domain of the regulator with the σ^54^-dependent transcription initiation complex. Binding of the aromatic effector to the A domain of the EBP at stake relieves the intramolecular repression thereby triggering transcription initiation (**Fig. 1D**; Pérez-Martín and de Lorenzo, 1995). Archetypal TFs of this sort include the XylR protein, which regulates the two σ^54^-dependent promoters found in the *xyl* operons for degradation of *m*-xylene borne by TOL plasmid pWW0 of *Pseudomonas putida* mt-2 (**Fig. 1**; Abril *et al.*, 1989). That the interaction of the aromatic effector with the TF is localized in a small protein segment that can be swapped with homolog moieties with other specificities in similar EBPs had made the A moiety of XylR an appealing platform for developing biosensors for a variety of aromatic compounds (Galvão and de Lorenzo, 2006; Huang *et al.,* 2008) including metabolites and explosives (Garmendia *et al.*, 2008; de Las Heras and de Lorenzo, 2011). Alas, this endeavour has been recurrently curbed by the lack of a reliable structural reference which has not only prevented directed mutagenesis of the residues involved in effector recognition, but also a mechanistic understanding of the many XylR mutants responsive to non-native inducers. This is because of the difficulty to purify the intact protein in a native form. ΔA truncated variants of XylR which activate constitutively the target σ^54^-promoters of the TOL plasmid (**Fig. 1**) can be purified in large amounts. But to this day, purification of full-length XylR (or its separate A domain) in an active form has been challenging owing to the tendency of the product to form insoluble inclusion bodies (Pérez-Martin *et al.*, 1997; Kim *et al.*, 2005).

**Figure 1.**
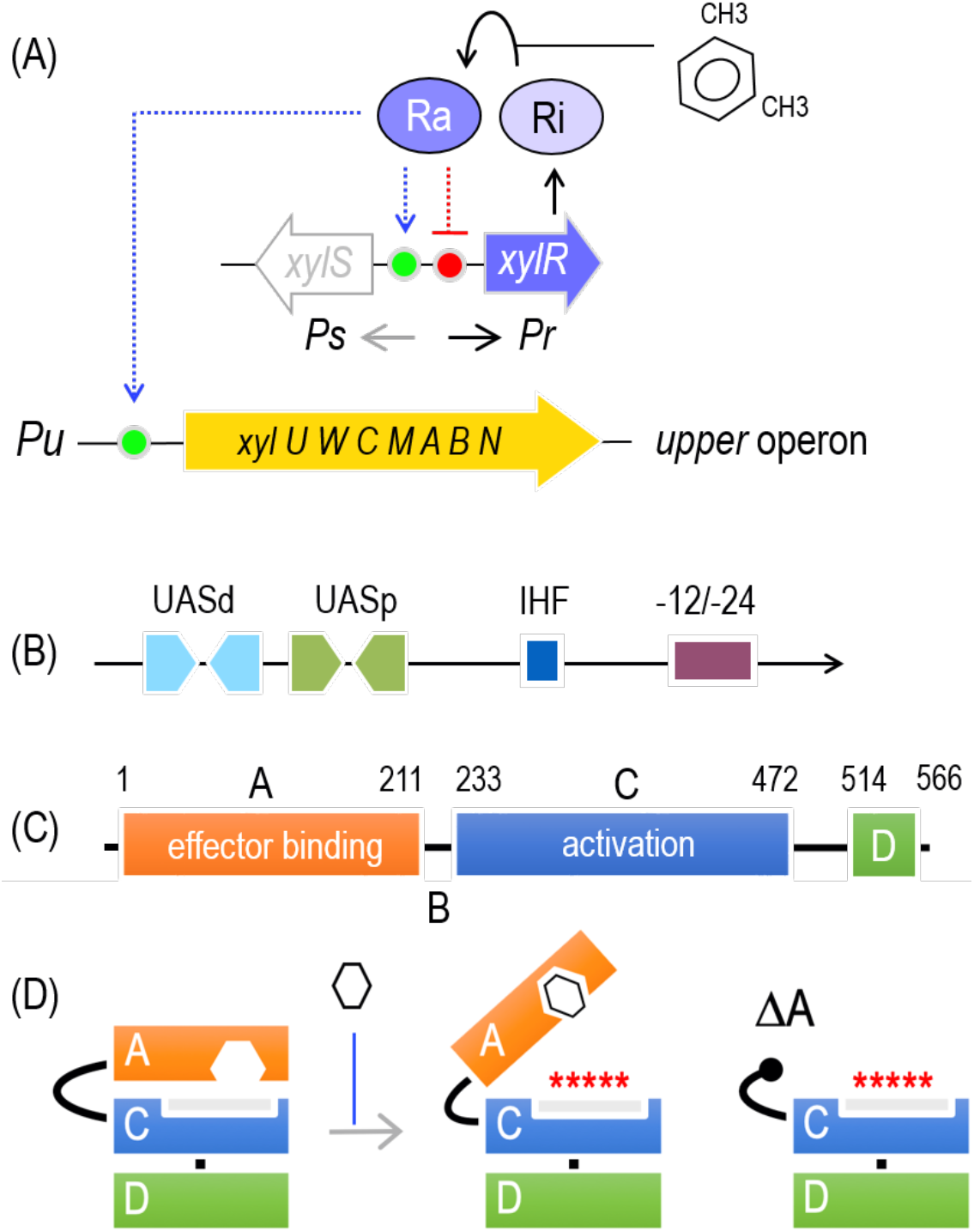
Biological and regulatory context of XylR and role of its A domain. (A) The products of the *upper* TOL pathway encoded by plasmid pWW0 transform *m*-xylene into 3-methylbenzoate, and the *lower* operon (not shown) that produces enzymes for further metabolism of this compound into TCA cycle intermediates. XylR and XylS are the transcriptional regulators that control the expression of either operon. The *xylR* is expressed from the *Pr* promoter and XylR produced in an inactive form (Ri) that, in the presence of the pathway substrate (*m*-xylene) changes to an active form (Ra). Ra activates both *Pu* and *Ps*, triggering expression of the *upper* pathway and XylS, respectively. Ra also acts as a repressor of its own transcription. Operons and regulatory elements not to scale. (B) The *Pu* promoter region. The DNA segment of interest is expanded, showing the location of distal and proximal upstream binding sites for XylR (UASd and UASp), the −12/-24 motif recognized by σ^54^-RNAP, and one integration host factor (IHF) binding site located in between. (C) Functional domains of XylR. The sketch indicates amino acid residues at the limits between the functional domains and the localization of the relevant functions within the protein sequence. (D) Activation of XylR by *m*-xylene. The TF folds such that the N-terminal A domain hinders an activation surface of the regulator. Effector binding to the A domain releases the inhibition and XylR is then able to activate σ^54^-RNAP. Deletion of the whole A domain originates an effector-independent, constitutively active variant of XylRΔA.

As a way to overcome these limitations, a structural model of XylR A domain was proposed by Devos *et al.* (2002) using the best bioinformatic tools for protein threading available at the time (**Supplementary Fig. S1**). The model was based on [i] crystallographic data collected for catechol o-methyltransferase (COMT: PDB ID: 1VID; Vidgren *et al.*, 1994), [ii] distribution of characteristic residues in sequences from related families (XylR, DmpR, or HbpR among others), nonpolar determinants and correlating mutations (multiple sequence alignment), and [iii] physico-chemical properties of the conserved amino acids in proteins from the family of σ^54^-dependent EBPs and the COMT enzyme. The XylR structural model had 207 residues and had the shape of a typical Rossmann fold architecture with eight α helices and seven β strands. The binding cavity was proposed to be formed by loops and residues M37, F65, E140, and E172. The remaining residues of mostly hydrophobic character were proposed to stabilise aromatic ligand in the binding site and define the specificity of XylR (Delgado and Ramos, 1994; Shingler and Pavel, 1995; Garmendia *et al.*, 2001). This model was for years the only one available to interpret XylR mutations and make sense of their phenotypes. Yet, it became obvious from the beginning that the structural basis of many mutants could not be easily recognized in the model and therefore a better one was badly needed.

Fortunately, in 2016 crystal structures were published of the A domains of the EBPs and XylR homologues PoxR and MopR (Patil *et al.*, 2016; Ray *et al.*, 2016) that activate σ^54^-promoters for phenol-degradation operons of *Ralstonia eutropha* and *Acinetobacter calcoaceticus,* respectively. The structure of the A domain of the archetypal dimethylphenol-responsive σ^54^-activator DmpR from *P. putida* has recently been published as well (Park *et al.*, 2020). Alignment of the N-terminal A-domains (211 amino acids) of these proteins (**Supplementary Fig. S2**) shows a high sequence identity among the four proteins. Furthermore, that the A domains of DmpR and XylR can be swapped without any loss of function other than the exchange of effector specificity (i.e. *m*-xylene vs. phenol; Shingler and Moore, 1994) indicates the likeness of their respective tertiary structures. This state of affairs has enabled us to revise the structure of the A domain of XylR with the reliable frame of PoxR, MopR, and DmpR.

In this work, we have developed and studied a model of the effector-binding moiety of XylR that both overcomes the shortcomings of that of Devos *et al.* (2002) and provides a structural rationale for many of the incomprehensible phenotypes of mutants generated in the past on this transcriptional factor. Furthermore, we provide genetic and biochemical evidence in support of the proposed tridimensional shape of the domain. Finally, we show data suggesting that aromatic effectors of XylR access the protein *in vivo* before it is fully folded into an otherwise non-receptive and transcriptionally dead form—rather than binding the mature TF *in vitro*.

## RESULTS

### Modelling of XylR A domain structure by molecular threading

XylR A domain shows 40 %, 41 %, and 46 % sequence identity with ligand recognition domains of PoxR, MopR, and DmpR respectively (**Supplementary Fig. S2**). Indeed, crystal structures of PoxR (PDB ID: 5FRU and 5FRV), MopR (PDB ID: 5KBE), and DmpR (PDB ID: 6IY8) sensory domains (or whole EBP in case of DmpR) obtained in high resolution (1.85, 1.90, 2.50, and 3.42 Å, respectively) appeared repeatedly among top 10 templates selected based on their significance from the LOMETS threading programs during I-TASSER calculations. The phenol-responsive sensory domain of PoxR (PDB ID: 5FRU) was, with 93 % coverage and TM score of 0.91, structurally closest to the modelled XylR A domain. The confidence of each model generated by I-TASSER is quantitatively measured by C-score which is typically in the range of −5 to + 2 (Yang and Zhang, 2015). The higher the value, the better is the model. C-score and RMSD (root-mean-square deviation of atomic positions) of the top-ranked model of XylR A domain were 0.93 and 3.6 ± 2.5 respectively, which signs highly accurate prediction.

The top-ranked model (**Fig. 2**), used for further work, shows typical structural features previously described for PoxR, MopR, and DmpR proteins (Patil *et al.*, 2016; Ray *et al.*, 2016; Park *et al.*, 2020). The XylR A domain is formed by a mixed α/β fold of seven α helices and seven β strands. N-terminal part consists of two α helices (α1 and α2) and three-stranded antiparallel β sheet between them. In PoxR, MopR, and DmpR this part of the sensory domain together with helix α5 play a crucial role in protein dimerization. It can also be assumed that XylR A domain forms a dimer. The core part of the domain comprises a four-stranded antiparallel β sheet (β4-β7) and a bundle of three α helices (α3, α4,α6). The binding pocket for aromatic ligands (**Fig. 2**), as predicted by I-TASSER and verified by CAVER web (Stourac *et al.*, 2019), lies in between and is formed by the residues F93, G96, P97, Y100, V108, V124, A126, W128, Y155, A156, Y159, F170, and I185 which originate in all four β strands and in helices α4 and α6. The calculated volume of the pocket is 209±12 Å^3^. Seven (G96, P97, V108, W128, Y155, A156, and Y159) out of the thirteen, mostly hydrophobic, residues are conserved among XylR, PoxR, MopR, and DmpR (**Supplementary Fig. S2**). Position 124 is variable but in all three proteins is occupied by a small hydrophobic residue. Positions 100, 126, and 170 are occupied by histidine, phenylalanine, and tyrosine, respectively, in PoxR and MopR, or by histidine, methionine, and phenylalanine, respectively, in DmpR. Particularly size of the pocket and alterations in the binding site residues seem to be the key determinants of the ligand specificity of XylR and related bacterial enhancers (Patil *et al.*, 2016; Ray *et al.*, 2016, 2018). To verify accuracy of the modelled binding pocket, three cognate effectors (*m*-xylene, 3-methylbenzoate, toluene) and three aromatic molecules that do not activate wild-type XylR (phenol, 2,4-dinitrotoluene, biphenyl) were docked in the cavity (**Supplementary Fig. S3**) and sorted based on ΔG of their top-ranked orientation (Abril *et al.*, 1989; Galvão *et al.*, 2007). All three XylR activators showed lower binding energies (−7.2, −6.9, and −6.5 kcal/mol for *m*-xylene, 3-methylbenzoate, and toluene, respectively) than phenol, 2,4-dinitrotoluene, and biphenyl (−5.9, −4.6, and −2.2 kcal/mol, respectively).

**Figure 2.**
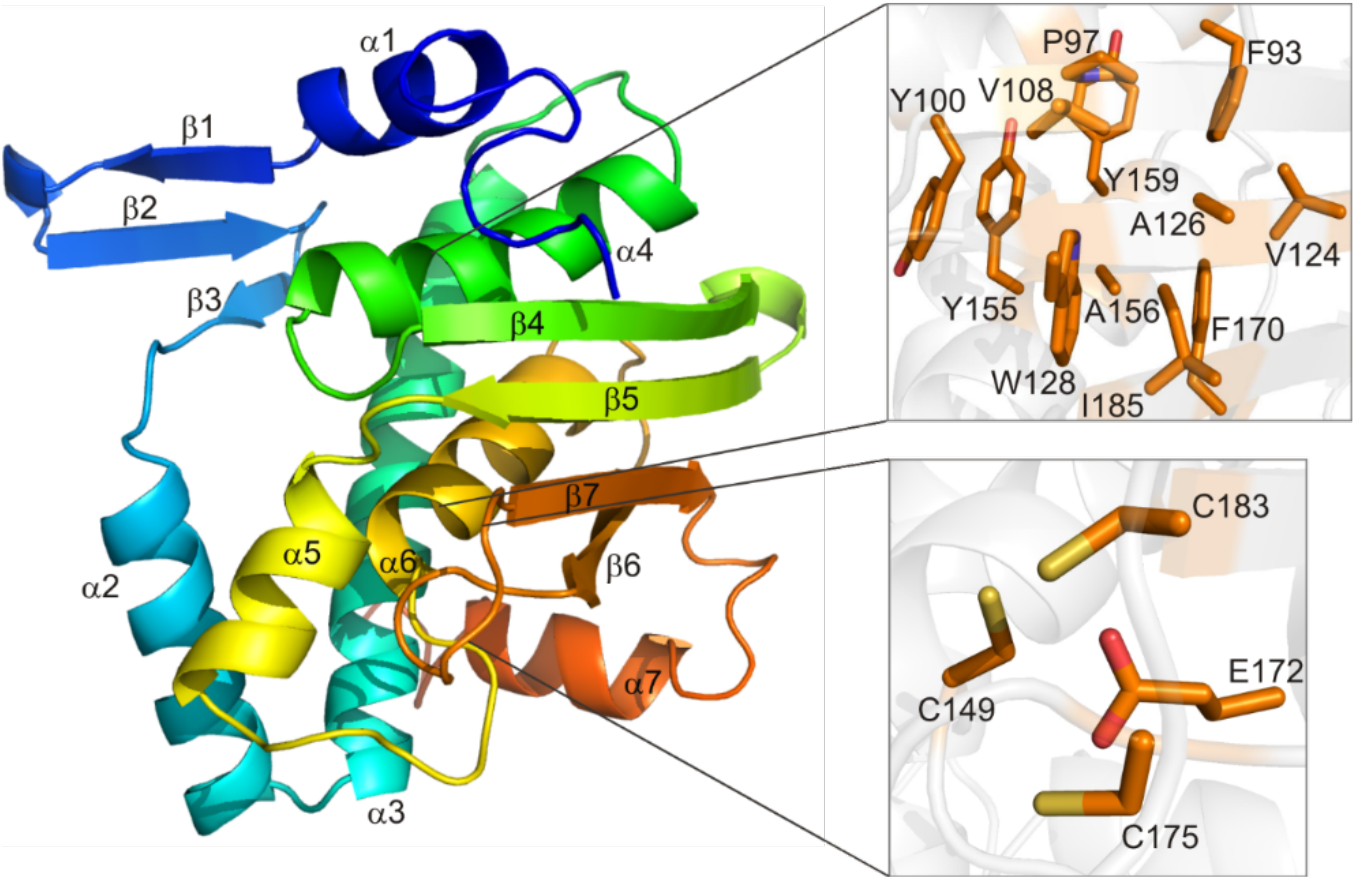
Tertiary structure of XylR effector binding domain A predicted by molecular threading with I-TASSER (Yang and Zhang, 2015) using 211 N-terminal amino acids of the transcription factor (Uniprot accession code: P06519). The structure backbone is coloured from N terminus (blue) to C terminus (orange). Secondary structure elements (seven βstrands and seven α helices) are labelled. Binding pocket residues (upper zoomed-in window) suggested by CAVER web 1.0 (Stourac *et al.*, 2019) and residues of a conserved zinc-binding site (lower zoomed-in window) identified previously in crystal structures of PoxR, MopR, and DmpR are highlighted as sticks. Only side chains of selected residues without hydrogens are visualised for better clarity (binding pocket residue G96 next to P97 is not shown). Dimerization interface (N motif) is supposedly formed by α helices α1 and α2, three-stranded antiparallel βsheet between them, and helix *α*5.

The inducer-recognition site subregion in the XylR A domain model is followed by a putative zinc-binding pocket between α6 and the second antiparallel hairpin motif (β6, β7). As was proven for PoxR, MopR, and DmpR, C149 from α6 and E172, C175, and C183 from the hairpin motif (XylR numbering is used) bind zinc atom which is important for structural integrity of the whole domain (Patil *et al.*, 2016; Ray *et al.*, 2016; Park *et al.*, 2020). The XylR E172K mutant reported by Delgado and Ramos (Delgado and Ramos, 1994) showed a substantially reduced response to cognate effector molecules such as toluene or *m*-xylene. This fact together with the highly conserved nature of the four residues among NtrC family members (**Supplementary Fig. S2**; Laitaoja *et al.*, 2013) sign the importance of the site and possible metal binding also in XylR. C-terminal part of the XylR A domain consists of helix α7 which transmits the signal upon effector binding to the B linker region.

### Prediction of inducer-access tunnels in the XylR A domain model

Because the predicted binding pocket is buried in the modelled A domain structure, a computational analysis was performed using CAVER web (Stourac *et al.*, 2019) to reveal possible entry site(s) and tunnel(s) that connect the cavity with the bulk solvent. Two hypothetical tunnels were predicted by the software – one going through the bundle of helices α3, α4, and α6 and the other one passing between β4 and α4 (**Fig. 3A**). Despite having a slightly narrower calculated bottleneck (0.7 vs. 0.9 Å), the latter tunnel seems to be the major passage for ligands. The main tunnel in PoxR sensory domain was proposed in a similar location and with bottleneck residues E96 and P112 (Patil *et al.*, 2016) which correspond to L94 and L110 suggested, together with F93 and M113, by CAVER for XylR (**Fig. 3B**). Moreover, the spatial shifts of the whole hairpin motif β 4 and β 5, including bottleneck residue P112, observed in PoxR sensory domain upon binding of ligands with different dimensions indicated certain flexibility of the binding pocket and its *mouth* between β4 and α4 (Patil *et al.*, 2016). This flexibility, possibly enhanced by the mobility of loop regions that interconnect secondary elements defining the binding pocket and its entry (**Fig. 3B**), might be another factor that shapes ligand specificity among related bacterial enhancers from the NtrC superfamily in diverse environmental conditions.

**Figure 3.**
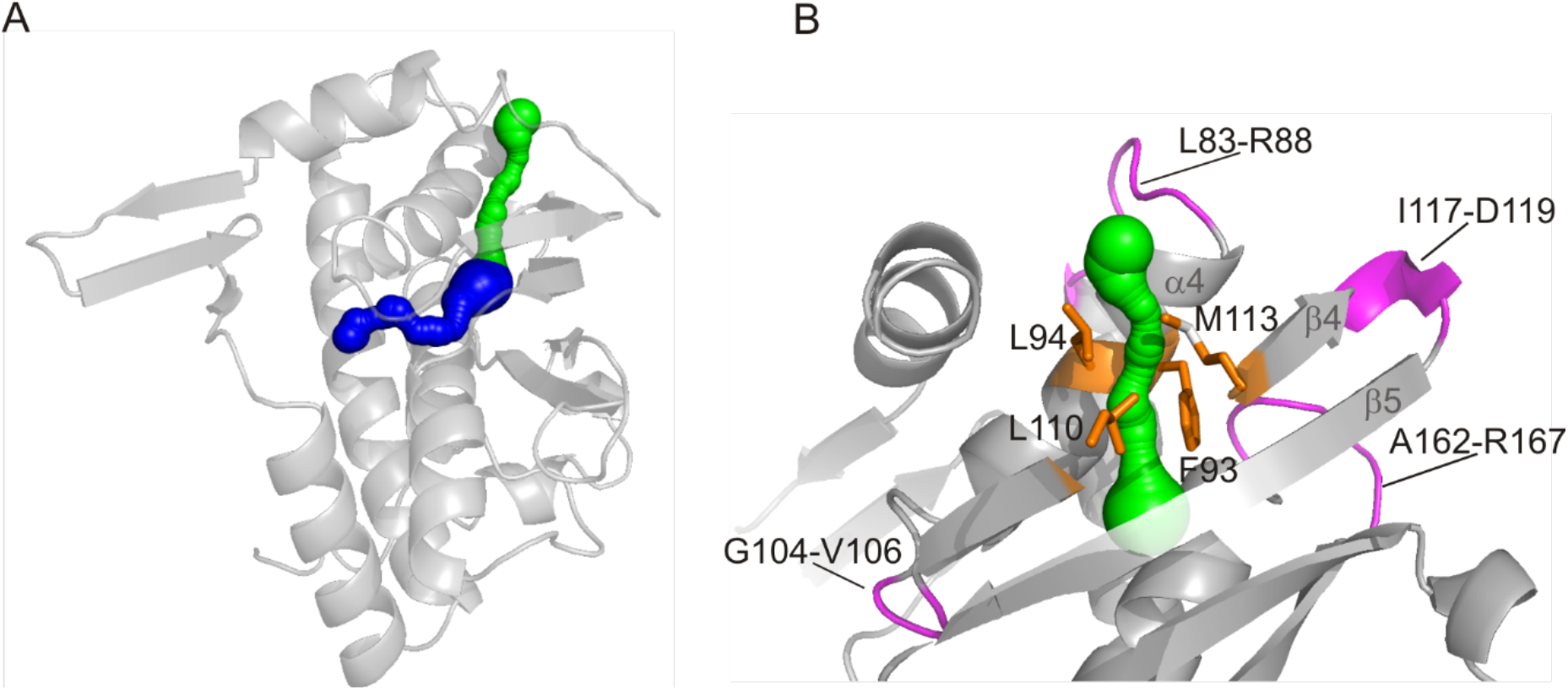
Protein tunnels for entry of effectors into the binding pocket of XylR identified by CAVER web 1.0 (Stourac *et al.*, 2019). (A) Two identified tunnels that connect XylR binding pocket with bulk solvent are shown in blue and green. (B) The probable main tunnel (in green) passing between helix α4 and strand β 4 is shown together with proposed bottleneck residues (orange sticks) and loop regions (in magenta) whose flexibility might affect the size of the tunnel and the binding pocket.

### Mapping and reinterpretation of mutations in XylR A domain

The new model allowed re-interpretation of mutations in the ligand recognition domain reported during the last three decades in studies focused on the alteration of XylR effector specificity (**Table 1, Fig. 4**). The specificity of XylR was modulated by targeted or random mutagenesis toward phenols and their derivatives (Garmendia *et al.*, 2001; Galvão *et al.*, 2007), precursors of explosive chemicals such as nitrotoluenes (Delgado and Ramos, 1994; Garmendia *et al.*, 2001; Galvão *et al.*, 2007; de Las Heras and de Lorenzo, 2011), or bulky effectors such as biphenyl (Garmendia *et al.*, 2001). As shown here by molecular docking, these compounds can be theoretically accommodated in the binding pocket of XylR A domain but their binding energies are higher than those of natural effectors (**Supplementary Fig. S3**). Hence, modification of the pocket’s shape and physico-chemical properties is a necessary prerequisite for the productive binding of these ligands.

**Table 1.** Reinterpretation of mutations in XylR A domain.

| Mutation | Former location and interpretation<sup>a</sup> | Updated location and interpretation | Effect | Reference |
| --- | --- | --- | --- | --- |
| F48I/T | Helix α3 close to the C terminus: Modulation of signal transfer from A to B and C domains of XylR | Helix α2: Dimer interface | Novel response to 2,4-dinitrotoluene (DNT), 3-NT, and chlorinated phenols; improved response to 2- and 3-NT | (Galvão *et al.*, 2007; de Las Heras and de Lorenzo, 2011) |
| F65L | Loop between β1 α4: Putative binding pocket | Helix α3: Dimer interface | Improved response to 2- and 4-NT and cognate effector 3-xylene; novel response to 3-NT | (Garmendia *et al.*, 2001) |
| P85S | Strand β2 | Loop between helices α3 and α4: Increased flexibility of the loop and binding pocket | Activation of transcription in absence of effector | (Delgado *et al.*, 1995) |
| P97A | Helix α5: Interdomain contacts | Helix α4: Binding pocket, near access tunnel | Novel response to 2,4-DNT | (Galvão *et al.*, 2007) |
| V124A | Helix α6: Interdomain contacts | Strand β5: Binding pocket | Stronger response to cognate effectors 3-xylene and benzene; improved response to NTs and biphenyls | (Garmendia *et al.*, 2001) |
| D135N | Strand β4: Interdomain contacts | Helix α5: Dimer interface | Activation of transcription in absence of effector; improved response to 2-NT and novel response to 3-NT | (Delgado *et al.*, 1995; Salto *et al.*, 1998) |
| I136T | Strand β4: Near putative binding pocket | Helix α5: Dimer interface | Improved response to new effector 2,4-DNT | (de Las Heras and de Lorenzo, 2011) |
| Y159F | Loop between α7 and β5 | Helix α6: Binding pocket, stabilisation of helices α6 and α4 forming bottom of binding pocket. | Novel response to 2,4-DNT and chlorinated phenols; improved response to NTs in combination with mutation in L222 | (Galvão *et al.*, 2007) |
| E172K | Loop between β5 and α8: Putative binding pocket | Zinc-binding site | Lower response to native effectors; novel response to 3-NT | (Delgado and Ramos, 1994) |
| S174R | Loop between β5 and α8: Putative binding pocket | C terminus of strand β6: Near zinc-binding site | Improved response to new effector 2,4-DNT | (de Las Heras and de Lorenzo, 2011) |
| L184I | Helix α8: Interdomain contacts | N terminus of strand β7: Near zinc-binding site | Improved response to 2- and 4-NT and benzene; novel response to 3-NT | (Garmendia *et al.*, 2001) |
| L222R | B linker transmitting signal from domain A do domain C. | B linker transmitting signal from domain A do domain C. | Novel response to 2,4-DNT | (Galvão *et al.*, 2007; de Las Heras and de<br>Lorenzo,<br>2011) |
| Shuffled<br>region 46-50* | Helix α3 | Helix α2: Dimer<br>interface | Improved response to NTs,<br>novel response to phenol<br>and biphenyl | (Garmendia<br>*et al.*, 2001) |
| Shuffled<br>region<br>161-166<sup>b</sup> | Strand β5 | Loop between α6<br>and β6: Flexibility of<br>the beta sheet<br>forming part of<br>binding pocket and<br>zinc-binding site;<br>signal transmission<br>to B linker | Improved response to NTs,<br>novel response to 3-NT,<br>phenol and biphenyl | (Garmendia<br>*et al.*, 2001) |

<sup>a</sup>Based on previous structural model published by Devos and co-workers (2002).
<sup>b</sup>This region was modified by DNA shuffling between A domains of XylR and DmpR.

**Figure 4.**
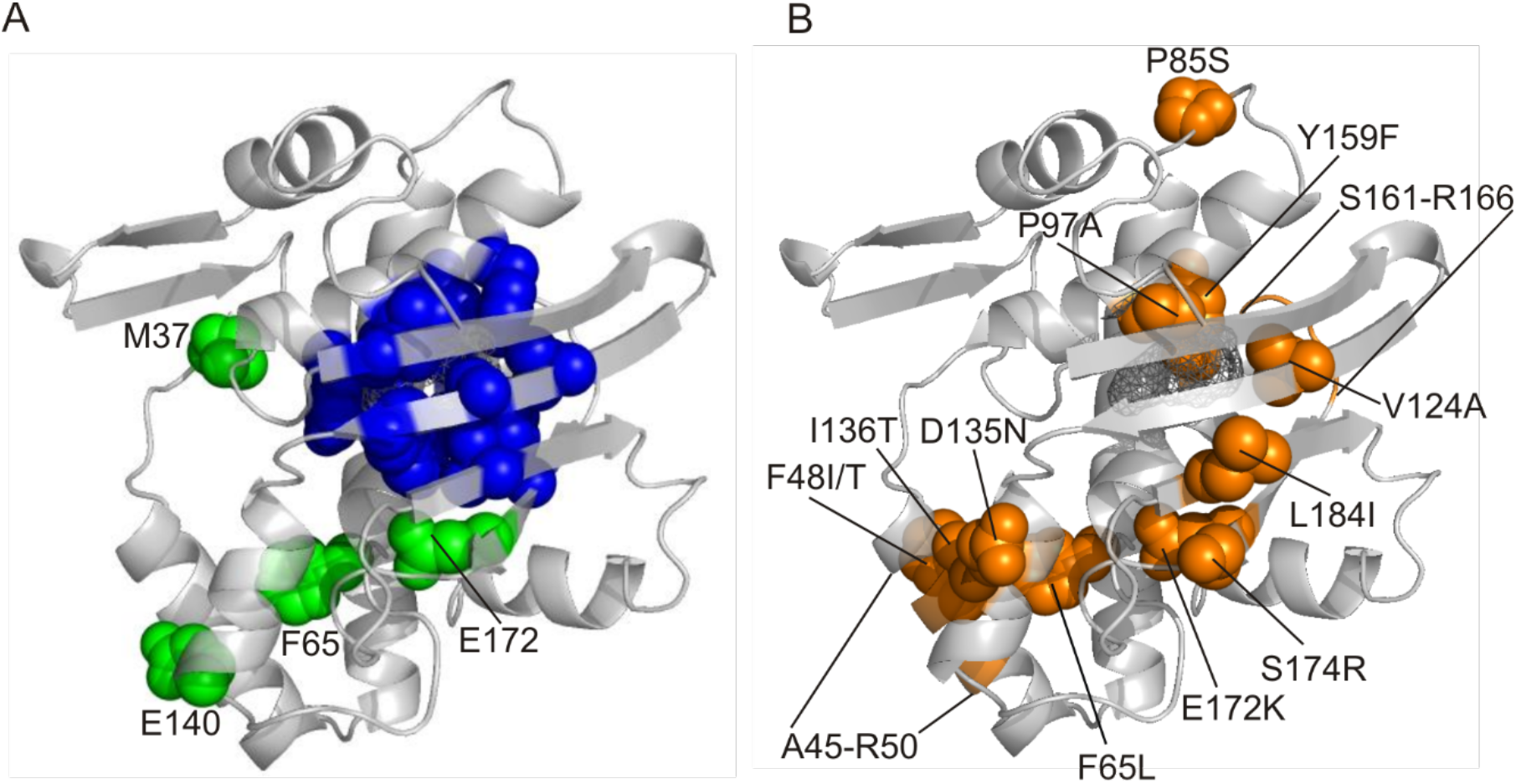
Mapping of revisited binding pocket residues and residues previously targeted by mutagenesis (listed in Table 1) on the predicted structure of XylR effector binding domain. (A) The residues proposed to interact with XylR effectors based on the structural model of Devos et al. (2002) are shown as green spheres and labelled. Side chains of the binding pocket residues proposed for the new model (F93, G96, P97, Y100, V108, V124, A126, W128, Y155, A156, Y159, F170, I185) are shown as blue spheres. (B) Single residues reported in mutagenesis studies are shown as orange spheres and labelled. Shuffled regions (A45-R50 and S161-R166) are shown as orange parts of the cartoon. The surface of the binding pocket is shown as a grey wireframe.

In the former model, Devos *et al.* (2002) proposed a binding groove to be formed by four loops, and particularly four conserved residues in this site (M37, F65, E140, and E172) were expected to contribute to the effector specificity of XylR (**Supplementary Fig. S1**). However, none of these amino acids makes part of the actual binding pocket nor are they in close proximity to it (**Fig. 4A**). M37, F65, and E140 are positioned in dimer interface (in β3, α3, and α5, respectively) while E172 is one of the residues forming putative zinc-binding site. Also, the majority of the reported mutations have different locations and structural effects than previously suggested (**Table 1, Figure 4B**). Only three amino acid substitutions (P97A, V124A, and Y159) occurred directly in the binding pocket. Interestingly, more mutations were located along with the A domain dimer interface (F48I/T, F65L, D135N, I136T, shuffled region 46-50). The frequent occurrence of mutations in this region in protein variants with altered inducer specificity was observed also for the other mentioned EBPs (Ray *et al.*, 2016). Several substitutions were present in the putative zinc-binding site (E172K) or in close proximity to it (S174R, L184I). These might affect the folding of the whole domain and secondarily also the flexibility of the binding cavity. In the case of the aforementioned substitution E172K, this effect was already too pronounced and detrimental for the productive binding of most of the tested ligands including natural effectors of XylR (**Table 1**; Delgado and Ramos, 1994). Last but not least, some reported mutations (P85S, shuffled region 161-166) can be found in the loops that connect secondary structure elements that shape the binding pocket and zinc-binding site. For instance, the substitution of rigid proline in position 85 for serine resulted in XylR variant capable of transcription activation in the absence of effector (Delgado *et al.*, 1995). This is not surprising from the current perspective, because a new A domain model locates the residue in the loop between helices α3 and α4 which form the bottom of the binding pocket and its entry tunnel. Hence the effect might be attributed to the loosening of these crucial parts of the protein. Also, mutation L222R in B linker, which is responsible for signal transduction between A and ATPase domain, is worth mentioning though it cannot be located in the current model. This substitution emerged repeatedly from error-prone PCR libraries together with mutations in A domain and gave rise to XylR variants responsive to the xenobiotic chemical 2,4-dinitrotoluene (**Table 1**; Galvão *et al.*, 2007).

Taken together, the hereby presented model of XylR A domain allows a realistic explanation of the effects of new and previously reported mutations. Despite the fact that the interpretations based on the old model of Devos et al. (2002) were mostly incorrect, the major conclusions of the former studies remain valid. Effector specificity of XylR and related transcription factors is a result of complex, sometimes counterintuitive, interactions of several factors (de Las Heras and de Lorenzo, 2011). It can be modulated by substitutions in the binding pocket (Ray *et al.*, 2018) as well as targeted or random mutations occurring in distant locations, particularly in A domain dimer interface, in certain loops, or in regions responsible for signal transduction.

### Rationale for experimental validation of the revised model

Next, we sought to verify the new model experimentally. One possible option was a chemical cross-linking with an effector molecule possessing a functional group that can form a covalent bond with certain amino acid moieties of the protein (Mattson *et al.*, 1993). Such a technique in combination with mass spectrometry can be used to confirm amino acids exposed to the solvent, including those in the ligand-binding pouch (Gingras *et al.*, 2007). Benzyl bromide with highly reactive bromomethyl substituent was selected as a suitable structural analogue of natural aromatic effectors such as *m*-xylene or toluene. Binding pocket in XylR A domain can theoretically accommodate benzyl bromide (**Supplementary Fig. S3G**), though the binding energy ΔG calculated during molecular docking of this ligand (−5.8 kcal/mol) was higher than that of *m*-xylene (−7.2 kcal/mol). Benzyl bromide acts as a selective alkylator of sulfur nucleophiles such as methionine or cysteine but can react also with other nucleophilic amino acids (including tyrosine, aspartic acid, or lysine present in the binding pocket or on the surface of XylR) when applied in a higher amount and for a longer time interval (Rogers *et al.*, 1976; Lang *et al.*, 2006). We hypothesized that interactions of XylR with benzyl bromide would provide information on site(s) in the protein structure which naturally interact(s) with aromatic effectors and amino acid residues exposed on the protein surface, and, thus, would either support or disprove the validity of the new model.

### In vivo evidence of interaction between XylR and benzyl bromide

Benzyl bromide interaction with XylR was initially studied using *E. coli* strain CC118 *Pu-lacZ* (**Supplementary Table S1**) bearing recombinant plasmid pCON916 with *xylR* gene under the control of its native *Pr* promoter (de Lorenzo *et al.*, 1991; Garmendia *et al.*, 2001). In this system, wild-type XylR activated by *m*-xylene induced expression of β-galactosidase whose activity was quantified (**Figs. 5A and 5B**; Miller, 1972). When benzyl bromide was used instead of *m*-xylene, no β-galactosidase activity (neither the background activity observed in the absence of effector) was detected (**Fig. 5C**). It is worth mentioning here that exposure to the vapours of *m*-xylene and benzyl bromide did not affect the viability of the bacterial strains used in this study. Hence, there were two possible explanations for the observed lack of β-galactosidase activity in cells exposed to benzyl bromide. Either benzyl bromide inactivated β-galactosidase or it interacted with XylR in a way that resulted in a protein form unable of *Pu-lacZ* induction. To test the first hypothesis, β-galactosidase activity was measured in strain *E. coli* MC4100 [MAD2] which expressed the mutant *xylRΔA* gene which encodes a constitutive variant of the transcription factor lacking the signal recognition domain (**Supplementary Table S1**). As expected, XylRΔA activated the *Pu-lacZ* fusion of *E. coli* MC4100 [MAD2] both in the presence and absence of the ligand. β-galactosidase activity was detectable in this strain even after the addition of benzyl bromide (**Fig. 5C**) therefore indicated that benzyl bromide did not affect the enzyme but rather interacted with XylR, most probably with its A domain. These results showed that the wild-type XylR is activated when it comes in contact with *m*-xylene and it loses its capacity to induce transcription upon interaction with benzyl bromide.

**Figure 5.**
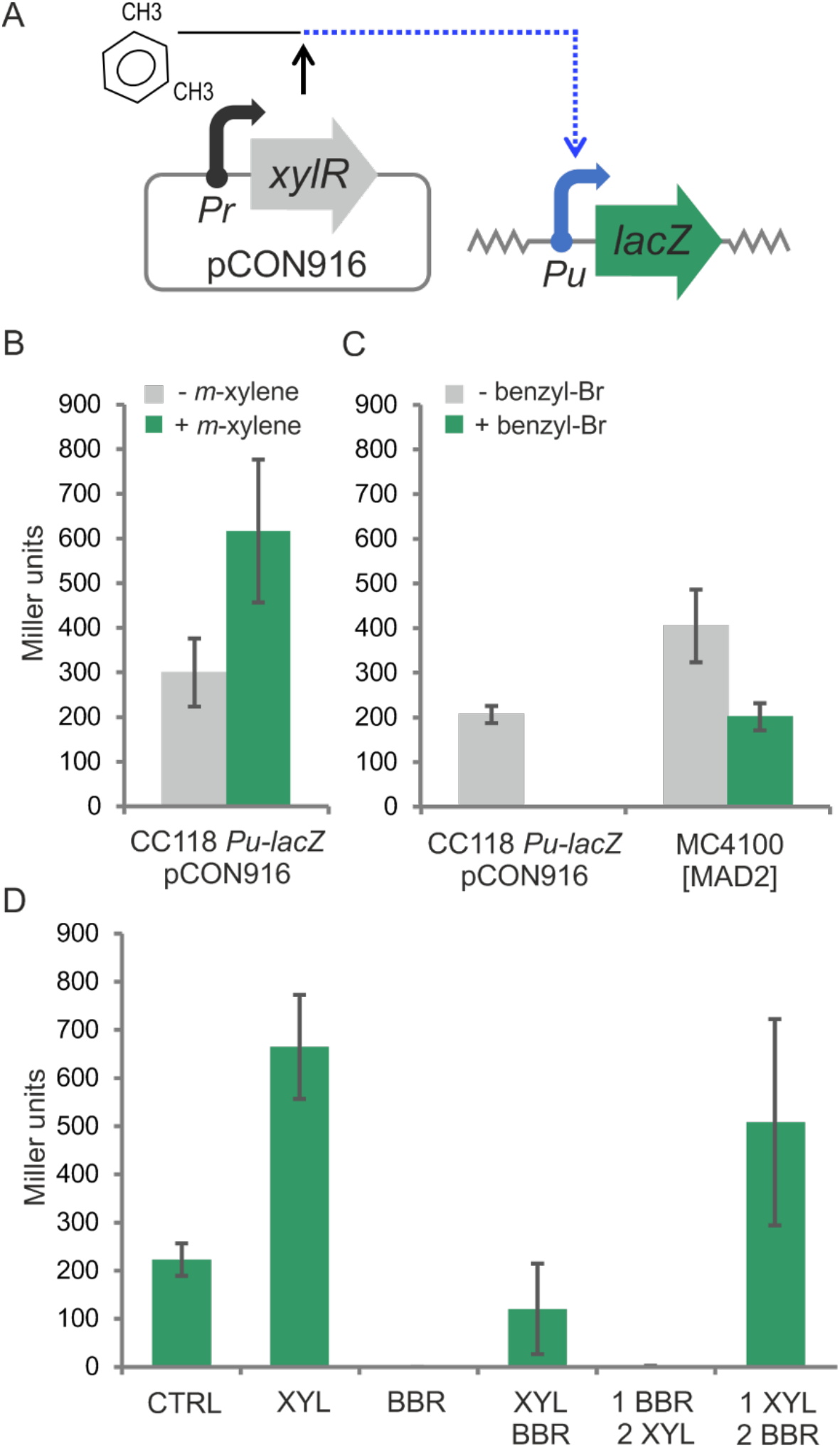
*In vivo* evidence of XylR interaction with *m*-xylene and benzyl bromide. A) A scheme of the sensor device in *Escherichia coli* CC118 *Pu-lacZ* strain with pCON916 plasmid bearing wild-type *xylR*. B) β-galactosidase activity of *E. coli* CC118 *Pu-lacZ* pCON916 in the absence or presence of *m*-xylene effector. C) β-galactosidase activity of *E. coli* strain CC118 *Pu-lacZ* pCON916 and strain MC4100 [MAD2], bearing *xylR* variant with deleted A domain, in absence or presence of benzyl bromide. D) β-galactosidase activity of *E. coli* CC118 *Pu-lacZ* pCON916 strain exposed to *m*-xylene and/or benzyl bromide in several specific conditions. Cells were grown to OD_600_ of 1.0 and then were: left uninduced (CTRL), exposed to saturated vapours of *m*-xylene (XYL), benzyl bromide (BBR), or of both chemicals in parallel (XYL/BBR), first exposed to benzyl bromide and then to *m*-xylene (1 BBR/2 XYL), or first exposed to *m*-xylene and then to benzyl bromide (1 XYL/2 BBR). Shown data represent means ± SD from two to three experiments.

Further tests were conducted again with *E. coli* CC118 *Pu-lacZ* pCON916. The strain was exposed to *m*-xylene and/or benzyl bromide in several different conditions to probe whether there was a competition between the two molecules for the same site in the structure of wild type XylR (**Fig. 5D**). Parallel exposure to *m*-xylene and benzyl bromide in a 1:1 ratio resulted in the significantly reduced β-galactosidase activity of the cells. Furthermore, XylR was completely unable to induce expression of *lacZ* when the cells were first exposed to benzyl bromide and then to *m*-xylene. However, the induction capacity of XylR was almost fully retained when the order of the two chemicals was reverse. These results suggested that benzyl bromide acted as an inhibitor of XylR activation by *m*-xylene and that such inhibition was probably competitive (i.e., inhibitor and effector molecules competed for the same binding site). To test this hypothesis, we turned to the biosensor strain *P. putida* BXPu-LUX, which bear chromosomal insertions of DNA encoding [i] the *xylR* gene expressed from its native promoter *Pr* and [ii] a *Pu-lux* transcriptional fusion (**Fig. 6A**; de Las Heras *et al*., 2008). In this strain, XylR activated by an effector induces expression of the *luxCDABE* operon from *Photorhabdus luminescens* and the specific bioluminescence of the whole cells can be detected non-disruptively in selected time intervals. On this background, *P. putida* BXPu-LUX cells were exposed to four different vapour pressures of *m*-xylene in the absence (**Fig. 6B**) or presence (**Fig. 6C**) of benzyl bromide and bioluminescence was recorded at fixed times. The inverse initial velocities of bioluminescence formation were plotted against the inverse vapour pressures in the double reciprocal plot (**Fig. 6D**). In the resulting graphics, the two dashed lines had the same Y-intercept, implying that benzyl bromide is a competitive inhibitor of XylR, i.e., it can enter and occupy non-productively the same binding site in the A domain.

**Figure 6.**
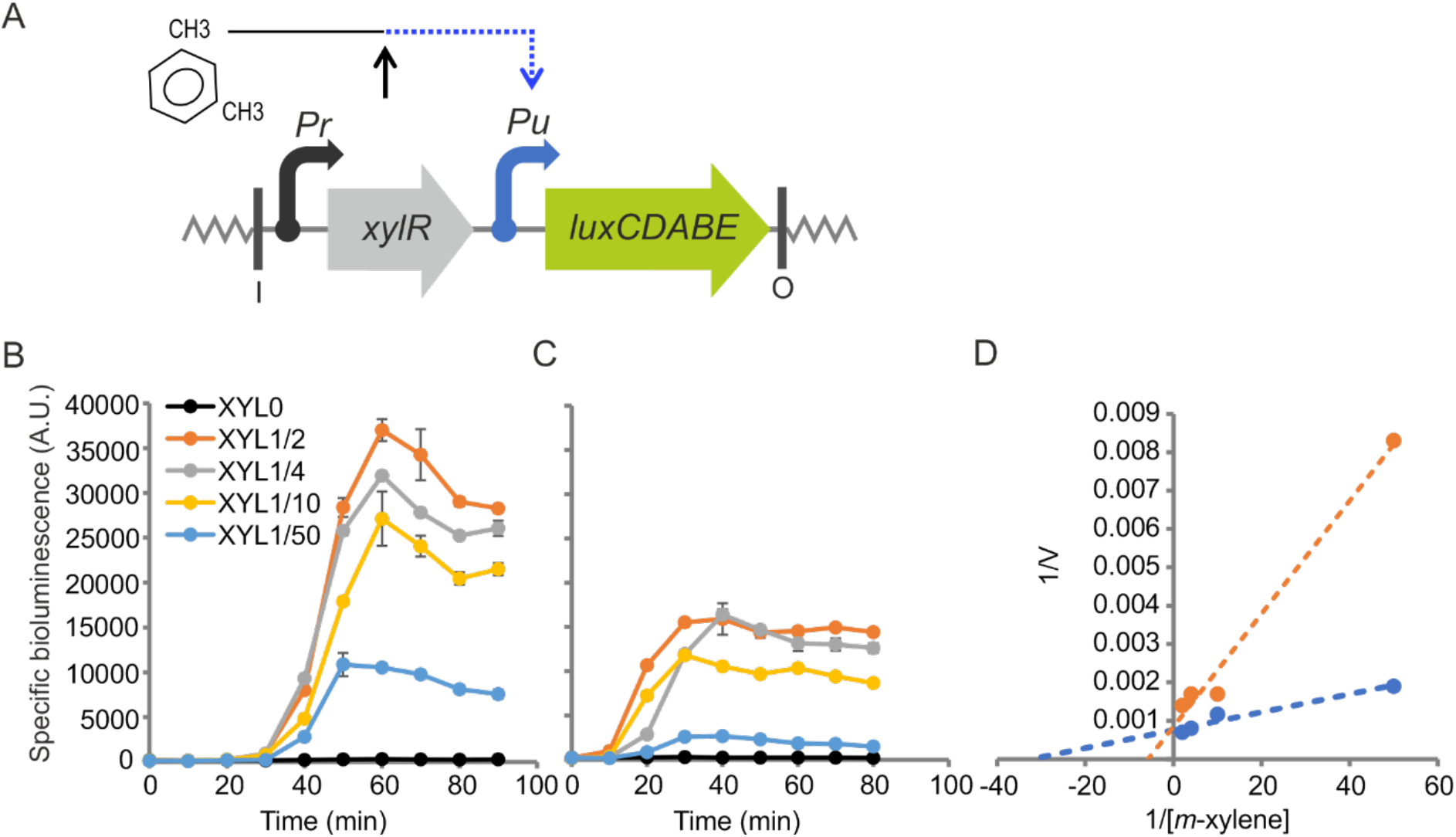
Determination of the competitive relation between *m*-xylene and benzyl bromide in XylR activation *in vivo*. A) A scheme of the sensor device in *Pseudomonas putida* BXPu-LUX strain (de Las Heras *et al*., 2008). B) Detection of specific bioluminescence of *P. putida* BXPu-LUX exposed to four different vapour pressures of *m*-xylene (XYL). The fractions shown in the figure legend correspond to the portions of the effector mixed with dibutyl phthalate solvent in 50 μl of the total volume of the mixture used for the induction of the cells. C) Detection of specific bioluminescence of *P. putida* BXPu-LUX exposed to four vapour pressures of *m*-xylene (identical to those used in the previous experiment) and constant vapour pressure of benzyl bromide. Benzyl bromide was mixed with dibutyl phthalate and *m*-xylene to form 1/20 of the total volume (50 μl) of the mixture. D) A double reciprocal plot for the four vapour pressures of *m*-xylene without (blue dots) and with (orange dots) a constant vapour pressure of benzyl bromide plotted against the velocity of production of specific bioluminescence of the strain BXPu-LUX. The velocities were calculated for time intervals 30 – 50 min in Figure 5A and 10 – 30 in Figure 5B. Shown data represent means ± SD from three biological replicates.

### Chemical cross-linking of purified XylR-His with benzyl bromide

The data above most plausibly indicated that benzyl bromide reaches out and interacts with the same target site in the A domain of XylR, but it is unable to activate the protein—instead, it blocks the ability of *m*-xylene to act as a bona fide inducer. Given the chemical reactivity of benzyl bromide, we entertained that the treatment of purified XylR with this reagent could form covalent bonds with nucleophilic amino acids exposed on the surface of the transcription factor and those present in the accessible effector binding cavity of the A domain. Such bound amino acids could then be determined with mass spectrometry and mapped on the predicted protein structure—thereby validating or challenging the model. To this end, we generated a recombinant XylR variant with 6xHis tag on C terminus of the protein (see Experimental procedures for details). To verify its functionality, the DNA sequence of the His-tagged protein was first expressed from pCON1238 (**Supplementary Table S1**) in *E. coli* CC118 *Pu-lacZ* and β-galactosidase activity of the cells was measured after induction with *m*-xylene and compared with *E. coli* CC118 *Pu-lacZ* (pCON916) encoding the wild type *xylR* gene (**Supplementary Fig. S4A and S4B**). *E. coli* CC118 *Pu-lacZ* (pCON1238) cells were also exposed to *m*-xylene and/or benzyl bromide in the same conditions described before for *E. coli* CC118 *Pu-lacZ* (pCON916) as shown in **Supplementary Fig. S4C, Fig. 5D**. Although the capacity of XylR-His to activate *Pu-lacZ* fusion was affected by the presence of the tag, the interaction pattern with *m*-xylene and benzyl bromide was identical to that observed for wild-type XylR.

Next, XylR-His was produced in soluble form in *Escherichia coli* BL21(DE3) pLysS transformed with pCON1238 plasmid and purified (>90 % purity) using immobilized metal affinity chromatography (**Supplementary Fig. S5, Experimental procedures**). XylR was then mixed with the excess of benzyl bromide and the mixture incubated overnight in moderately alkali conditions to promote cross-linking with nucleophilic amino acids. The mixture was then loaded on SDS-PAGE gel and the XylR-His band was cut out for mass spectrometry analysis. XylR-His non-exposed to benzyl bromide was used as a control. Peptides with mass increments corresponding to benzylation were identified (**Supplementary information file 2**) and their amino acid sequence was determined. There were in total 16 modified amino acids in 15 different peptides (**Table 2**). These amino acids were distributed along the whole XylR sequence but, interestingly, eight of them (M14, W31, R60, M75, R81, R118, C175, S205) were concentrated in A domain which represents only about one third of the whole protein size (**Fig. 7A**). The side chains of all amino acids that reacted with benzyl bromide (3x Met, 3x Arg, 3x Trp, 2x Cys, 1x Ser, 1x Tyr, 1x Asp, 1x Lys, and 1x His) are potential nucleophiles in the deprotonated state with the following relative order of nucleophilicity of functional groups: R-S-> R-NH2 > R-COO-= R-O-(Rogers *et al.*, 1976; Lang *et al.*, 2006; Bischoff and Schlüter, 2012). Heteroaromatic systems of Trp and His are less prone to benzylation but such reaction is possible due to the nature of *pi* electrons of indole and imidazole rings, respectively, and was probably promoted by the longer reaction times of our experiment.

**Table 2.**
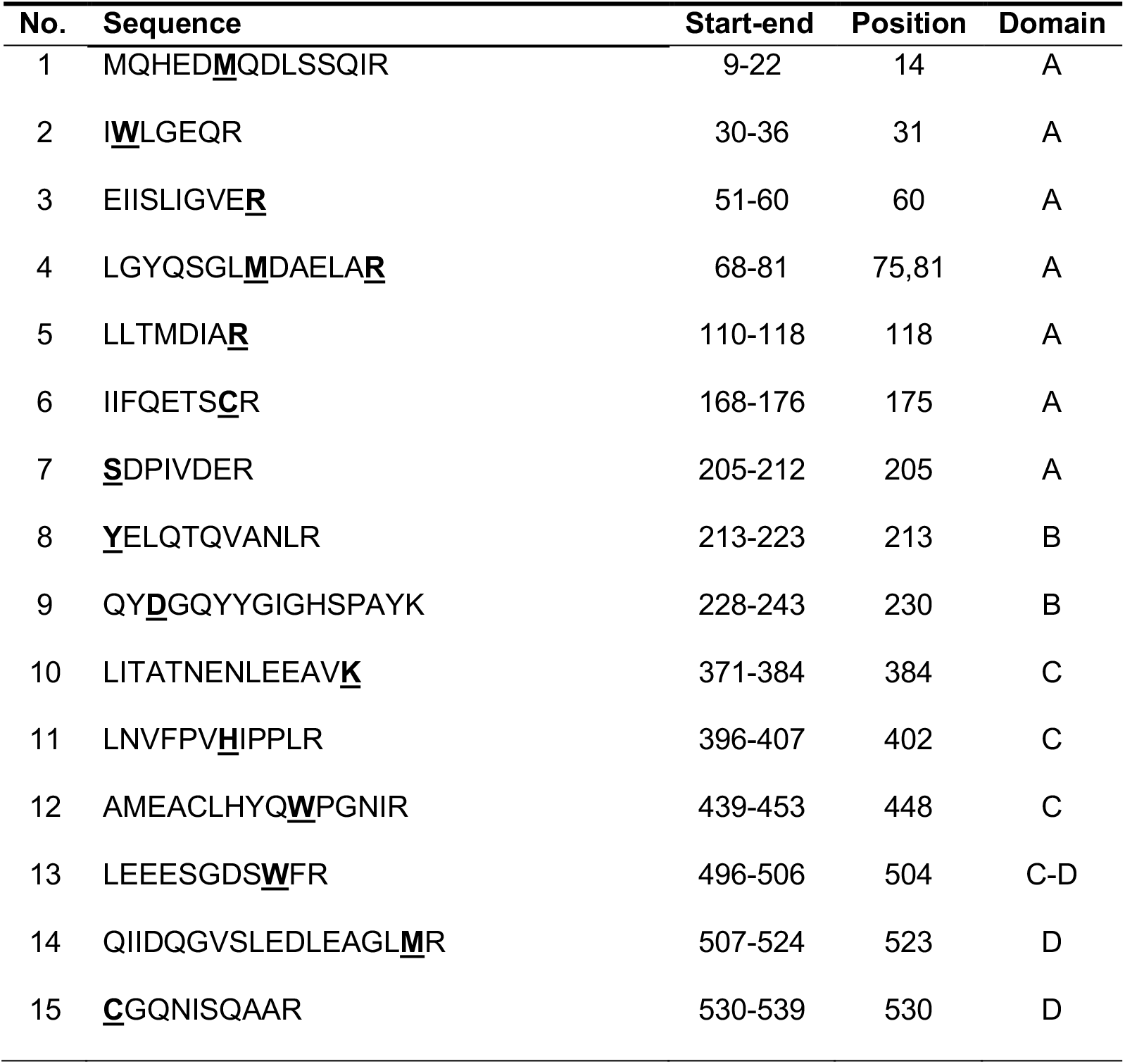
Peptides with amino acids cross-linked with benzyl bromide (highlighted) and their respective positions in the XylR-His protein.

**Figure 7.**
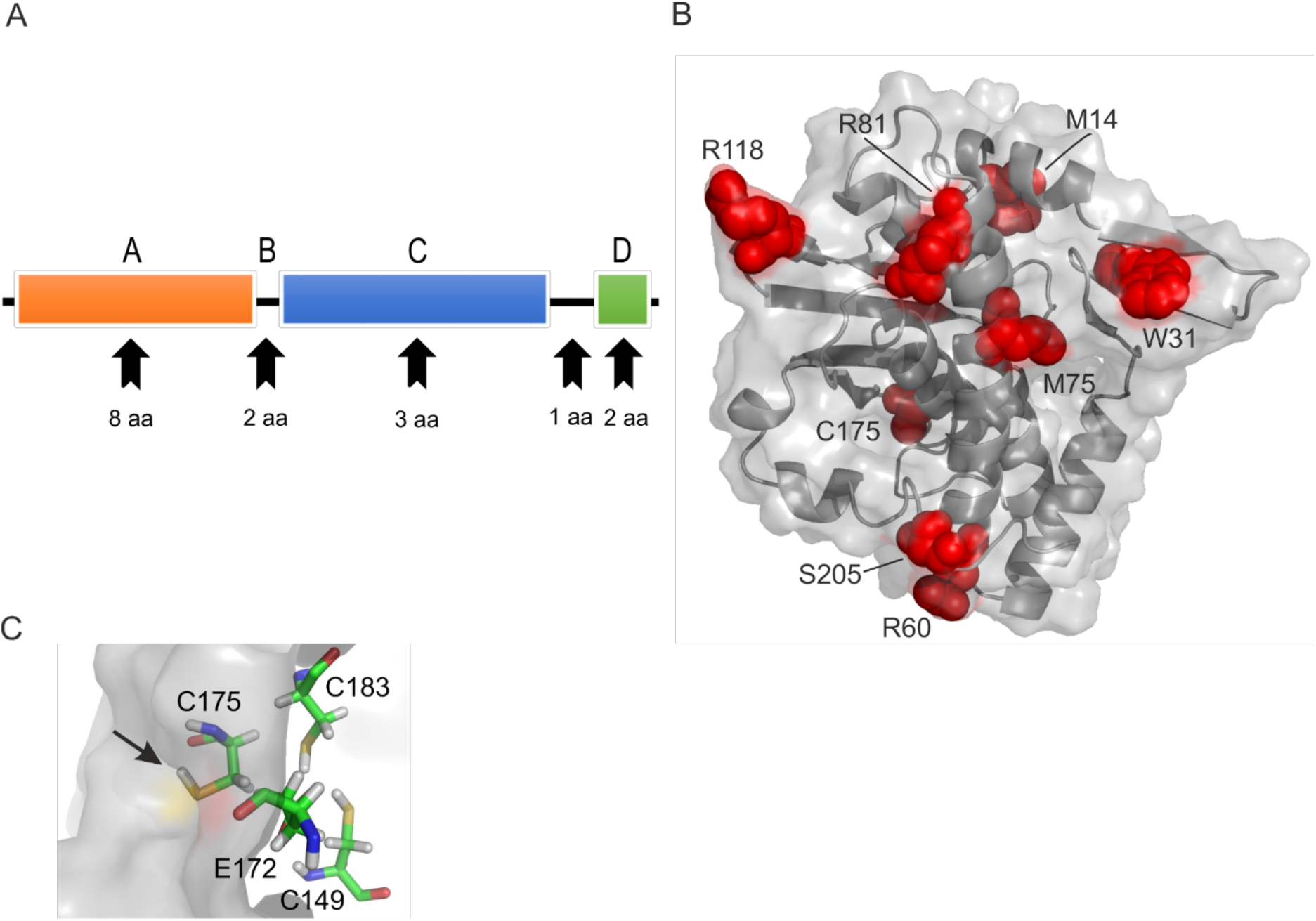
Mapping of the amino acid residues cross-linked with benzyl bromide in the XylR structure. (A) Modular scheme of XylR structure with all its domains and distribution of amino acids modified by benzyl bromide. The visualised XylR domains are: A domain which represents 37 % of the whole XylR sequence, B linker (4 %), C domain (ATPase domain, 42 %), and D domain (DNA binding domain, 8 %). (B) Mapping of the amino acid residues cross-linked with benzyl bromide in the predicted structure of XylR A domain. Modified residues are shown as red spheres in the XylR A domain structure depicted in cartoon and surface mode. (C) Detail of putative zinc-binding site and thiol group of C175 (black arrow) in the A domain model. Zinc-binding residues are shown as sticks coloured by element (carbon is green, nitrogen is blue, hydrogen is white, oxygen is red, sulphur is yellow), protein surface is depicted in grey.

Mapping of the thereby modified residues on the structural model of the XylR A was done manually in PyMOL and verified using NetSurfP 2.0 server (Klausen *et al.*, 2019). As shown in **Fig. 7B** and **Supplementary Fig. S6**, the chains of seven of these amino acids (M14, W31, R60, M75, R81, R118, S205), were exposed on the protein surface. The one exception was C175 that appeared *buried* in the protein structure. Note, however, that the nucleophilic thiol group of C175 is turned towards the protein surface (**Fig. 7C**) and it is the most accessible residue to benzylation out of the three cysteine residues that form a putative zinc-binding site in this part of the signal recognition domain. It was remarkable, that none of the amino acids predicted to shape the effector binding pocket or the tunnel (**Figs. 2 and 3**) was crosslinked with benzyl bromide, although such a reaction could in principle be possible with residues Y100, M113, W128, Y155, and Y159. Taken together, these results suggest that benzyl bromide did react with amino acids accessible on the A domain surface but, under given conditions, could not penetrate the protein structure to reach the binding pocket. As discussed below, we believe this is significant and, in fact, hints towards a possible distinct mechanism of activation of XylR.

## DISCUSSION

The data above calls for leaving behind the structural model of the XylR A domain proposed by Devos *et al.* (2002) and adopting an updated model prepared by the molecular threading of the cognate sequence within the available crystal structures of the A domains of related transcription regulators PoxR, MopR, and DmpR. That the new model is way more reliable than the previous one is accredited by the results of treating the purified protein with benzyl bromide. This reagent is known to covalently bind highly nucleophilic methionines (Rogers *et al.*, 1976; Lang *et al.*, 2006), but due to the relatively long time of treatment, it can also react with less nucleophilic amino acids, including surface-exposed tryptophans or histidines. These experiments, which define precisely the interior and the exterior of the domain, delivered a virtually perfect match between the model and the results (**Fig. 7**).

The new, dependable structure of the XylR A domain, in particular the organization of the effector-binding site, has enabled us to apply a wealth of structural analysis tools for the reinterpretation of abundant genetic data on this TF and it has provided hints for decoding its possible activation mechanism. Software for the detection of effector-binding cavities in proteins (I-TASSER and Caver) was specially useful in this respect. For instance, eight out of the thirteen residues forming the predicted binding pocket overlapped the so called *minimal binding region* (amino acids 110-186) previously defined by mutagenesis of XylR (Pérez-Martín and de Lorenzo, 1996). Furthermore, when the modelled binding cavity was probed by molecular docking, the binding energies of the bound ligands matched well the actual effector preferences of the wild-type XylR (Abril *et al.*, 1989; Galvão *et al.*, 2007). Most of the residues in the predicted binding cavity are conserved among PoxR, MopR, DmpR, and XylR. However, two residues in XylR pocket (Y100, and A126) differ from phenol binders PoxR, MopR, and DmpR. Replacement of His102 (Y100 in XylR) for Tyr in PoxR made the TF insensitive to phenol (Patil *et al.*, 2016). Single substitution H106Y (corresponds to Y100 in XylR) changed the specificity of MopR from phenols toward benzene ligands (Ray *et al.*, 2018). Replacement of an additional two residues F132A (A126 in XylR) and Y176F (F170 in XylR) further enlarged the MopR cavity for bulkier benzene derivatives. These analyses provide a good picture of how the architecture of the aromatic binding pocket looks like and they pave the way for developing XylR-based biosensors with rationally re-designed effector specificities. Yet, note that effector specificity of XylR and TFs might be determined not only by the geometry of the cavity that accommodates the aromatic compound (**Table 1**) but also by other structural and mechanistic constraints (de las Heras et al., 2011). In fact, it has been observed that effector mutants can very often arise from changes in locations distant from the binding pocket and the predicted tunnel. In this respect, the updated model suggests the presence in the XylR A domain of a distinct segment – so called N motif (**Fig. 2**) - involved in dimerization of the equivalent section of PoxR, MopR, and DmpR (Bush and Dixon, 2012; Patil *et al.*, 2016; Ray *et al.*, 2016; Park *et al.*, 2020). In this segment, the majority of effector mutants of DmpR, MopR, and XylR were mapped (**Table 1;**(Patil *et al.*, 2016; Ray *et al.*, 2016). How can such dimerization interface determine agonist specificity? Perhaps such mutants affect intramolecular signal transmission downstream of the effector binding proper. In these cases, the mutation away from the binding site may upgrade a good binder but non-productive aromatic into an efficient inducer. Obviously, how this can happen deserves further studies.

Still, the most intriguing aspect of the A domain geometry is the fact that the ultimate binding site of the protein for the aromatic effector is well buried in the internal core of the protein structure and not readily accessible from the outside. The only apparent way to reach out such a site is by penetrating a narrow tunnel (0.7 Å at the narrower part) coated with apolar amino acids that can hardly sustain the passage of molecules with a radius of ~7 Å like *m*-xylene (Li *et al.*, 2020). We were puzzled that despite our finding that benzyl bromide can act as a competitive inhibitor of xylene-activated XylR *in vivo* (in both *E. coli* and *P. putida*, **Figs. 5D and 6**), no benzyl bromide crosslink occurred *in vitro* with any of the potentially nucleophilic amino acids of the access tunnel and binding pocket (candidates were Y100, M113, W128, Y155, or Y159, **Figs. 2 and 3**) of the purified A domain. In contrast, binding pockets of purified PoxR, MopR, and DmpR can accept aromatic ligands (Patil *et al.*, 2016; Ray *et al.*, 2016; Park *et al.*, 2020). Because MopR sensory domain could be crystallized only in the presence of a ligand, Ray and co-workers (2016) have argued that it is quite flexible and exists in the open and closed forms. The switch from the open to the crystallizable compact form of A domain is proposed to be prompted by the binding of a ligand and zinc atom. Very recently, Park and co-workers (2020) reported that the phenol-binding pocket in the resolved DmpR protomer structures (which showed stronger electron density and supposedly higher occupation by phenol) had a smaller volume than the pocket in protomers with weaker electron density. This result also implies the existence of open and compact forms of A domains of certain EBPs. Our observations suggest a possible distinct yet complementary behaviour of XylR. Chances are that the A domain possesses an open ligand-accepting form *in vivo* but it cannot bind effectors under *in vitro* conditions when any potential port of entry to the binding cavity seems to be blocked. It could well be that the inducer interacts with the protein while it is being produced and folded, not after it has matured. In this way, the constellation of contacts could occur while the A domain is still a partially structured domain and thus accessible to the aromatic agonist. While such a folding-upon-binding scenario needs to be verified with additional experiments, it would explain the benzyl bromide data above as well as the recurrent failure to have a full-length XylR protein transcriptionally responsive to *m*-xylene in any *in vitro* test done thus far—while maintaining its full DNA-binding ability (Pérez-Martin *et al.*, 1997). Note that there may not be a clear-cut boundary between direct effector binding to the complete protein, binding a flexible form, or interacting during folding and it is plausible that different EBPs have followed different evolutionary itineraries to the same end.

## EXPERIMENTAL PROCEDURES

### Strains, plasmids, and growth conditions

Bacterial strains and plasmids used in this study are listed in Table S1. Nutrient-rich lysogeny broth (LB; (Sambrook *et al.*, 1989) and defined M9 minimal medium (Miller, 1972) with 2 mM MgSO4, 2 ‰ thiamine, and 0.2 % sodium citrate were used for growth of *E. coli* and *P. putida* strains. Bacteria were also grown in Petri dishes on LB agar (1.5 %) solid medium or M9 agar (1.6 %) medium with 2 mM MgSO4, 2 ‰ thiamine, and 0.2 % sodium citrate. *E. coli* and *P. putida* strains were grown at 37 °C and 30 °C, respectively, unless stated otherwise. If needed, liquid and solid media were supplemented with 100 or 500 μg/ml ampicillin (Ap, higher concentration was used for *P. putida*), 75 μg/ml kanamycin (Km), 30 μg/ml chloramphenicol (Cm), 10 μg/ml tetracycline (Tc), 1 mg/ml carbenicillin (Cb), or 10 μg/ml gentamicin (Gm) to select for bacteria with plasmid(s). The 5-bromo-4-chloro-3-indolyl-β-D-galactopyranoside (X-Gal; 40 μg/ml) was added to detect β-galactosidase activity. For the induction of liquid cultures with volatile compounds, 50 μl of *m*-xylene or benzyl bromide was dropped into the reservoir in the center of a specially designed culture flask. In the case of induction of cells grown on solid media, *m*-xylene (50 μl) was pipetted in a 200 μl plastic tip. The tip was fixed in the centre of the lid of a Petri dish with adhesive tape. Volatile reagents of superior purity (>99%) were purchased from Sigma Aldrich. Strains *E. coli* CC118 *Pu-lacZ* pCON916 and *E. coli* CC118 *Pu-lacZ* pCON1238 were freshly prepared by transforming chemocompetent *E. coli* cells (Sambrook *et al*., 1989) with respective plasmids.

### β-galactosidase and bioluminescence assays

For determination of β-galactosidase activity levels, *E. coli* strains CC118 *Pu-lacZ* pCON916, CC118 *Pu-lacZ* pCON1238, or MC4100[MAD2] were grown overnight in LB medium, in the morning diluted to OD_600_ of 0.05 in fresh LB and cultured till OD_600_ of 1.0. At this point, cells were exposed to saturated vapours of *m*-xylene (50 μl), benzyl bromide (50 μl), or 1:1 mixture of both chemicals (25 μl each) in conditions described in figure legends. After a 4 h interval, LacZ activity levels were determined in cells permeabilised by chloroform and sodium dodecyl sulphate using the method of Miller (1972). Alternatively, cells were first exposed to one of the two chemicals (25 μl) for 2 h and then the second compound was added (25 μl) for the remaining 2 h. Bioluminescence assays with biosensor strain *P. putida* BXPu-LUX were performed accordingly. After 4 h exposure to aromatic effector(s), culture aliquots (200 μl) were placed in 96-well plate (NUNC) and emission of light was measured with Victor II 1420 Multilabel Counter (PerkinElmer). Recorded bioluminescence was normalised by the optical density of cells in each well. A double reciprocal plot (**Fig. 6D**) was constructed from values of used *m*-xylene volumes and velocities of bioluminescence formation in the time interval between 30 and 50 min for assay without benzyl bromide (**Fig. 6B**) and between 10 and 30 min for assay with benzyl bromide (**Fig. 6C**). Exposure to the volatile chemicals had no negative effect on the viability of the cells.

### Purification of XylR-His

*E. coli* BL21(DE3) pLysS cells were transformed with pCON1238 plasmid bearing *xylR*-*His* gene, plated on LB agar plate with Ap and grown at 30 °C. Following day, cells from the plate were collected in liquid LB with Ap and grown at 30 °C (170 rpm) till reaching OD_600_ of 3.0. Cells were then diluted to OD_600_ of 0.15 in fresh LB with Ap and grown at 19 °C. Expression of *xylR-His* was induced with 0.4 mM IPTG at the OD_600_ of 0.7 and culture continued at 19 °C till reaching OD_600_ of 2.0. Cells were then centrifuged, re-suspended in buffer A (20 mM sodium phosphate, 1.0 M NaCl, 0.1 % Triton X-100, 10 % glycerol, 1 mM β-mercaptoethanol, pH 7.2) and frozen at −80 °C. Melted cells, kept on ice, were lysed by sonication and the crude extract was separated from cell ballast by centrifugation. The crude extract was filtered through a 0.22 μm membrane filter (Merck Millipore) and loaded on a disposable purification column packed with TALON Superflow Resin (Clontech). The resin was washed with buffer A with an increasing concentration of imidazole (0 – 200 mM). Samples taken from individual fractions were analysed on denaturing SDS polyacrylamide gels (8 %) stained with Coomassie Brilliant Blue (Bio-Rad). Fractions containing pure (>90 %) XylR-His protein (64.6 kDa) were pooled, concentrated in Amicon Ultra centrifugal filter (Merck Millipore), and dialysed against buffer B (20 mM sodium phosphate, 0.5 M NaCl, 0.1 % Triton X-100, 30 % glycerol, 1 mM β -mercaptoethanol, pH 7.5). Protein aliquots of concentration of 0.55 mg/ml were stored at −80 °C for further use.

### Exposure of purified XylR-His to benzyl bromide, mass spectrometry analysis of modified TF

Purified XylR-His (6.96 μM) in buffer B was mixed with benzyl bromide (74 mM) in a total volume of 20 μl in a microtube. The protein was left to react with the aromatic chemical overnight at room temperature with modest agitation. The whole volume of the reaction mixture was then loaded on denaturing SDS polyacrylamide gel (8 %) and the XylR-His band was cut for mass spectrometry analysis. The mass spectrometry analysis of the peptides obtained after trypsin digest of the protein sample was performed by Proteomics Core Facility of the Spanish National Center for Biotechnology (CNB-CSIC) in Madrid. Peptides were first separated by liquid chromatography using Ultimate 3000 nano LC system (Dionex) equipped with 75 mm I.D 100 mm reversed-phase column (300 nl/min flow) and then analysed using mass spectrometers 4800 MALDI TOF/TOF (Applied Biosystems) or HCT Ultra Ion-Trap (Bruker Daltonics) working in dynamic exclusion mode. These two sources of experimental evidence were complementary. In some cases, multiple reaction monitoring mode was used to isolate and fragment specific *m/z* values corresponding to putative benzyl bromide-labelled peptides. For protein identification, LC-ESI-MS/MS spectra were transferred to BioTools 2.0 interface (Bruker Daltonics) to search in the Uniprot database using a licensed version of Mascot v.2.2.04 search engine (Matrix Science). Search parameters were set as follows: carbamidomethyl cysteine as fixed modification by the treatment with iodoacetamide, oxidized methionines, and benzylation as variable modifications, peptide mass tolerance of 0.5 Da for the parental mass and fragment masses and 1 missed cleavage site. In all protein identifications, the probability mowse scores were greater than the minimum score fixed as significant with a *p*-value minor than 0.05. Peptides obtained by fragmentation of non-modified XylR-His were used as a control. Analysis of signal intensities allowed the identification of peptides with mass increments of +90 or +91 Da caused by benzylation of amino acids with nucleophilic functional groups.

### Multiple sequence alignment and prediction of XylR A domain structure by molecular threading

The amino acid sequences of transcriptional regulatory proteins XylR from *Pseudomonas putida* (UniProt ID: P06519), PoxR from *Ralstonia sp.* E2 (UniProt ID: O84957), MopR from *Acinetobacter guillouiae* (UniProt ID: Q43965), and DmpR (also known as CapR) from *P. putida* (UniProt ID: Q7WSM9) were retrieved in FASTA format and 211 amino acids of the respective ligand recognition domains were selected for multiple sequence alignment ClustalW (Chenna *et al.*, 2003) and prediction of secondary structure elements using and ESPript 3.0 (Robert and Gouet, 2014). The 211 amino acid sequence of XylR A domain was subsequently used for molecular threading by I-TASSER server (Iterative Threading ASSEmbly Refinement; Yang and Zhang, 2015) which represents a hierarchical approach for prediction of protein structure and function. I-TASSER has been repeatedly ranked as a top server in Community Wide Experiment on the Critical Assessment of Techniques for Protein 3D Structure Prediction (http://www.predictioncenter.org) and can be thus considered a reliable accurate tool for the given purpose. The structural model of XylR A domain was built based on multiple-threading alignments by LOMETS performed with templates from PDB and iterative TASSER assembly simulations. The model with the highest confidence (C) score was selected for further work.

### Prediction of binding pocket and tunnels in the model of XylR A domain

Ligand binding site in the structural model of XylR A domain was first predicted by COFACTOR and COACH approaches on I-TASSER server (Yang and Zhang, 2015). The set of the residues forming the binding pocket was deduced from the top-ranked PDB of the homologous phenol-responsive sensory domain of PoxR (PDB ID: 5FRU). The binding pocket in XylR A domain PDB file prepared in PyMOL 1.6.0.0 (Schrödinger) was then predicted also using CAVER web 1.0 (Stourac *et al.*, 2019). Pocket with the second highest druggability score in CAVER corresponded to the one previously proposed by I-TASSER. Pocket residues suggested by both I-TASSER and CAVER and manually verified in the modelled structure (F93, G96, P97, Y100, V108, V124, A126, W128, Y155, A156, Y159, F170, I185) are used in this work. Tunnels and bottleneck residues were calculated by CAVER using default program parameters (minimum probe radius 0.9 Å).

### Molecular docking of aromatic ligands in XylR A domain model

The modelled structure of XylR A domain was prepared for molecular docking in PyMOL. The structure with added hydrogens was saved as PDB file and uploaded together with a ligand molecule in MOL2 format. The geometries of ligands were optimised using Avogadro 1.2. The binding pocket region of XylR was selected for the docking performed by PyMOL Autodock Vina Plugin for Windows 2.2 (Trott and Olson, 2010). The binding pocket region was defined by a grid box of 40 × 40 × 40 Å. Ligand poses with the lowest binding energies were saved and visualized using PyMOL. The docking experiment was repeated three times for each ligand.

## ACKNOWLEDGEMENTS

Authors are indebted to the Severo Ochoa Programme of Centres of Excellence. This work was funded by the SETH (RTI2018-095584-B-C42) (MINECO/FEDER), SyCoLiM (ERA-COBIOTECH 2018 – PCI2019-111859-2) Project of the Spanish Ministry of Science and Innovation. MADONNA (H2020-FET-OPEN-RIA-2017-1-766975), BioRoboost (H2020-NMBP-BIO-CSA-2018-820699), SynBio4Flav (H2020-NMBP-TR-IND/H2020-NMBP-BIO-2018-814650) and MIX-UP (MIX-UP H2020-BIO-CN-2019-870294) Contracts of the European Union and the InGEMICS-CM (S2017/BMD-3691) Project of the Comunidad de Madrid - European Structural and Investment Funds - (FSE, FECER). The grant 19-06511Y of the Czech Science Foundation to P.D. is also gratefully acknowledged. The Authors declare that they have no conflict of interest.

## Supplementary Materials

**Supplementary information File 1:**

**Table S1** Strains and plasmids used in this study.

**Figure S1** Original structural model of XylR A domain proposed by Devos et al. (2002).

**Figure S2** Multiple sequence alignment of effector binding domain A of XylR, PoxR, MopR, and DmpR from *Pseudomonas putida* mt-2, *Ralstonia sp*. E2, *Acinetobacter guillouiae*, and *P. putida* KCTC 1452, respectively

**Figure S3** Docking of ligands in predicted binding pocket of XylR A domain.

**Figure S4** *In vivo* evidence of XylR-His interaction with *m*-xylene and benzyl bromide.

**Figure S5** Sodium dodecyl sulfate polyacrylamide gel electrophoresis (8 % gel) of purified XylR-His protein.

**Figure S6** Surface accessibility of amino acids in XylR A domain predicted by NetSurfP 2.0 server.

**Supplementary information File 2:**

The mass spectrometry analysis for identification of XylR peptides with nucleophilic amino acids cross-linked with benzyl bromide.

## Notes

### Competing Interest Statement

The authors have declared no competing interest.

