## Supplemental File 1 for "An updated structural model of the A domain of the *Pseudomonas putida* XylR regulator exposes a distinct interplay with aromatic effectors"

by

Pavel Dvořák[1], Carlos Alvarez-Carreño[2], Sergio Ciordia[3], Alberto Paradela[3] and Víctor de Lorenzo[3]

[1]*Department of Experimental Biology (Section of Microbiology), Faculty of Science, Masaryk University, Kamenice 753/5, 62500, Brno, Czech Republic.*

[2] *Centro Tecnológico José Lladó, División de Desarrollo de Tecnologías Propias, Técnicas Reunidas, Calle Sierra Nevada, 16, San Fernando de Henares, Madrid 28830, Spain*

[3]*Centro Nacional de Biotecnología-CSIC, Campus de Cantoblanco, Madrid 28049, Spain.*

* Correspondence to:

Víctor de Lorenzo

Centro Nacional de Biotecnología (CNB-CSIC)

C. Darwin 3, Campus de Cantoblanco, Madrid 28049, Spain

### Contents

**Supplementary tables** ……………………………………………………………………..3

**Table S1.** Strains and plasmids used in this study.

**Supplementary figures** …………………………………………………..………………..5

**Figure S1.** Original structural model of XylR A domain proposed by Devos *et al*. (2002).

**Figure S2.** Multiple sequence alignment of effector binding domain A of XylR, PoxR, MopR, and DmpR from *Pseudomonas putida* mt-2, *Ralstonia sp.* E2, *Acinetobacter guillouiae*, and *P. putida* KCTC 1452, respectively.

**Figure S3.** Docking of ligands in predicted binding pocket of XylR A domain.

**Figure S4.** *In vivo* evidence of XylR-His interaction with *m*-xylene and benzyl bromide.

**Figure S5.** Sodium dodecyl sulfate polyacrylamide gel electrophoresis (8 % gel) of purified XylR-His protein.

**Figure S6.** Surface accessibility of amino acids in XylR A domain predicted by NetSurfP 2.0 server.

**References** …………………………………………………………...…………………11

### Supplementary tables

**Supplementary Table S1.** Strains and plasmids used in this study.

| Strain or plasmid | Characteristics | Source or reference |
|---|---|---|
| ***Escherichia coli*** | | |
| BL21(DE3) pLysS | F- *ompT gal dcm lon hsdS*$_B$($r_B^-$ $m_B^-$) λ(DE3) with pLysS plasmid; $Tc^r$ | Agilent Technologies |
| CC118 *Pu-lacZ* | CC118 with mini-Tn5 chromosomal insertion of transcriptional fusion *Pu-lacZ*; $Sm^r$ | (de Lorenzo *et al.*, 1991) |
| MC4100[MAD2] | *araD319* Δ(*argF'lac*)*U169 rpsL150 relA1flbB5301 deoC1 ptsF25* with MAD2 cassette (*xylRΔA Pu-lacZ*); $Cm^r$ | Gift from Dr. Ildefonso Cases |
| ***Pseudomonas putida*** | | |
| BXPu-LUX | Biosensor strain for *m*-xylene: derivative of KT2440 with chromosomal insertion of *Pr-xylR-Pu-luxCDABE* | (de Las Heras *et al.*, 2008) |
| **plasmids** | | |
| pCON916 | Expression vector: *oriV*(RSF1010) *Pr-xylR*; $Cb^R$ | (Garmendia *et al.*, 2001) |
| pCON1238 | Expression vector: derivative of commercial pET3C (Novagen, Merck) with cloned *xylR-His*; $Ap^r$ | Gift from Dr. Victoria Shingler |

### Supplementary figures

**Figure S1.** Original structural model of XylR A domain proposed by Devos *et al*. (2002).

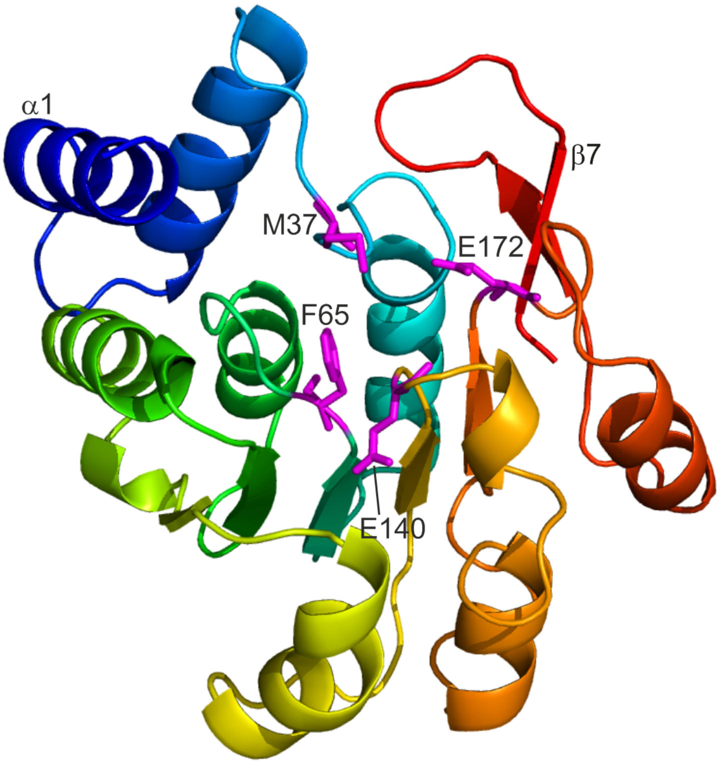


The structure backbone (207 amino acids) is coloured from N terminus (α1, blue) to C terminus (β7, red). The threading model was based on the crystallographic data of the catechol *o*-methyltransferase (COMT; PDB code 1vid), a typical α/β fold, consisting of eight α-helices and seven β-strands. Residues of the theoretical binding pocket (M37, F65, E140, and E172) are shown as magenta sticks.

**Figure S2.** Multiple sequence alignment of effector binding domain A of XylR, PoxR, MopR, and DmpR from *Pseudomonas putida* mt-2, *Ralstonia sp.* E2, *Acinetobacter guillouiae*, and *P. putida* KCTC 1452, respectively (UniProt ID: P06519, O84957, Q43965, Q7WSM9, respectively).

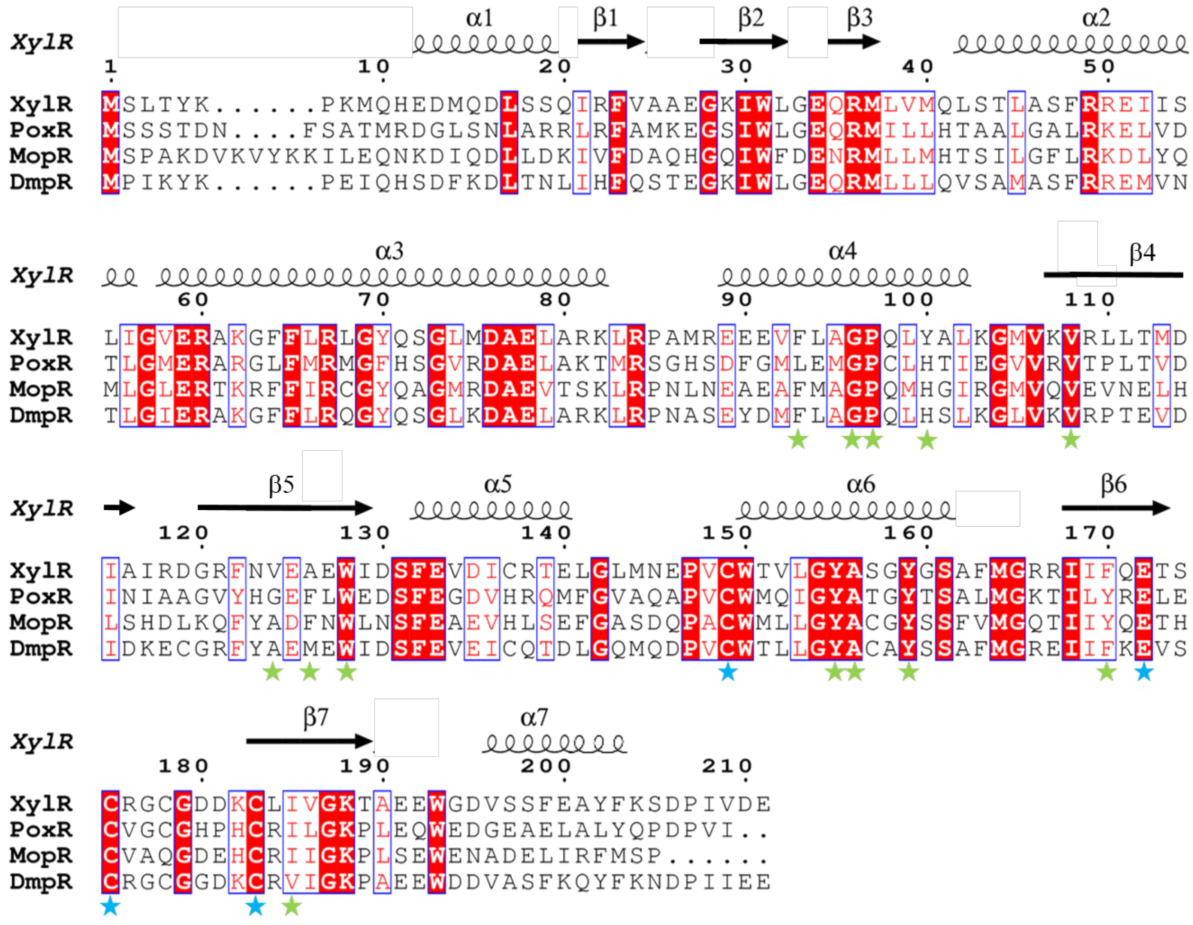

Secondary structure elements for XylR A domain are visualised at the top of the alignment. Conserved residues identified by ESPript 3.0 software (Robert and Gouet, 2014) are highlighted with red background, highly similar residues (global similarity score > 0.8) are in red and framed in blue. XylR binding pocket residues suggested by COFACTOR and COACH functions based on I-TASSER structure prediction (Yang *et al.*, 2015) and verified by CAVER web (Stourac et al., 2019) are marked with green stars. Residues of conserved zinc-binding site identified previously in crystal structures of PoxR, MopR, and DmpR are marked with blue stars.

**Figure S3.** Docking of ligands in predicted binding pocket of XylR A domain.

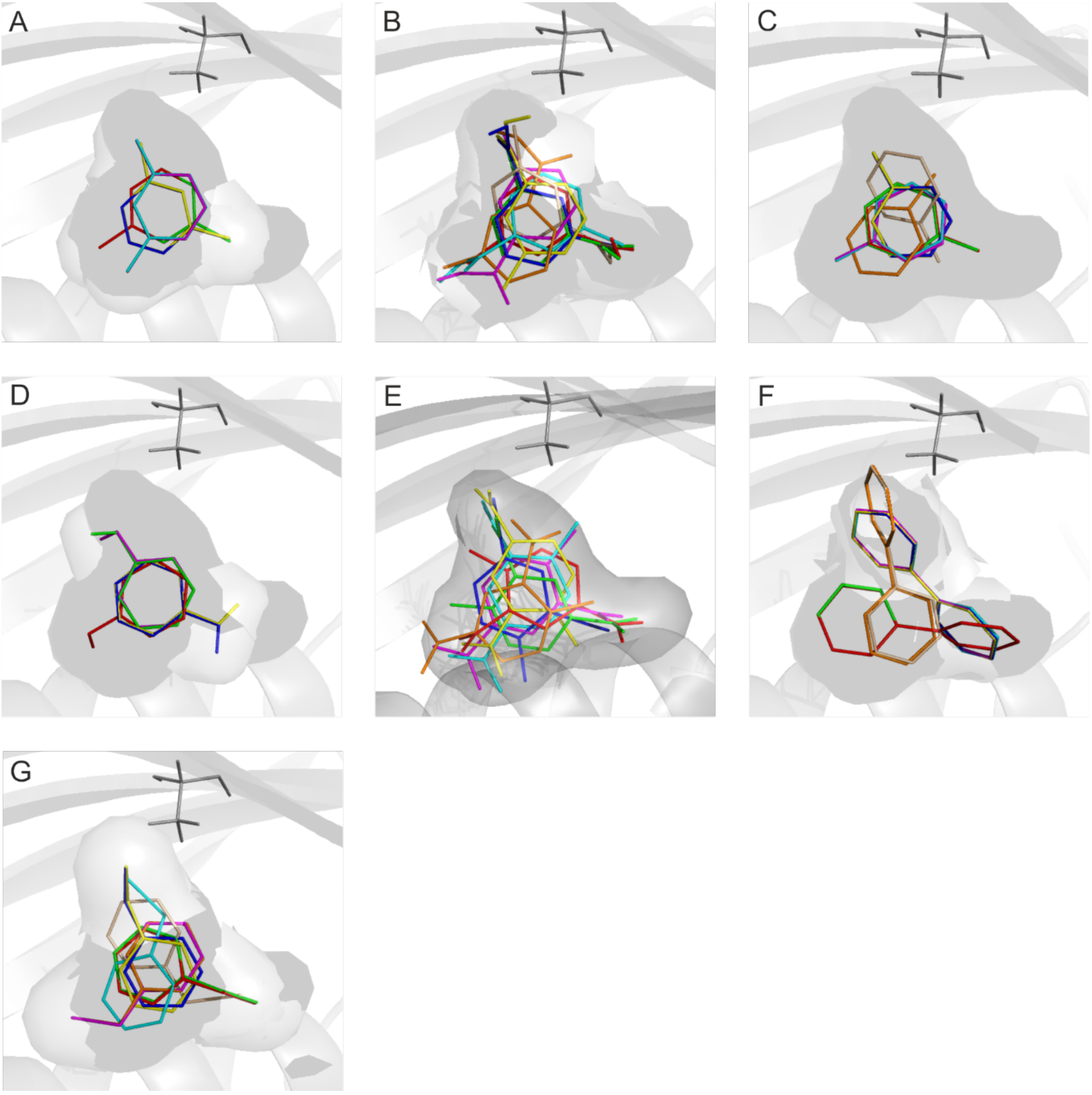


The predicted binding pocket of XylR was docked with three native effector molecules - *m*-xylene (**A**), 3-methylbenzoate (**B**), and toluene (**C**) - and four aromatic ligands which do not activate this transcription factor (Galvão and de Lorenzo, 2006) - phenol (**D**), 2,4-dinitrotoluene (**E**), biphenyl (**F**), and benzyl bromide (**G**) - using Autodock Vina v. 2.2 (Trott and Olson, 2010). Five to eight best-ranked orientations of each ligand are shown. Binding pocket is depicted in transparent surface mode. One of the key residues that shape binding pocket, A126, is shown as grey lines in each panel.

**Figure S4.** *In vivo* evidence of XylR-His interaction with *m*-xylene and benzyl bromide.

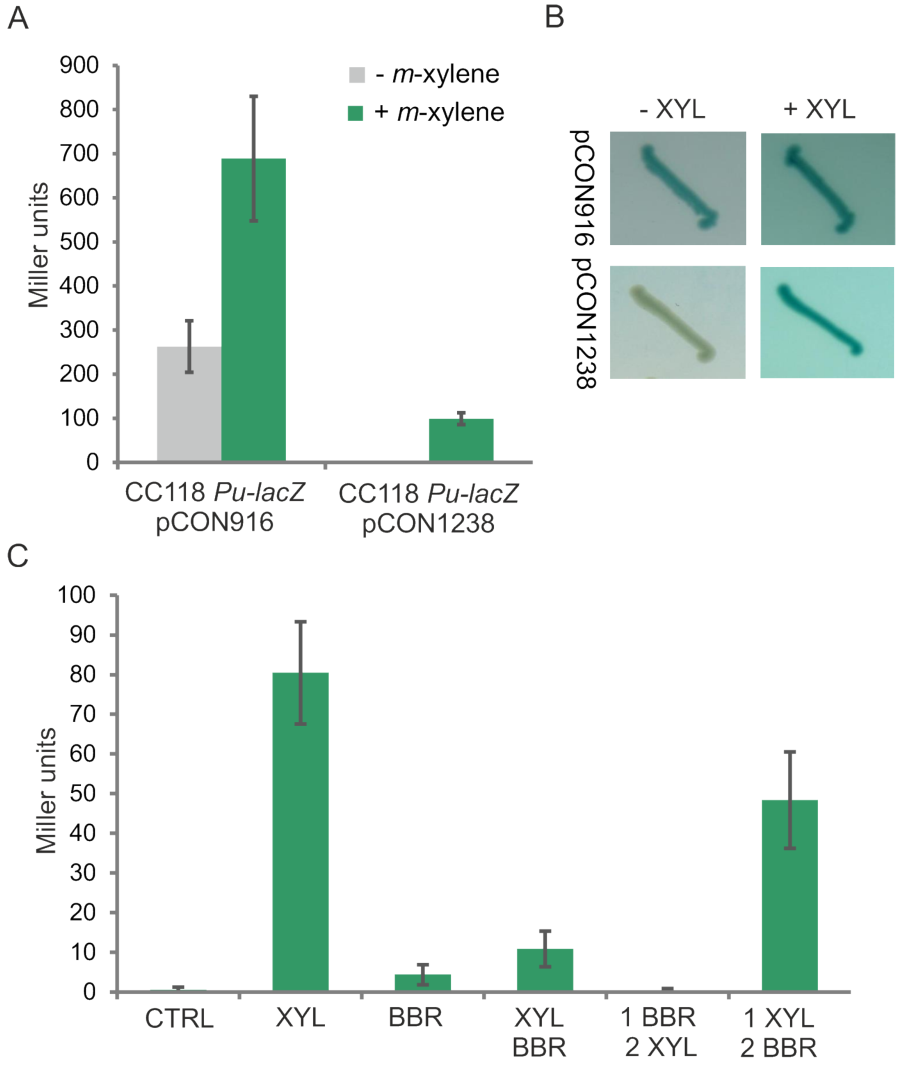


**A)** β-galactosidase activity of *E. coli* CC118 *Pu-lacZ* with pCON916 plasmid bearing wild-type *xylR* and *E. coli* CC118 *Pu-lacZ* with pCON1238 plasmid bearing *xylR-His* in absence or presence of *m*-xylene effector in liquid culture and (**B**) on agar plate. **C)** β-galactosidase activity of *E. coli* CC118 *Pu-lacZ* pCON1238 strain exposed to *m*-xylene and/or benzyl bromide in several specific conditions. Cells were grown to $OD_{600}$ of 1.0 and then were: left uninduced (CTRL), exposed to saturated vapours of *m*-xylene (XYL), benzyl bromide (BBR), or of both chemicals in parallel (XYL/BBR), first exposed to benzyl bromide and then to *m*-xylene (1 BBR/2 XYL), or first exposed to *m*-xylene and then to benzyl bromide (1 XYL/2 BBR). Columns represent means ± SD from at least two independent experiments.

**Figure S5.** Sodium dodecyl sulfate polyacrylamide gel electrophoresis (8 % gel) of purified XylR-His protein.

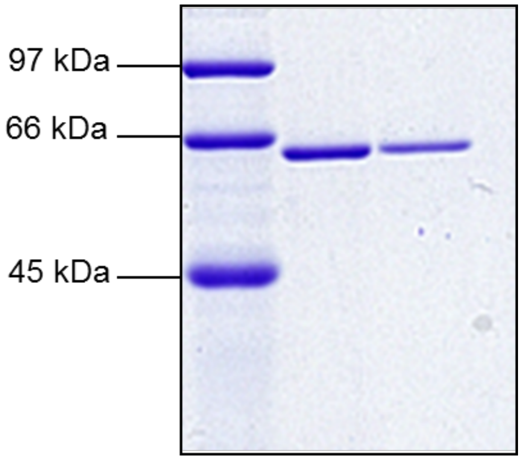


The recombinant *xylR-His* gene was expressed from pCON1238 in *Escherichia coli* BL21(DE3) pLysS and purified as described in Experimental procedures. The theoretical molecular weight of XylR-His is 64.6 kDa.

**Figure S6.** Surface accessibility of amino acids in XylR A domain predicted by NetSurfP 2.0 server (Klausen *et al.*, 2019).

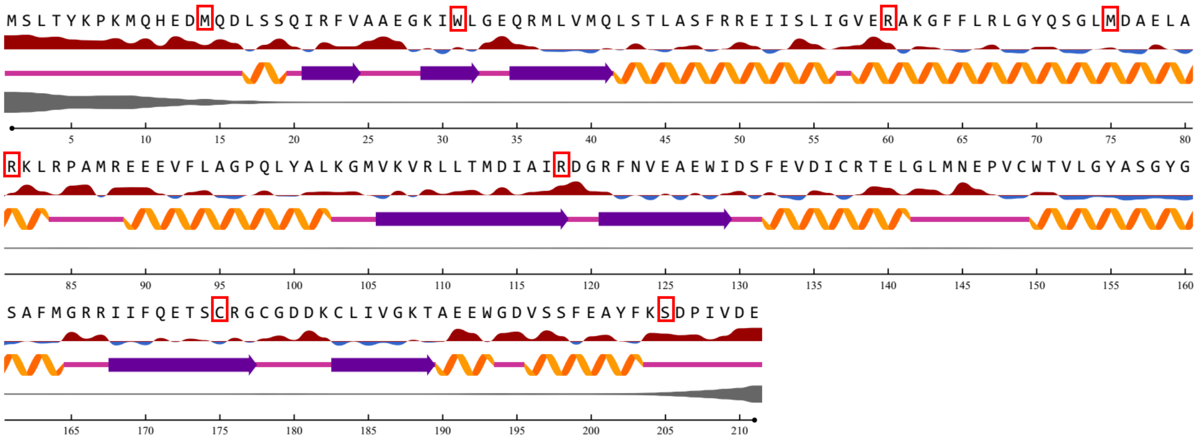


Amino acids (depicted using single letter code) exposed on the protein surface have a red line below them, buried residues have a blue line. The thickness of the line corresponds to the relative surface accessibility of the given residue (25 % threshold). Eight residues modified after protein cross-linking with benzyl bromide are highlighted with red boxes. Secondary structure elements are shown as orange helices (α helices), violet arrows (β strands), and pink lines (loops). The thickness of the grey line equals the probability of disordered residue.
