## Supplemental File 2 for "An updated structural model of the A domain of the *Pseudomonas putida* XylR regulator exposes a distinct interplay with aromatic effectors"

**Supporting information file 2:**
### The mass spectrometry analysis for identification of XylR peptides with nucleophilic amino acids cross-linked with benzyl bromide (BrBz)

Contents


Liquid chromatography conditions……………………………………………..2

Mascot search results for peptides analysed by MALDI-TOF/TOF…………...3

Mass spectra (MS1) of ten modified peptides (MALDI-TOF/TOF)…………….6

Mass spectra (MS2, pages A) and elution profiles (pages B) of five modified peptides and non-modified controls (Ultra Ion Trap)…………………………..16

### SEPARATION OF PEPTIDES WITH PHASE NANO HPLC

**Monolith Trapping column –** PS-DVB, 200-µm i.d. × 5 mm
**Monolith C18 Column –** 150 mm X 0.1 mm

**Loading pump** (20 µl/min)
98 : 2 : 0.1 (H2O:AcN:TFA)

**Nano pump** (1 µl/min)
A.- 98 : 2 : 0.1 (H2O : AcN : TFA)
B.- 20 : 80 : 0.1 (H2O : AcN : TFA)

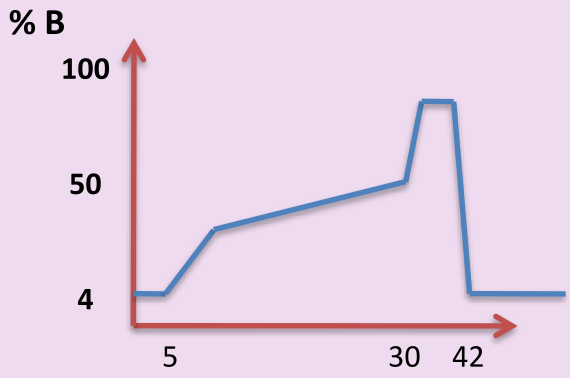


**Grad 30 min**

| Min | %B |
|---|---|
| 0 | 2 |
| 5 | 2 |
| 5.1 | 20 |
| 30 | 50 |
| 36 | 95 |
| 41 | 95 |
| 42 | 2 |
| 47 | 2 |

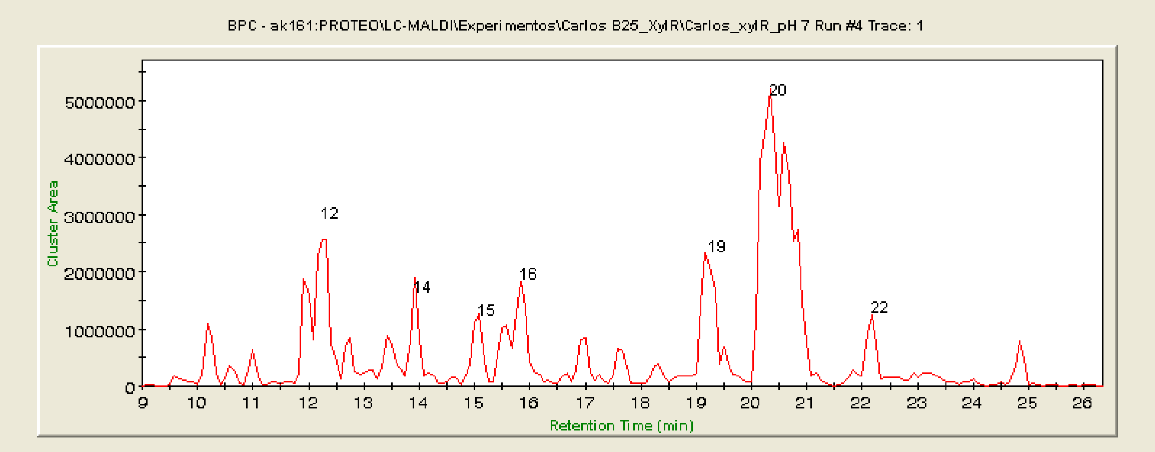

### Mascot Search Results

#### Probability Based Mowse Score

Ions score is -10*Log(P), where P is the probability that the observed match is a random event.
Individual ions scores > 60 indicate identity or extensive homology (p<0.05).
Protein scores are derived from ions scores as a non-probabilistic basis for ranking protein hits.

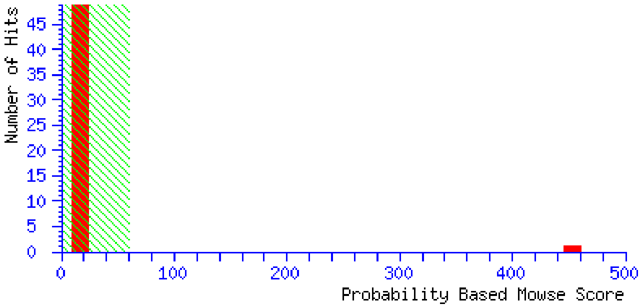


#### Peptide Summary Report

| | | | |
|---|---|---|---|
| Format As | Peptide Summary | | Help |
| | Significance threshold p< 0.05 | Max. number of hits 5 | |
| | Standard scoring ◉ MudPIT scoring ○ | Ions score or expect cut-off 19 | Show sub-sets 0 |
| | Show pop-ups ◉ Suppress pop-ups ○ | Sort unassigned Decreasing Score | Require bold red ☐ |

#### Peptide Summary Report

P06519 **Mass:** 63704 **Score:** 437 **Queries matched:** 14
Transcriptional regulatory protein xylR OS=Pseudomonas putida GN=xylR PE=4 SV=1
☐ Check to include this hit in error tolerant search or archive report

| | Query | Observed | Mr(expt) | Mr(calc) | ppm | Miss | Score | Expect | Rank | Peptide |
|---|---|---|---|---|---|---|---|---|---|---|
| ☑ | 5 | 1135.67 | 1134.66 | 1134.65 | 12.8 | 0 | 35 | 4.9 | 1 | R.LLTMDIAIR.D + BrBz (CHKMRSTWY) |
| ☑ | 9 | 1218.71 | 1217.70 | 1217.70 | -2.40 | 0 | 42 | 1.2 | 1 | R.EIISLIGVER.A + BrBz (CHKMRSTWY) |
| ☑ | 10 | 1424.75 | 1423.74 | 1423.75 | -4.23 | 0 | 65 | 0.012 | 1 | R.YELQTQVANLR.N + BrBz (CHKMRSTWY) |
| ☑ | 13 | 1491.88 | 1490.87 | 1490.88 | -2.59 | 0 | 22 | 1.6e+002 | 1 | R.LNVFPVHIPPLR.E + BrBz (CHKMRSTWY) |
| ☑ | 18 | 1613.80 | 1612.79 | 1612.79 | 1.81 | 0 | (33) | 23 | 1 | R.LGYQSGLMDAELAR.K + BrBz (CHKMRSTWY) |
| ☑ | 21 | 1613.81 | 1612.80 | 1612.79 | 5.66 | 0 | (22) | 3e+002 | 1 | R.LGYQSGLMDAELAR.K + BrBz (CHKMRSTWY) |
| ☑ | 22 | 1613.82 | 1612.81 | 1612.79 | 10.1 | 0 | 98 | 7.3e-006 | 1 | R.LGYQSGLMDAELAR.K + BrBz (CHKMRSTWY) |
| ☑ | 24 | 1634.85 | 1633.84 | 1633.86 | -10.24 | 0 | 60 | 0.05 | 1 | R.LITATNENLEEAVK.M + BrBz (CHKMRSTWY) |
| ☑ | 25 | 1634.86 | 1633.85 | 1633.86 | -2.23 | 0 | (59) | 0.068 | 1 | R.LITATNENLEEAVK.M + BrBz (CHKMRSTWY) |
| ☑ | 34 | 1807.82 | 1806.81 | 1806.80 | 3.62 | 0 | 52 | 0.35 | 1 | K.MQHEDMQDLSSQIR.F + BrBz (CHKMRSTWY) |
| ☑ | 39 | 1935.89 | 1934.89 | 1934.89 | -3.18 | 0 | 36 | 19 | 1 | R.AMEACLHYQWPGNIR.E + BrBz (CHKMRSTWY); Carbamidomethyl (C) |
| ☑ | 40 | 1935.91 | 1934.90 | 1934.89 | 4.01 | 0 | (26) | 1.9e+002 | 1 | R.AMEACLHYQWPGNIR.E + BrBz (CHKMRSTWY); Carbamidomethyl (C) |
| ☑ | 44 | 1937.88 | 1936.87 | 1936.89 | -7.57 | 0 | 72 | 0.0048 | 1 | K.QYDGQYYGIGHSPAYK.R + BrBz 91 (DEM) |
| ☑ | 61 | 2077.06 | 2076.05 | 2076.06 | -1.73 | 0 | 20 | 9.8e+002 | 1 | R.QIIDQGVSLEDLEAGLMR.T + BrBz (CHKMRSTWY) |

P06519 **Mass:** 63704 **Score:** 478 **Expect:** 6.3e-043 **Queries matched:** 64

Transcriptional regulatory protein xylR OS=Pseudomonas putida GN=xylR PE=4 SV=1

| Observed | Mr(expt) | Mr(calc) | ppm | Start | End | Miss | Ions | Peptide |
|---|---|---|---|---|---|---|---|---|
| 1063.54 | 1062.53 | 1062.53 | -0.76 | 454 - | 461 | 0 | --- | R.ELENALER.G + BrBz (CHKMRSTWY) |
| 1063.55 | 1062.55 | 1062.53 | 11.6 | 454 - | 461 | 0 | --- | R.ELENALER.G + BrBz (CHKMRSTWY) |
| 1063.56 | 1062.55 | 1062.53 | 19.0 | 454 - | 461 | 0 | --- | R.ELENALER.G + BrBz (CHKMRSTWY) |
| 1135.65 | 1134.64 | 1134.65 | -3.75 | 110 - | 118 | 0 | (16) | R.LLTMDIAIR.D + BrBz (CHKMRSTWY) |
| 1135.67 | 1134.66 | 1134.65 | 12.8 | 110 - | 118 | 0 | 35 | R.LLTMDIAIR.D + BrBz (CHKMRSTWY) |
| 1162.61 | 1161.60 | 1161.60 | 1.08 | 350 - | 358 | 0 | 19 | R.VLQEGELER.V + BrBz (CHKMRSTWY) |
| 1162.61 | 1161.61 | 1161.60 | 4.01 | 350 - | 358 | 0 | --- | R.VLQEGELER.V + BrBz (CHKMRSTWY) |
| 1162.63 | 1161.62 | 1161.60 | 13.7 | 350 - | 358 | 0 | --- | R.VLQEGELER.V + BrBz (CHKMRSTWY) |
| 1218.71 | 1217.70 | 1217.70 | -2.40 | 51 - | 60 | 0 | 42 | R.EIISLIGVER.A + BrBz (CHKMRSTWY) |
| 1424.75 | 1423.74 | 1423.75 | -4.23 | 213 - | 223 | 0 | 65 | R.YELQTQVANLR.N + BrBz (CHKMRSTWY) |
| 1424.75 | 1423.75 | 1423.75 | 0.48 | 213 - | 223 | 0 | (8) | R.YELQTQVANLR.N + BrBz (CHKMRSTWY) |
| 1491.74 | 1490.73 | 1490.88 | -96.63 | 396 - | 407 | 0 | --- | R.LNVFPVHIPPLR.E + BrBz (CHKMRSTWY) |
| 1491.88 | 1490.87 | 1490.88 | -2.59 | 396 - | 407 | 0 | 22 | R.LNVFPVHIPPLR.E + BrBz (CHKMRSTWY) |
| 1613.79 | 1612.78 | 1612.84 | -32.97 | 167 - | 176 | 1 | 3 | R.RIIFQETSCR.G + 3 BrBz (CHKMRSTWY); BrBz 91 (DEM) |
| 1613.80 | 1612.79 | 1612.79 | -1.47 | 68 - | 81 | 0 | (5) | R.LGYQSGLMDAELAR.K + BrBz (CHKMRSTWY) |
| 1613.80 | 1612.79 | 1612.79 | -0.98 | 68 - | 81 | 0 | (13) | R.LGYQSGLMDAELAR.K + BrBz (CHKMRSTWY) |
| 1613.80 | 1612.79 | 1612.79 | 1.19 | 68 - | 81 | 0 | (11) | R.LGYQSGLMDAELAR.K + BrBz (CHKMRSTWY) |
| 1613.80 | 1612.79 | 1612.79 | 1.81 | 68 - | 81 | 0 | (33) | R.LGYQSGLMDAELAR.K + BrBz (CHKMRSTWY) |
| 1613.80 | 1612.79 | 1612.79 | 1.87 | 68 - | 81 | 0 | (11) | R.LGYQSGLMDAELAR.K + BrBz (CHKMRSTWY) |
| 1613.80 | 1612.80 | 1612.79 | 3.55 | 68 - | 81 | 0 | (18) | R.LGYQSGLMDAELAR.K + BrBz (CHKMRSTWY) |
| 1613.81 | 1612.80 | 1612.79 | 5.66 | 68 - | 81 | 0 | (22) | R.LGYQSGLMDAELAR.K + BrBz (CHKMRSTWY) |
| 1613.82 | 1612.81 | 1612.79 | 10.1 | 68 - | 81 | 0 | 98 | R.LGYQSGLMDAELAR.K + BrBz (CHKMRSTWY) |
| 1613.83 | 1612.83 | 1612.84 | -10.52 | 1 - | 8 | 0 | --- | -.MSLTYKPK.M + 3 BrBz (CHKMRSTWY); 2 BrBz(180) (K); Oxidation (M) |
| 1634.85 | 1633.84 | 1633.86 | -10.24 | 371 - | 384 | 0 | 60 | R.LITATNENLEEAVK.M + BrBz (CHKMRSTWY) |
| 1634.86 | 1633.85 | 1633.86 | -2.23 | 371 - | 384 | 0 | (59) | R.LITATNENLEEAVK.M + BrBz (CHKMRSTWY) |
| 1664.87 | 1663.86 | 1663.97 | -64.92 | 255 - | 269 | 1 | --- | R.GRVSVLLLGETGVGK.E + 2 BrBz (CHKMRSTWY) |
| 1669.89 | 1668.89 | 1668.85 | 23.1 | 167 - | 176 | 1 | --- | R.RIIFQETSCR.G + 4 BrBz (CHKMRSTWY); Carbamidomethyl (C) |
| 1669.90 | 1668.89 | 1668.85 | 26.6 | 167 - | 176 | 1 | --- | R.RIIFQETSCR.G + 4 BrBz (CHKMRSTWY); Carbamidomethyl (C) |
| 1669.98 | 1668.97 | 1668.85 | 72.9 | 167 - | 176 | 1 | --- | R.RIIFQETSCR.G + 4 BrBz (CHKMRSTWY); Carbamidomethyl (C) |
| 1796.82 | 1795.82 | 1795.91 | -52.43 | 68 - | 81 | 0 | --- | R.LGYQSGLMDAELAR.K + 3 BrBz 91 (DEM) |
| 1796.84 | 1795.83 | 1795.91 | -42.97 | 68 - | 81 | 0 | --- | R.LGYQSGLMDAELAR.K + 3 BrBz 91 (DEM) |
| 1807.82 | 1806.81 | 1806.80 | 3.06 | 9 - | 22 | 0 | (4) | K.MQHEDMQDLSSQIR.F + BrBz (CHKMRSTWY) |
| 1807.82 | 1806.81 | 1806.80 | 3.62 | 9 - | 22 | 0 | 52 | K.MQHEDMQDLSSQIR.F + BrBz (CHKMRSTWY) |

| | | | | | | | |
|---|---|---|---|---|---|---|---|
| 1935.86 | 1934.85 | 1935.02 | -86.89 | 326 - 342 | 0 | --- | R.ANGGTIFLDEVIELTPR.A + BrBz 91 (DEM) |
| 1935.89 | 1934.89 | 1934.89 | -3.18 | 439 - 453 | 0 | 36 | R.AMEACLHYQWPGNIR.E + BrBz (CHKMRSTWY); Carbamidomethyl (C) |
| 1935.91 | 1934.90 | 1934.89 | 4.01 | 439 - 453 | 0 | (26) | R.AMEACLHYQWPGNIR.E + BrBz (CHKMRSTWY); Carbamidomethyl (C) |
| 1935.91 | 1934.90 | 1935.02 | -61.82 | 326 - 342 | 0 | --- | R.ANGGTIFLDEVIELTPR.A + BrBz 91 (DEM) |
| 1937.81 | 1936.81 | 1936.98 | -87.74 | 68 - 82 | 1 | --- | R.LGYQSGLMDAELARK.L + 3 BrBz (CHKMRSTWY); Oxidation (M) |
| 1937.88 | 1936.87 | 1936.89 | -7.93 | 228 - 243 | 0 | (7) | K.QYDGQYYGIGHSPAYK.R + BrBz 91 (DEM) |
| 1937.88 | 1936.87 | 1936.89 | -7.57 | 228 - 243 | 0 | 72 | K.QYDGQYYGIGHSPAYK.R + BrBz 91 (DEM) |
| 1937.88 | 1936.87 | 1936.98 | -52.07 | 68 - 82 | 1 | --- | R.LGYQSGLMDAELARK.L + 3 BrBz (CHKMRSTWY); Oxidation (M) |
| 1939.86 | 1938.85 | 1939.03 | -93.38 | 37 - 50 | 1 | --- | R.MLVMQLSTLASFRR.E + 2 BrBz (CHKMRSTWY); BrBz 91 (DEM); Oxidation (M) |
| 1939.89 | 1938.88 | 1939.03 | -78.78 | 37 - 50 | 1 | --- | R.MLVMQLSTLASFRR.E + 2 BrBz (CHKMRSTWY); BrBz 91 (DEM); Oxidation (M) |
| 1939.90 | 1938.89 | 1939.03 | -74.91 | 37 - 50 | 1 | --- | R.MLVMQLSTLASFRR.E + 2 BrBz (CHKMRSTWY); BrBz 91 (DEM); Oxidation (M) |
| 1939.91 | 1938.91 | 1939.03 | -65.89 | 37 - 50 | 1 | --- | R.MLVMQLSTLASFRR.E + 2 BrBz (CHKMRSTWY); BrBz 91 (DEM); Oxidation (M) |
| 1939.99 | 1938.99 | 1939.03 | -24.63 | 37 - 50 | 1 | --- | R.MLVMQLSTLASFRR.E + 2 BrBz (CHKMRSTWY); BrBz 91 (DEM); Oxidation (M) |
| 1951.89 | 1950.89 | 1950.89 | 0.33 | 439 - 453 | 0 | --- | R.AMEACLHYQWPGNIR.E + BrBz (CHKMRSTWY); Carbamidomethyl (C); Oxidation (M) |
| 1951.90 | 1950.89 | 1950.89 | 2.12 | 439 - 453 | 0 | (2) | R.AMEACLHYQWPGNIR.E + BrBz (CHKMRSTWY); Carbamidomethyl (C); Oxidation (M) |
| 1951.90 | 1950.89 | 1950.89 | 2.74 | 439 - 453 | 0 | (8) | R.AMEACLHYQWPGNIR.E + BrBz (CHKMRSTWY); Carbamidomethyl (C); Oxidation (M) |
| 1951.90 | 1950.90 | 1950.89 | 4.32 | 439 - 453 | 0 | --- | R.AMEACLHYQWPGNIR.E + BrBz (CHKMRSTWY); Carbamidomethyl (C); Oxidation (M) |
| 1951.91 | 1950.90 | 1950.89 | 7.50 | 439 - 453 | 0 | (1) | R.AMEACLHYQWPGNIR.E + BrBz (CHKMRSTWY); Carbamidomethyl (C); Oxidation (M) |
| 1955.82 | 1954.81 | 1954.86 | -25.05 | 189 - 204 | 0 | --- | K.TAEEWGDVSSFEAYFK.S + BrBz (CHKMRSTWY) |
| 1968.92 | 1967.91 | 1967.92 | -2.76 | 439 - 453 | 0 | --- | R.AMEACLHYQWPGNIR.E + 2 BrBz (CHKMRSTWY) |
| 1968.94 | 1967.93 | 1967.92 | 7.35 | 439 - 453 | 0 | --- | R.AMEACLHYQWPGNIR.E + 2 BrBz (CHKMRSTWY) |
| 2073.95 | 2072.95 | 2072.94 | 1.50 | 490 - 506 | 1 | 12 | R.LSSEGRLEEESGDSWFR.Q + BrBz (CHKMRSTWY) |
| 2076.96 | 2075.95 | 2076.06 | -50.86 | 507 - 524 | 0 | --- | R.QIIDQGVSLEDLEAGLMR.T + BrBz (CHKMRSTWY) |
| 2077.06 | 2076.05 | 2076.06 | -1.73 | 507 - 524 | 0 | 20 | R.QIIDQGVSLEDLEAGLMR.T + BrBz (CHKMRSTWY) |
| 2178.07 | 2177.07 | 2177.06 | 3.38 | 226 - 243 | 1 | --- | R.LKQYDGQYYGIGHSPAYK.R + BrBz (CHKMRSTWY) |
| 2178.08 | 2177.07 | 2177.06 | 5.03 | 226 - 243 | 1 | --- | R.LKQYDGQYYGIGHSPAYK.R + BrBz (CHKMRSTWY) |
| 2178.10 | 2177.09 | 2177.06 | 16.0 | 226 - 243 | 1 | 5 | R.LKQYDGQYYGIGHSPAYK.R + BrBz (CHKMRSTWY) |
| 2521.37 | 2520.37 | 2520.28 | 34.8 | 63 - 81 | 1 | --- | K.GFFLRLGYQSGLMDAELAR.K + 3 BrBz (CHKMRSTWY); BrBz 91 (DEM); Oxidation (M) |

**No match to:** 1664.79

Pages 7-15 summarize some examples of MALDI-TOF/TOF MS1 spectra showing BrBz modification (+90 Da) of peptides obtained after tryptic digestion of protein XylR. Examples of unmodified (purple boxes) and BrBz-modified (+90 Da; light blue boxes) peptides are highlighted.

#### Peptide 1135 Da

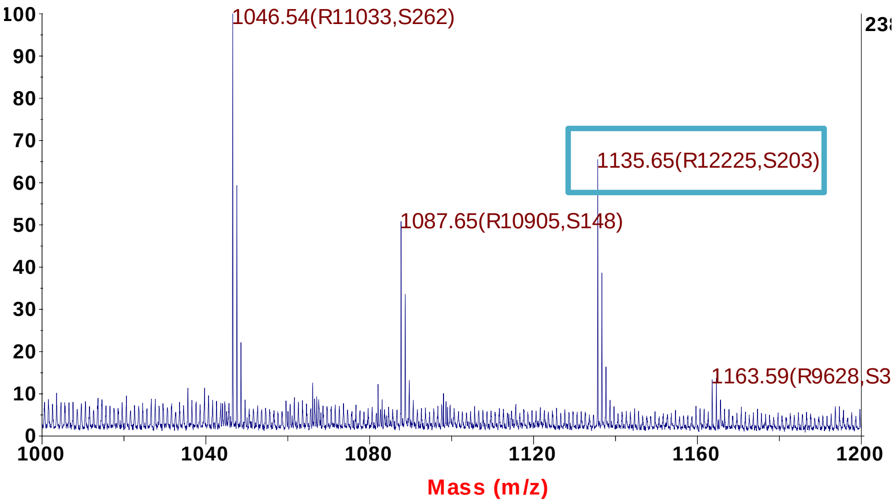


#### Peptide 2077 Da

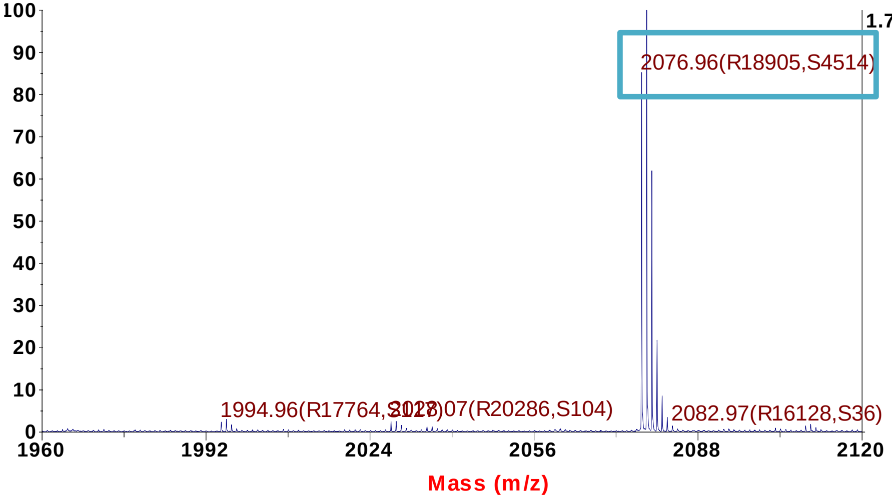


#### Peptide 1135 Da

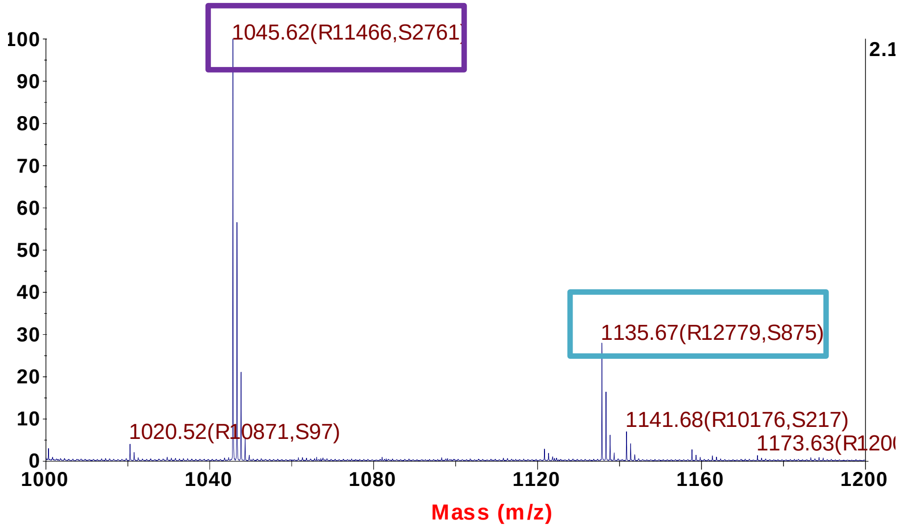


#### Peptide 2077 Da

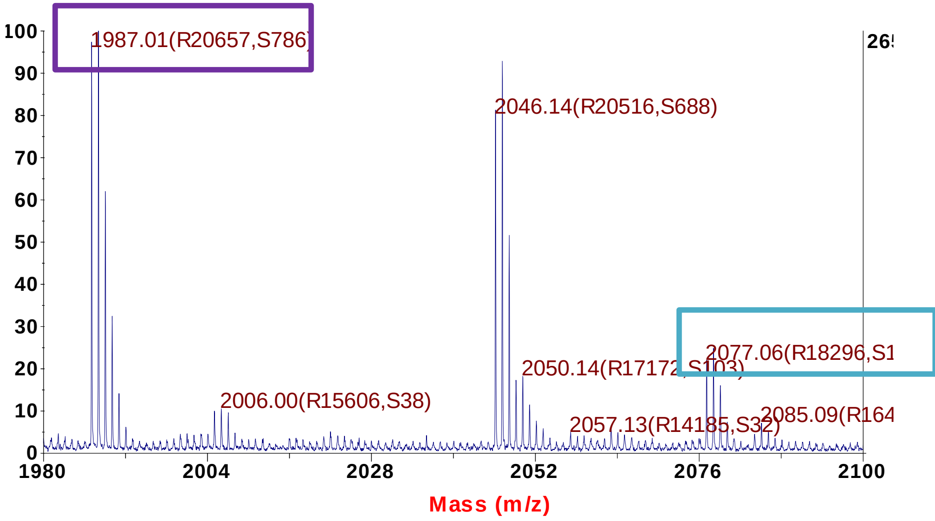

#### Peptide 1613 Da

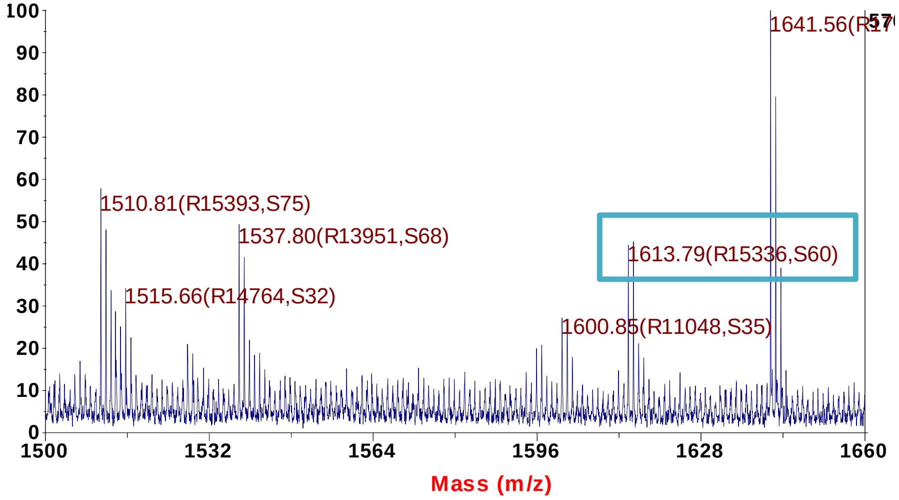


#### Peptide 1613 Da

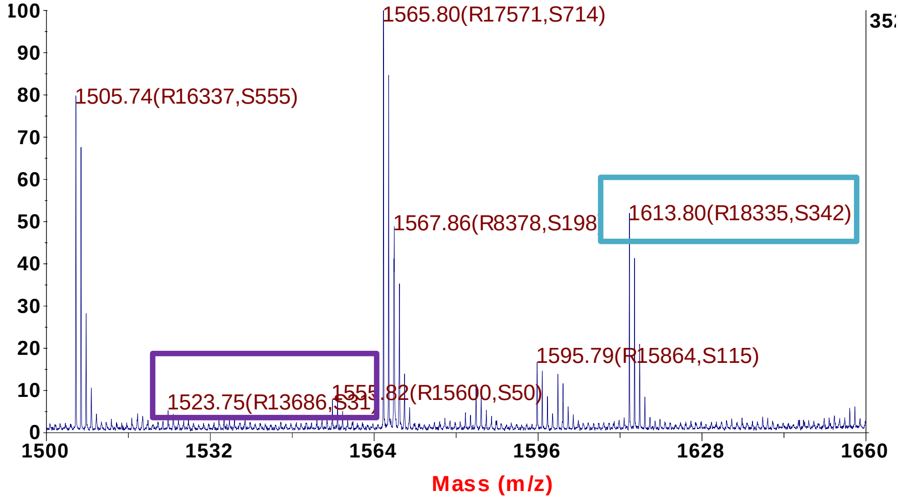


#### Peptide 1613 Da

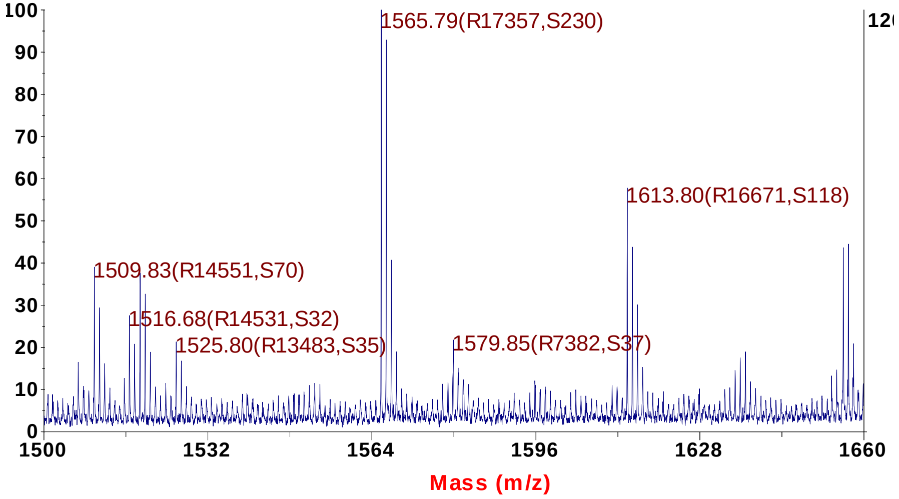


#### Peptide 1613 Da

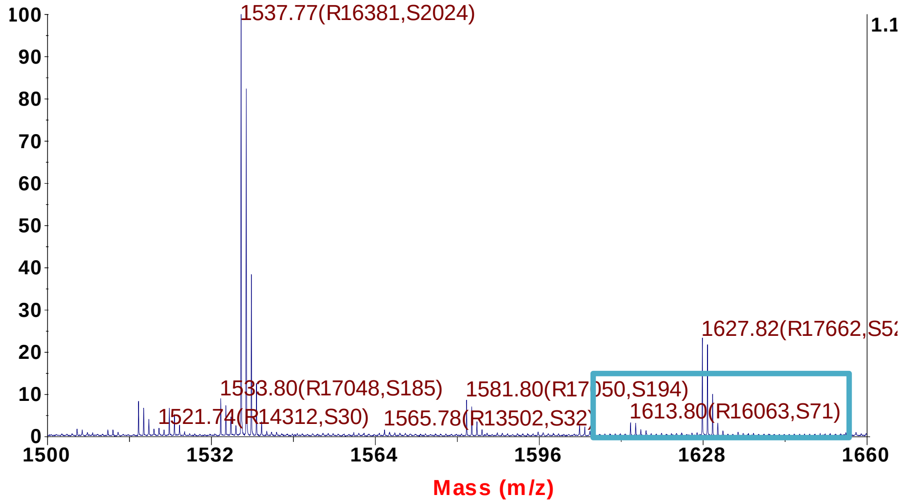

#### Peptide 1613 Da

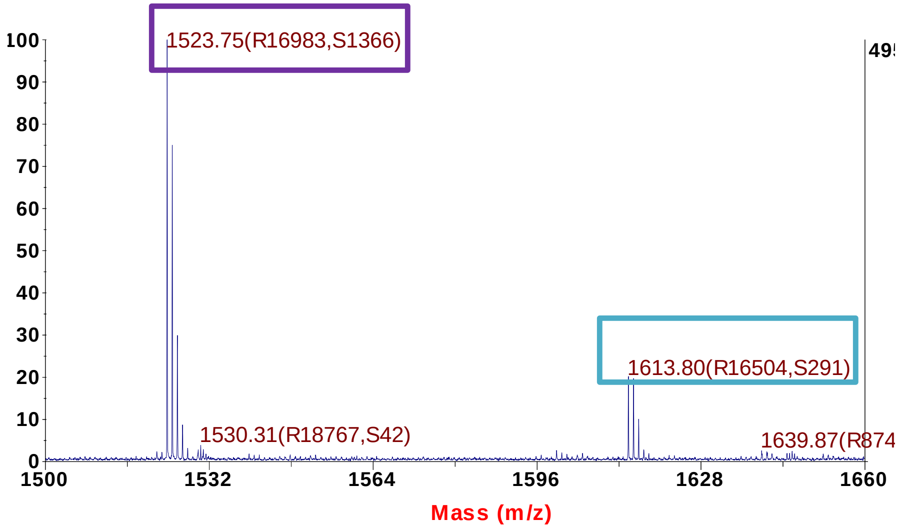


#### Peptide 1613 Da

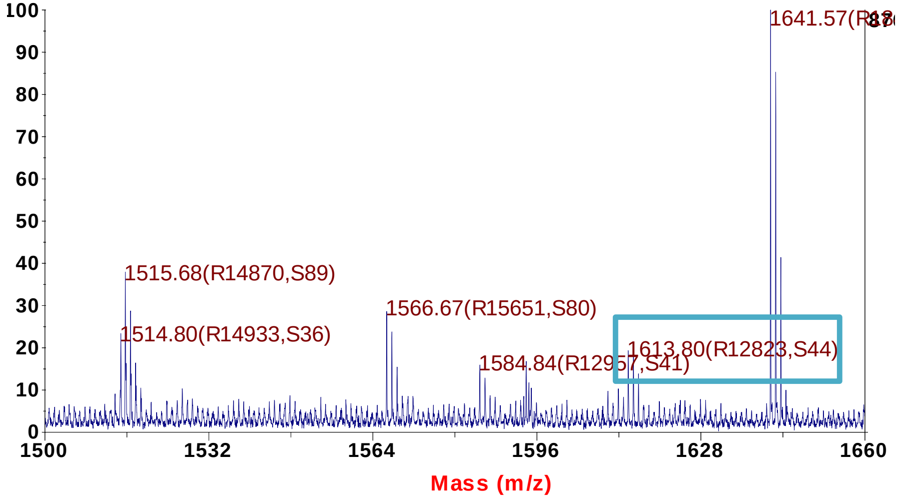


#### Peptide 1613 Da

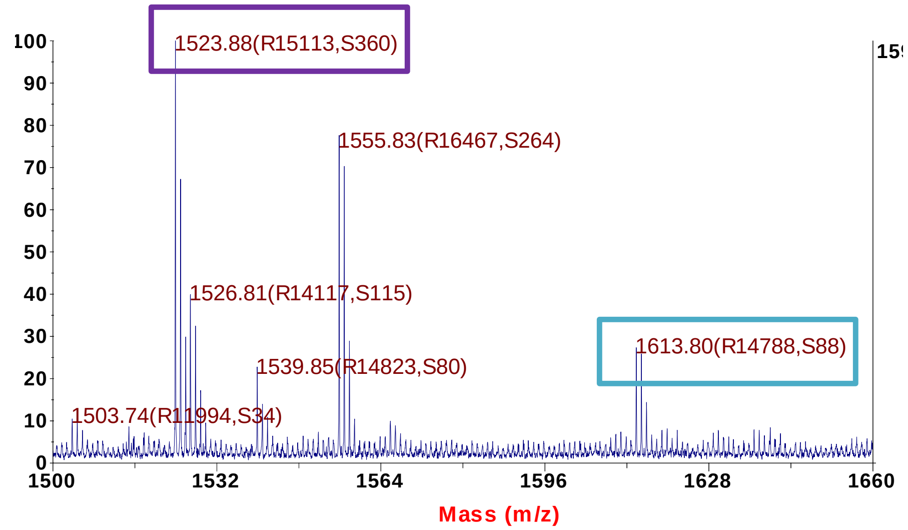


#### Peptide 1613 Da

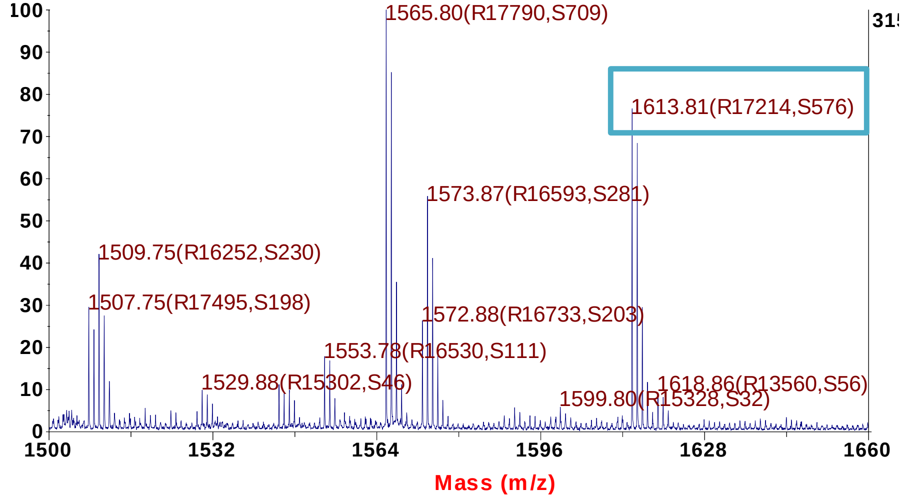

#### Peptide 1613 Da

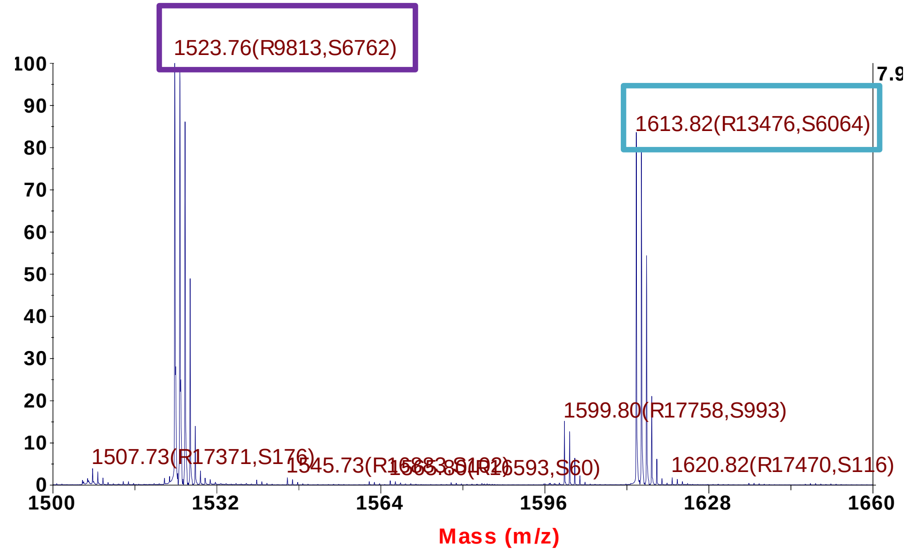


#### Peptide 1807 Da

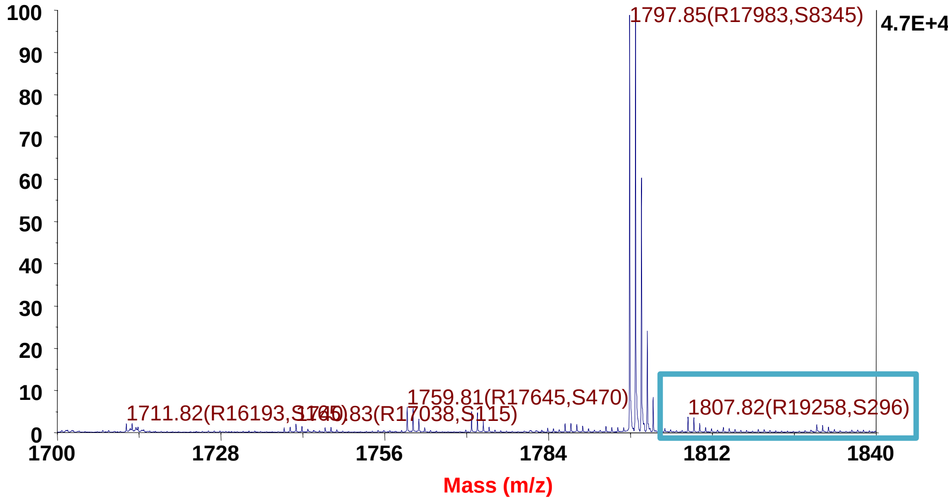


#### Peptide 1613 Da

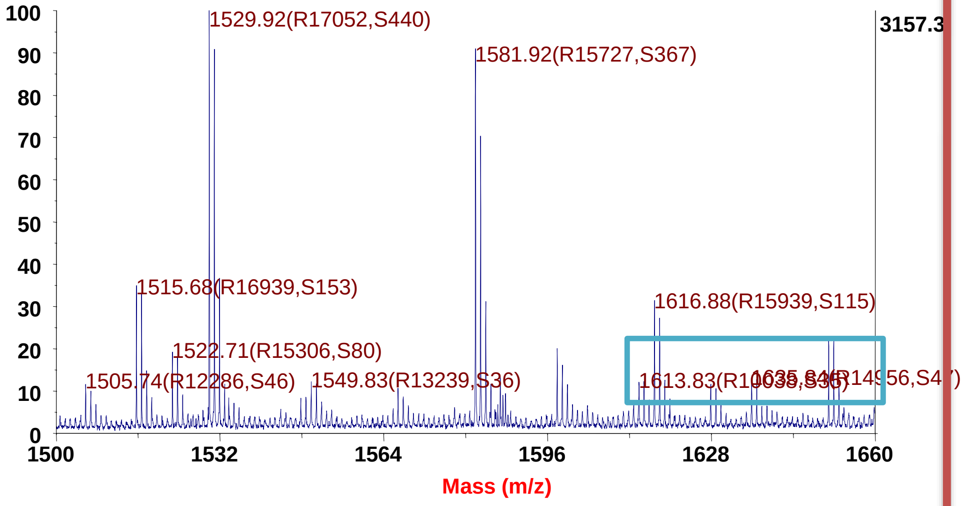


#### Peptide 1807 Da

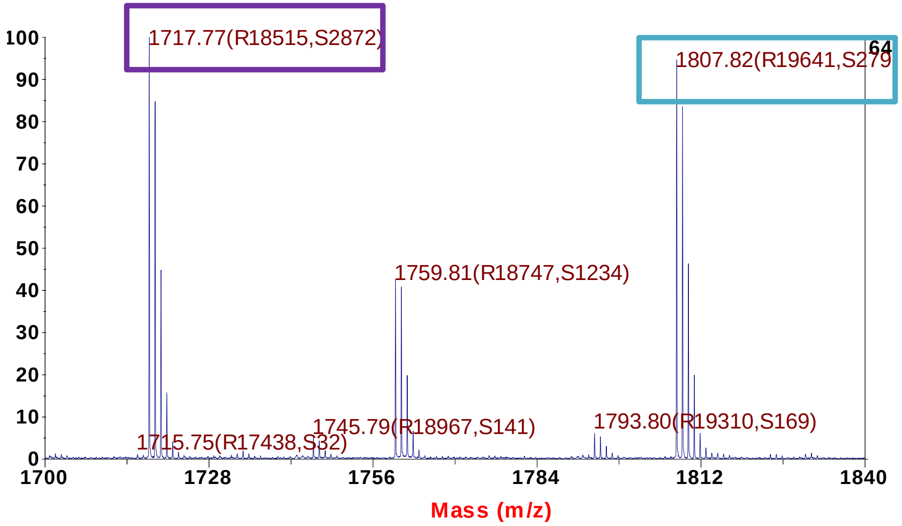

### Peptide 1807 Da

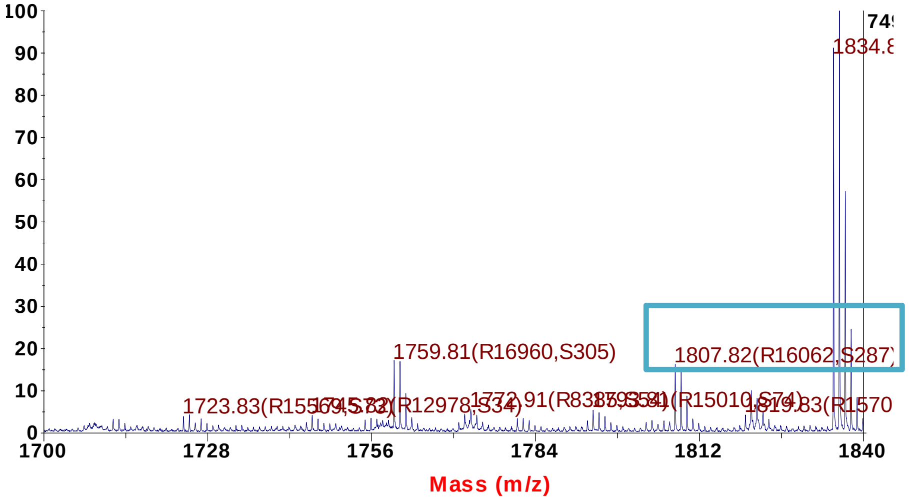


### Peptide 1218 Da

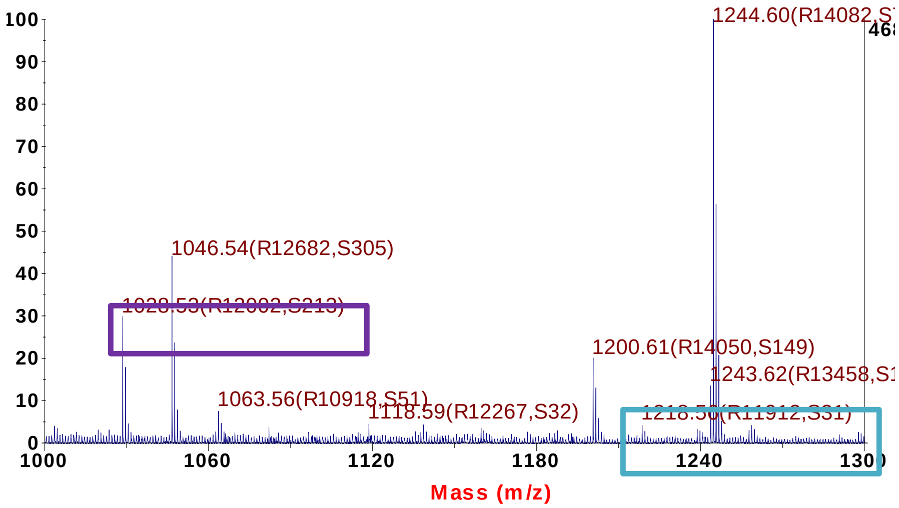


### Peptide 1491 Da

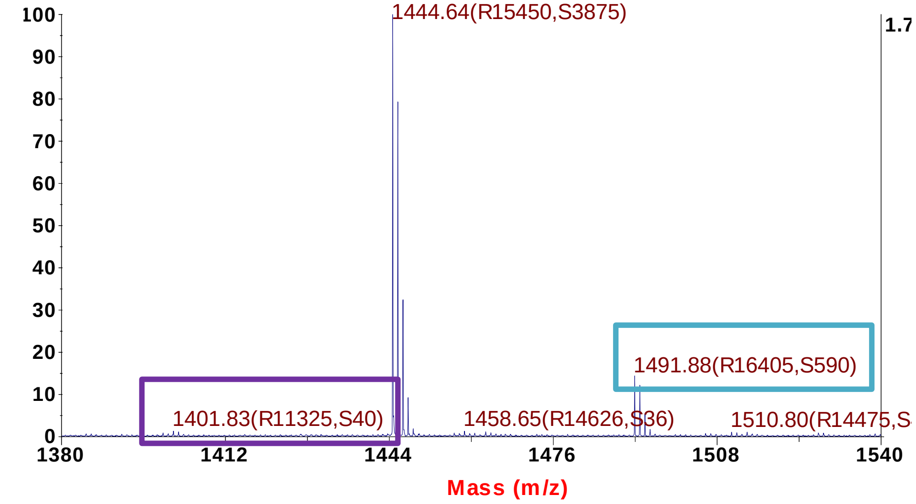


### Peptide 1218 Da

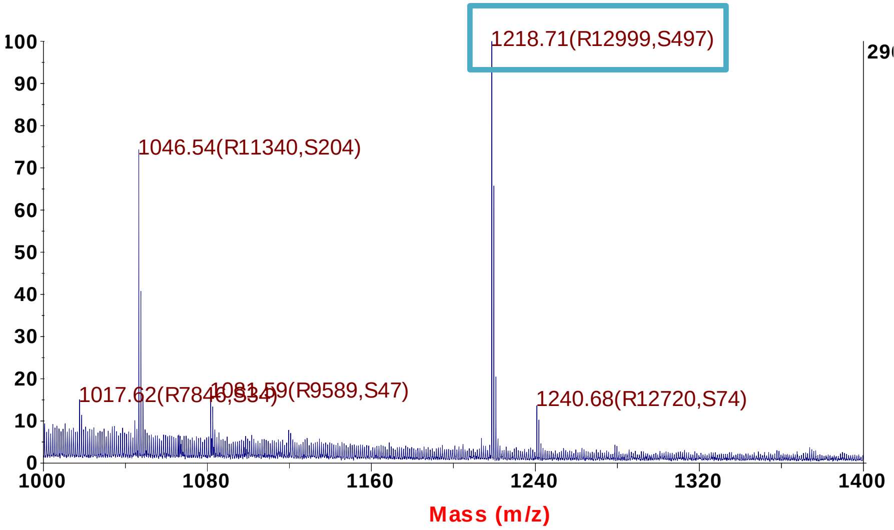

#### Peptide 1424 Da

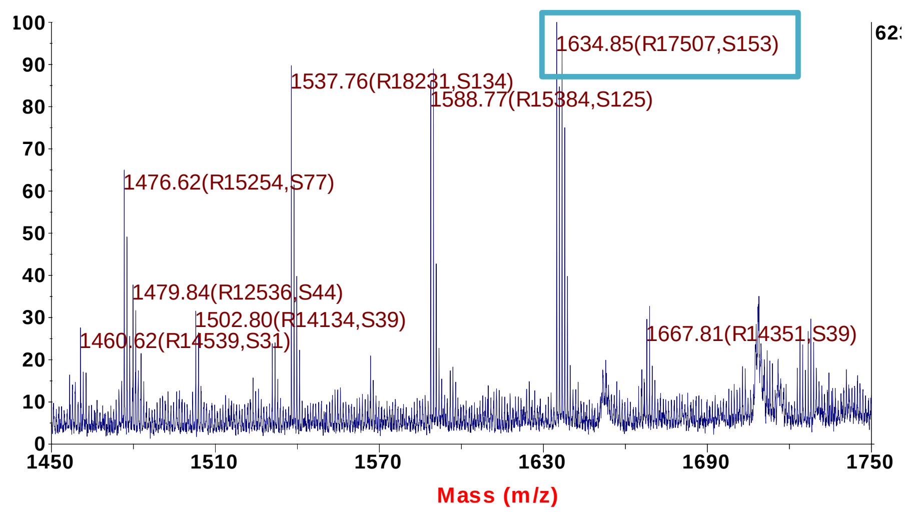


#### Peptide 1424 Da

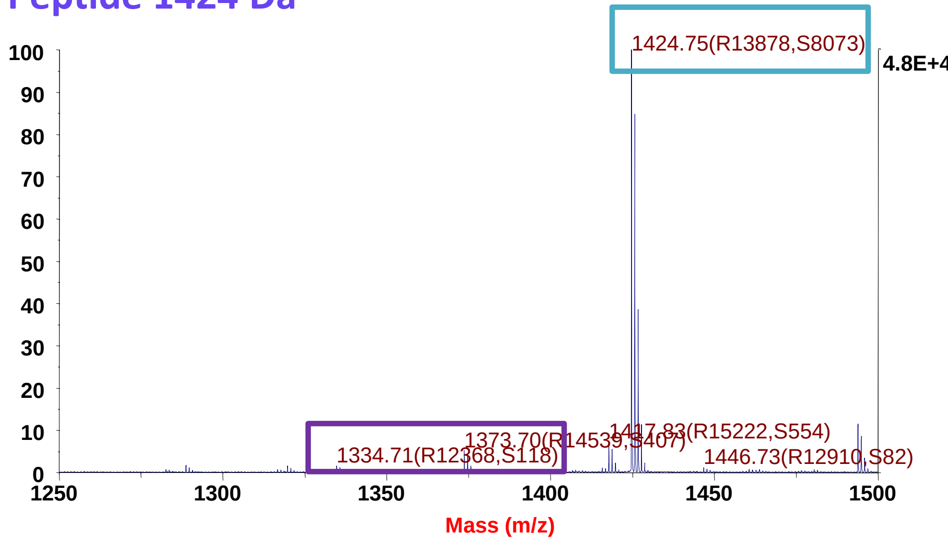


#### Peptide 1424 Da

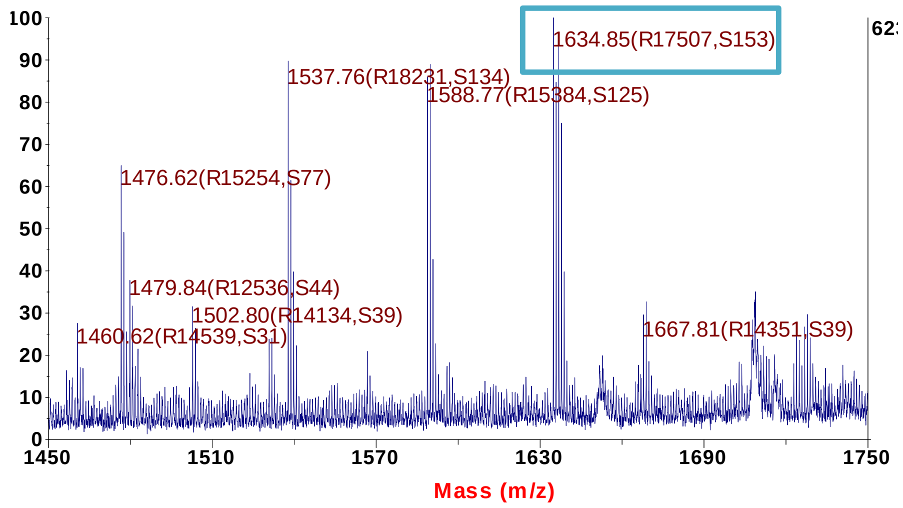


#### Peptide 1424 Da

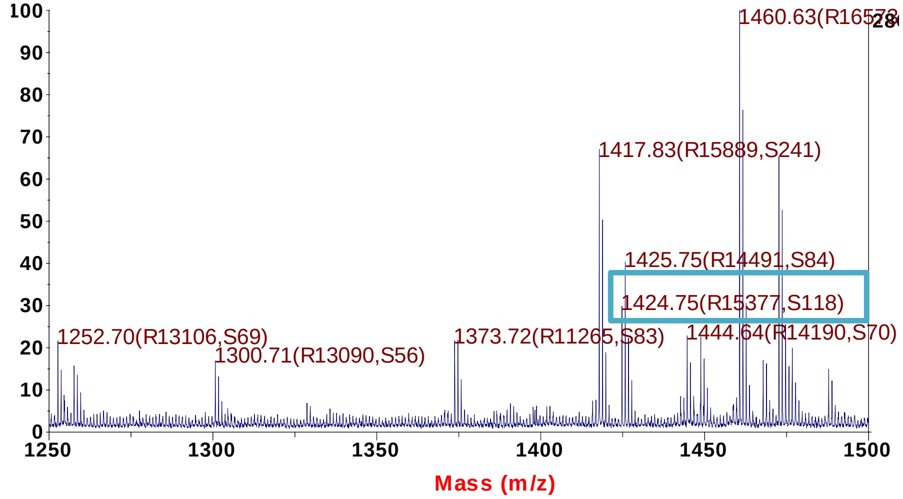

#### Peptide 1634 Da

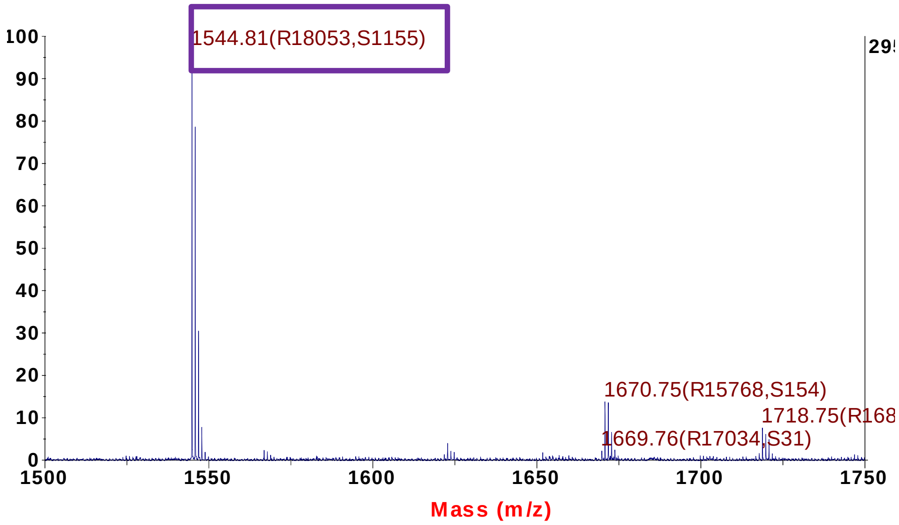


#### Peptide 1634 Da

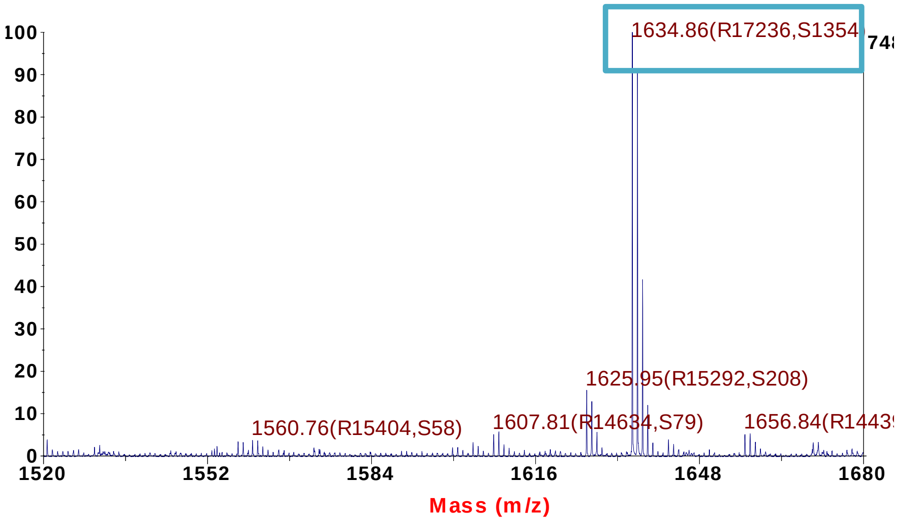


#### Peptide 1634 Da

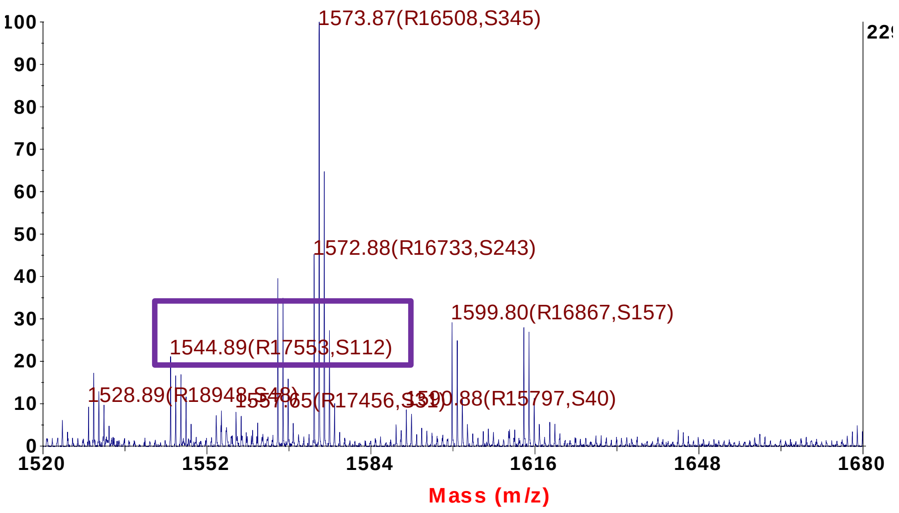


#### Peptide 1634 Da

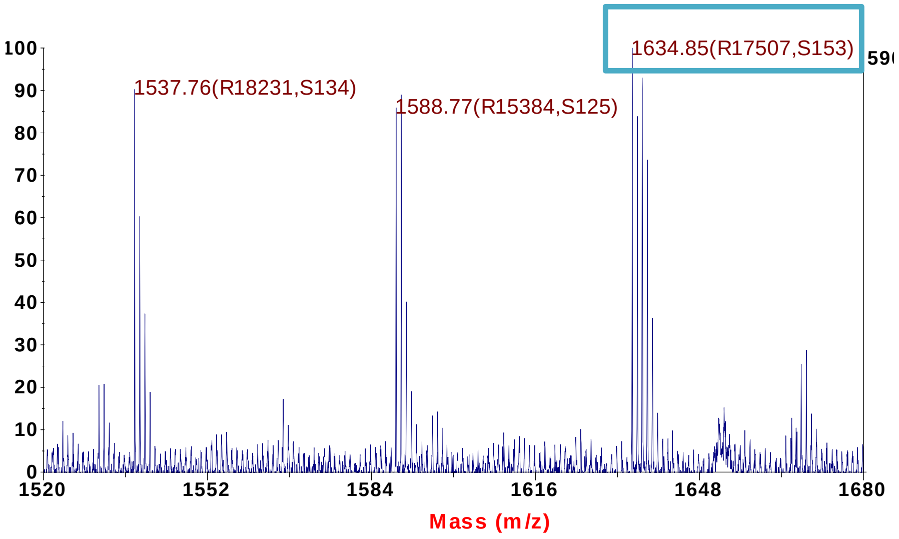

### Peptide 1937 Da

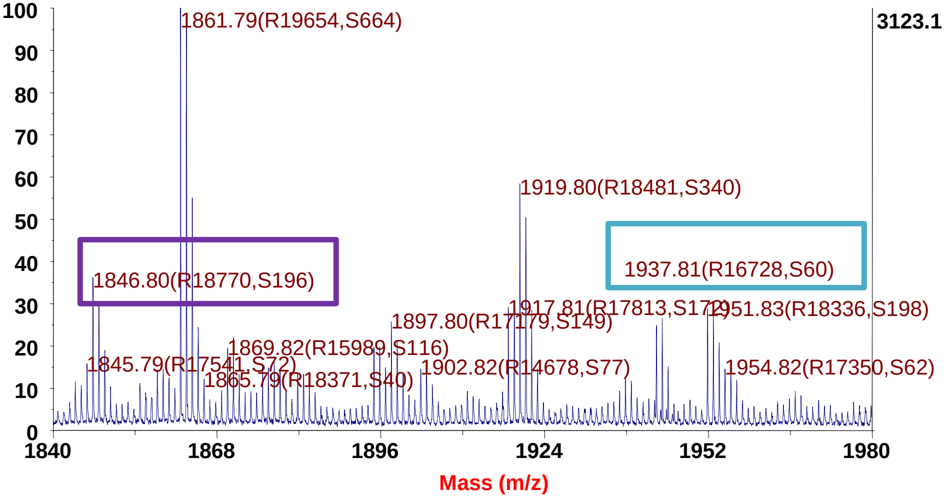


### Peptide 1937 Da

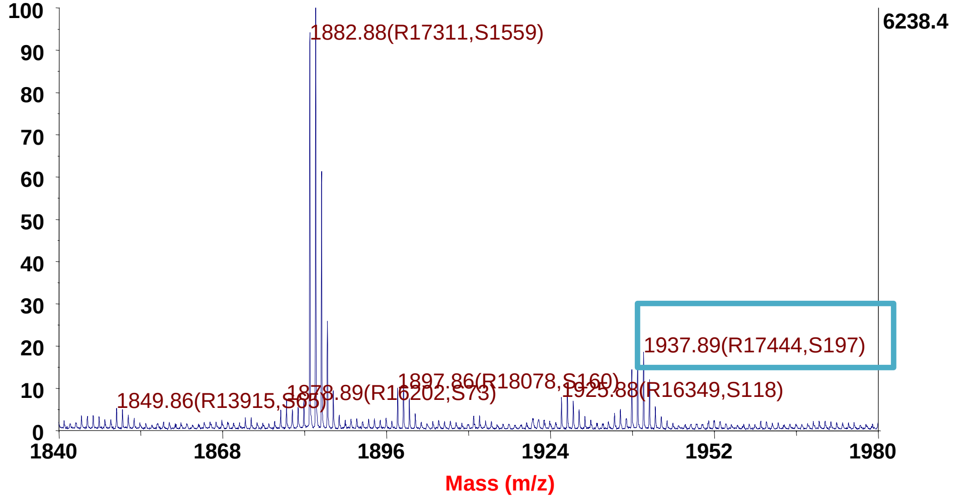


### Peptide 1937 Da

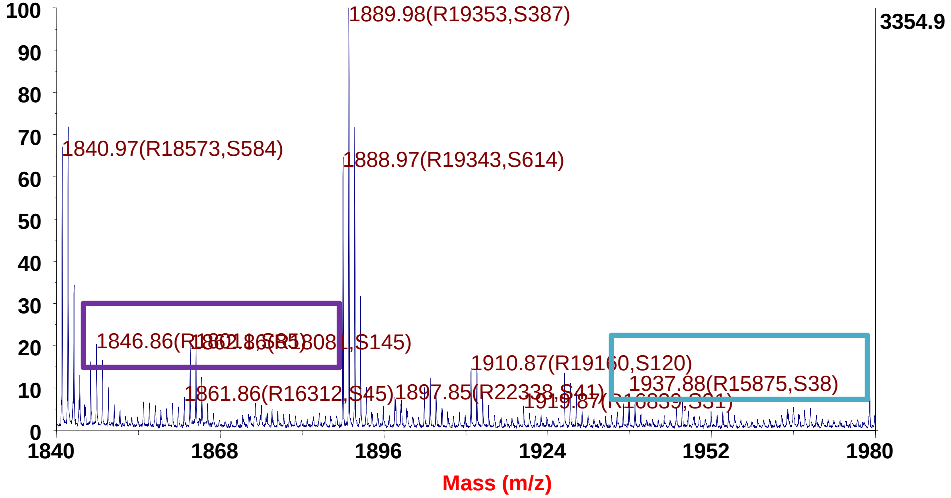


### Peptide 1937 Da

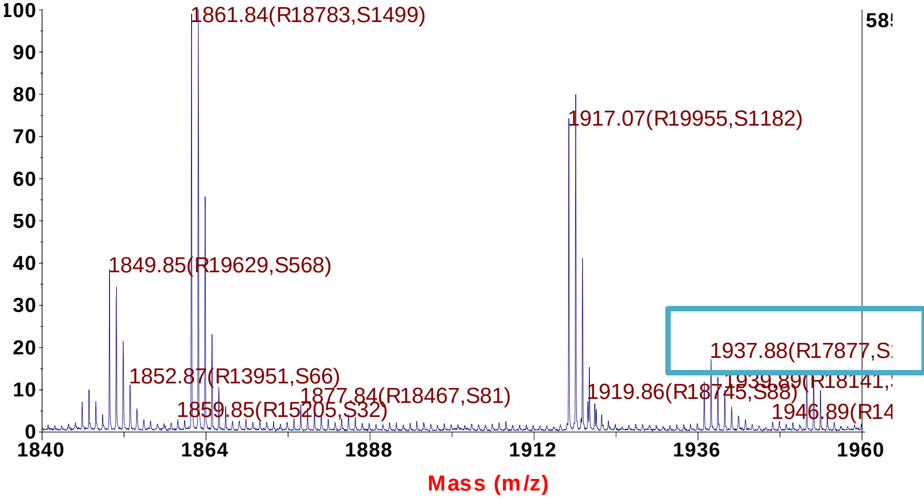

### Peptide 1935 Da

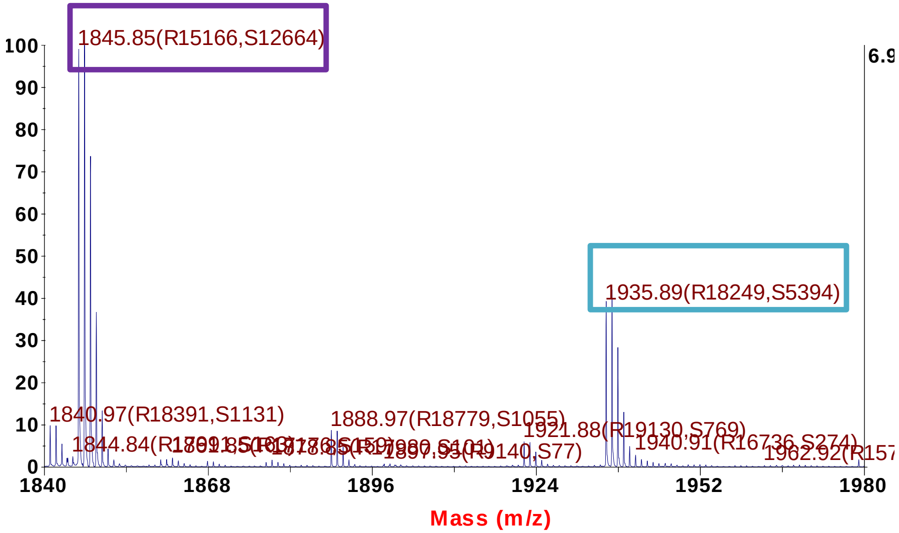


### Peptide 1935 Da

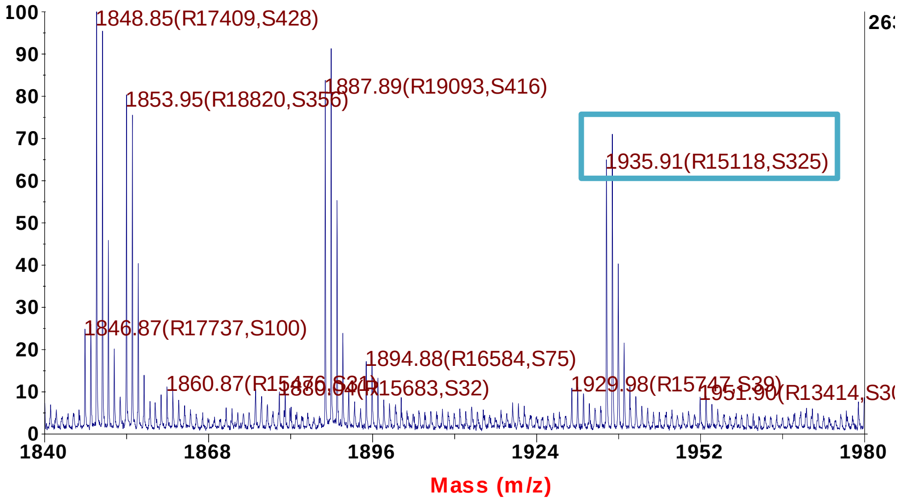

Pages 17-26 show MS2 spectra from LC-ESI MS/MS (liquid chromatography coupled to electrospray mass spectrometry) of peptides obtained after tryptic digestion of protein XylR. Amino acids modified with BrBz (+90 Da) are highlighted in red.

- Used chromatographic system: Dionex Ultimate 3000
- Mass spectrometer: Bruker HCT Ultra ion trap
- Gradient lenght: 90 min

A

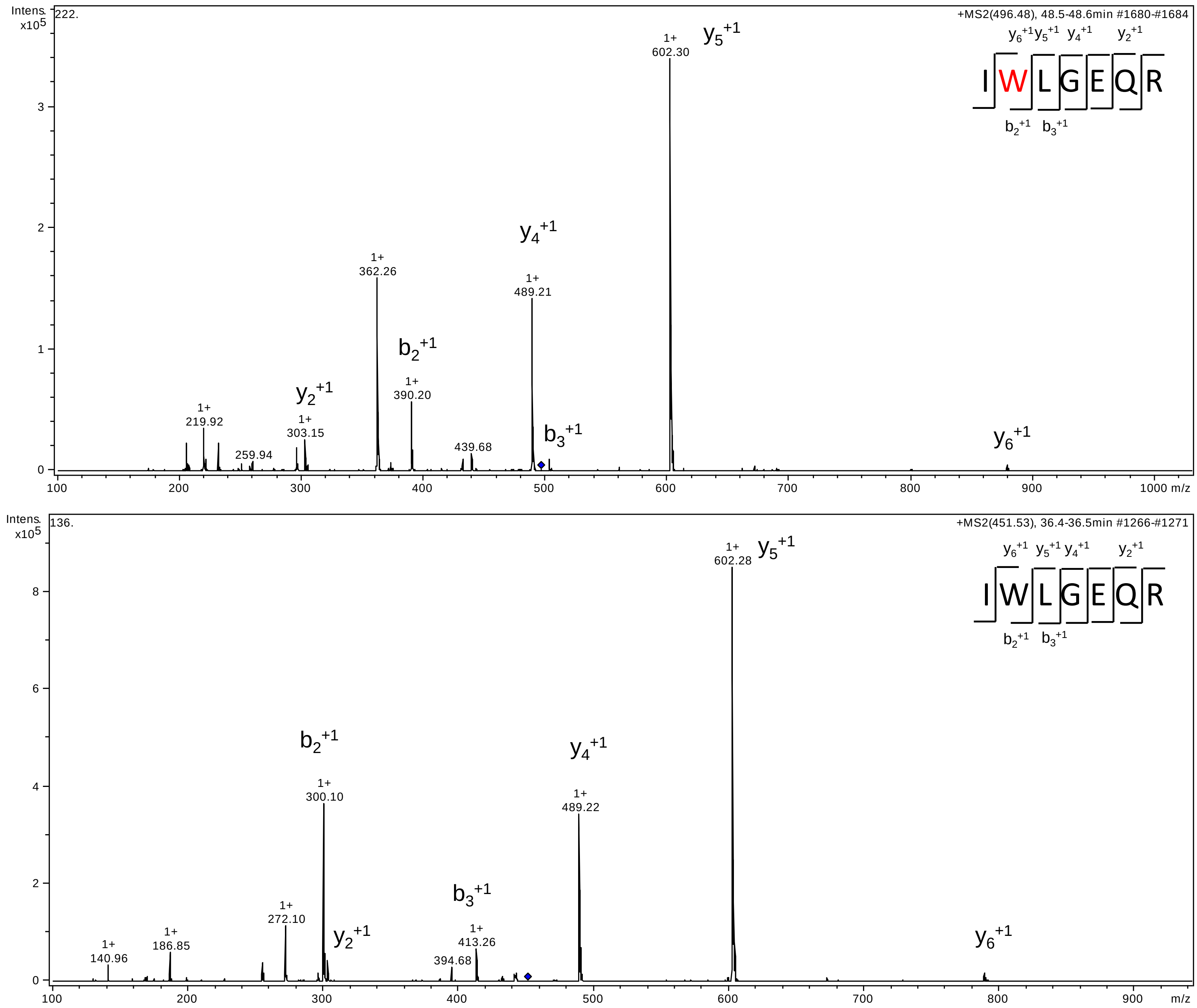

Intens. x10^5
222.
+MS2(496.48), 48.5-48.6min #1680-#1684
IWLGEQR
y6+1 y5+1 y4+1 y2+1
b2+1 b3+1
1+ 602.30
1+ 362.26
1+ 489.21
1+ 390.20
1+ 219.92
1+ 303.15
259.94
439.68
m/z
Intens. x10^5
136.
+MS2(451.53), 36.4-36.5min #1266-#1271
IWLGEQR
1+ 602.28
1+ 300.10
1+ 489.22
1+ 272.10
1+ 186.85
1+ 140.96
1+ 413.26
394.68
m/z

B

Extracted ion-chromatogram of the MS signal (y-axis) corresponding to unmodified IWLGEQR (green line) or IW(BrBz)LGEQR (blue line) and their respective HPLC retention times

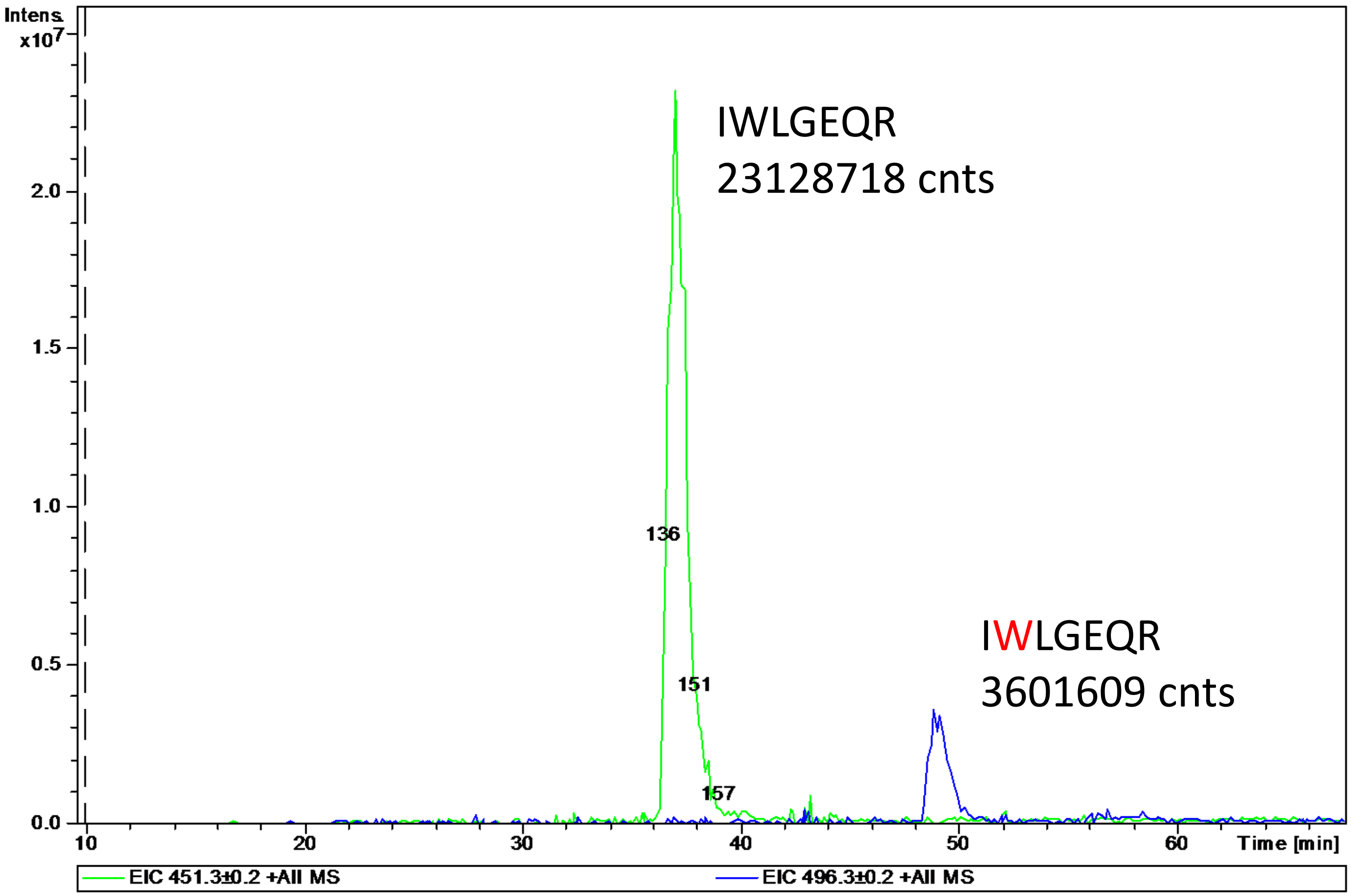

A

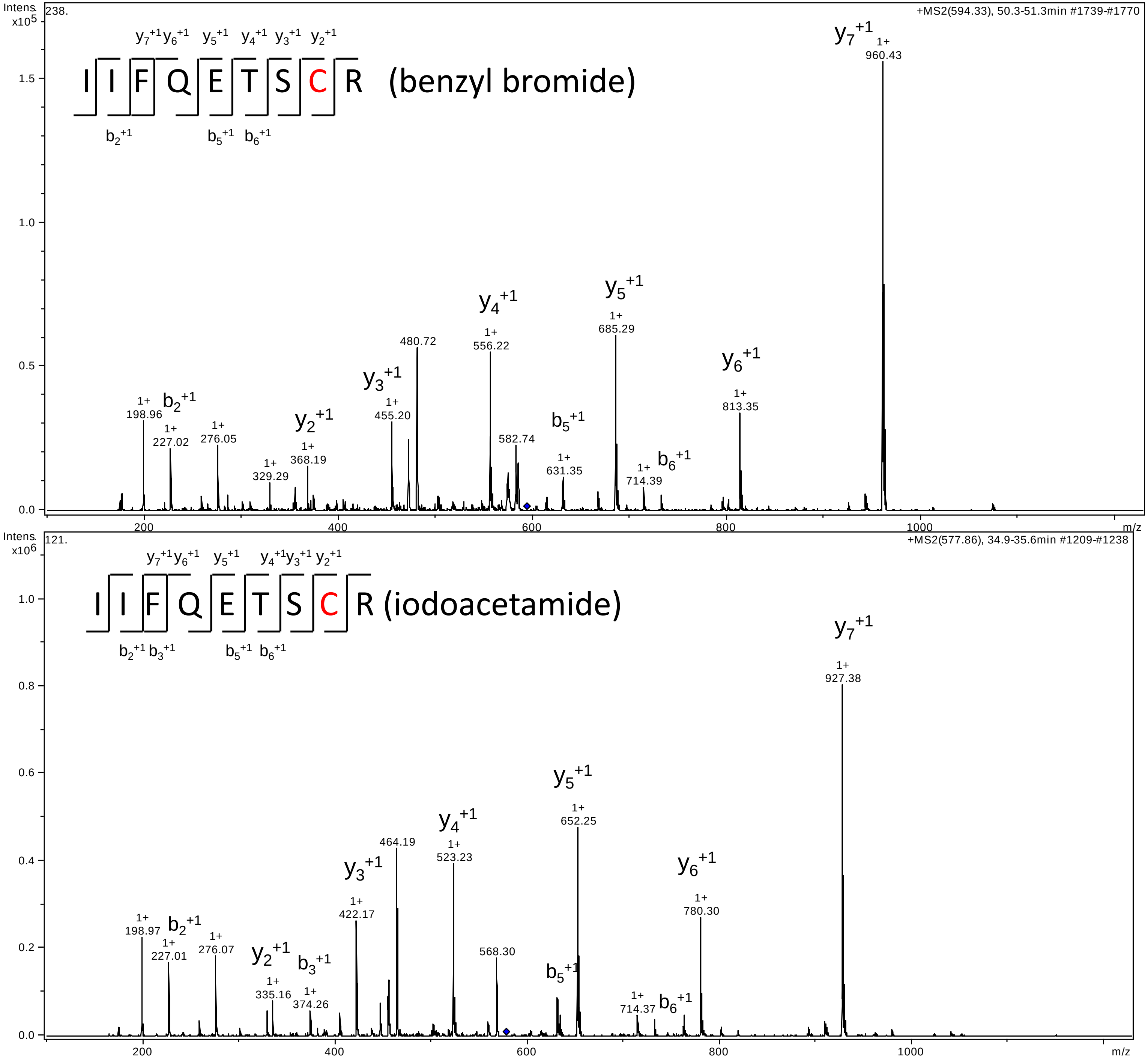

Intens. x10^5
238.
+MS2(594.33), 50.3-51.3min #1739-#1770
I I F Q E T S C R (benzyl bromide)
y7+1 y6+1 y5+1 y4+1 y3+1 y2+1
b2+1 b5+1 b6+1
1.5
1.0
0.5
0.0
y7+1
1+ 960.43
y5+1
1+ 685.29
y4+1
1+ 556.22
480.72
y6+1
1+ 813.35
y3+1
1+ 455.20
b2+1
1+ 198.96
1+ 227.02
1+ 276.05
y2+1
1+ 368.19
1+ 329.29
582.74
b5+1
1+ 631.35
b6+1
1+ 714.39
200
400
600
800
1000
m/z
Intens. x10^6
121.
+MS2(577.86), 34.9-35.6min #1209-#1238
I I F Q E T S C R (iodoacetamide)
y7+1 y6+1 y5+1 y4+1 y3+1 y2+1
b2+1 b3+1 b5+1 b6+1
1.0
0.8
0.6
0.4
0.2
0.0
y7+1
1+ 927.38
y5+1
1+ 652.25
y4+1
1+ 523.23
464.19
y3+1
1+ 422.17
y6+1
1+ 780.30
b2+1
1+ 198.97
1+ 227.01
1+ 276.07
y2+1
1+ 335.16
b3+1
1+ 374.26
568.30
b5+1
1+ 714.37
b6+1
200
400
600
800
1000
m/z

B

Extracted ion-chromatogram of the MS signal (y-axis) corresponding to IIFQETSC(ia)R (blue line) or IIFQETSC(BzBr)R (green line) and their respective HPLC retention times

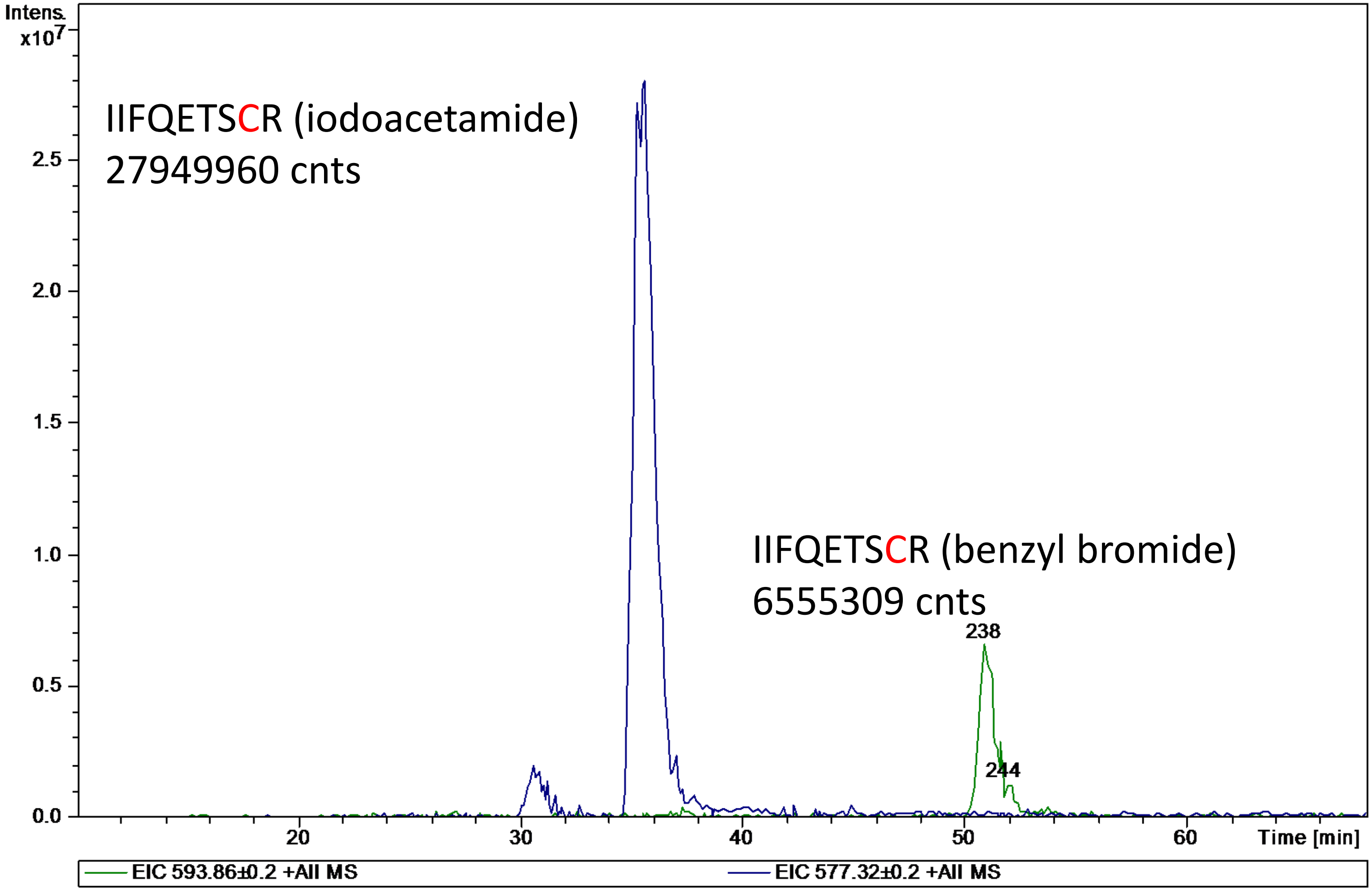

A

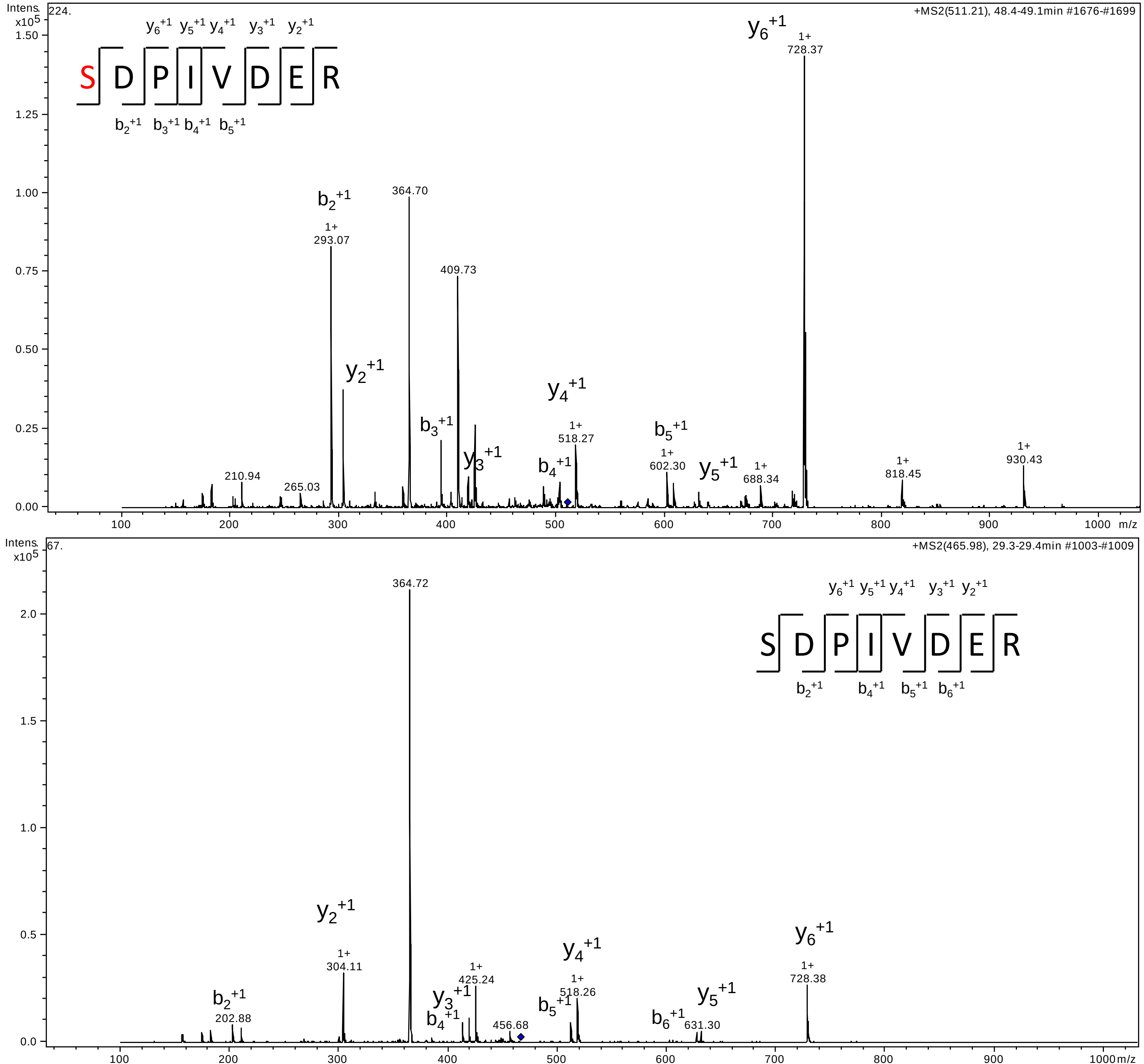

Intens. x10^5
224.
+MS2(511.21), 48.4-49.1min #1676-#1699
SDPIVDER
y6+1 y5+1 y4+1 y3+1 y2+1
b2+1 b3+1 b4+1 b5+1
y6+1 1+ 728.37
364.70
b2+1 1+ 293.07
409.73
y2+1
y4+1 1+ 518.27
b3+1
b5+1 1+ 602.30
y3+1
b4+1
y5+1
1+ 688.34
1+ 818.45
1+ 930.43
210.94
265.03
m/z
Intens. x10^5
67.
+MS2(465.98), 29.3-29.4min #1003-#1009
364.72
SDPIVDER
y6+1 y5+1 y4+1 y3+1 y2+1
b2+1 b4+1 b5+1 b6+1
y2+1 1+ 304.11
y6+1 1+ 728.38
y4+1 1+ 518.26
y3+1 1+ 425.24
b2+1 202.88
b4+1
456.68
b5+1
y5+1
b6+1 631.30
m/z

B

Extracted ion-chromatogram of the MS signal (y-axis) corresponding to unmodified SDPIVDER (green line) or S(BzBr)DPIVDER (red line) and their respective HPLC retention times

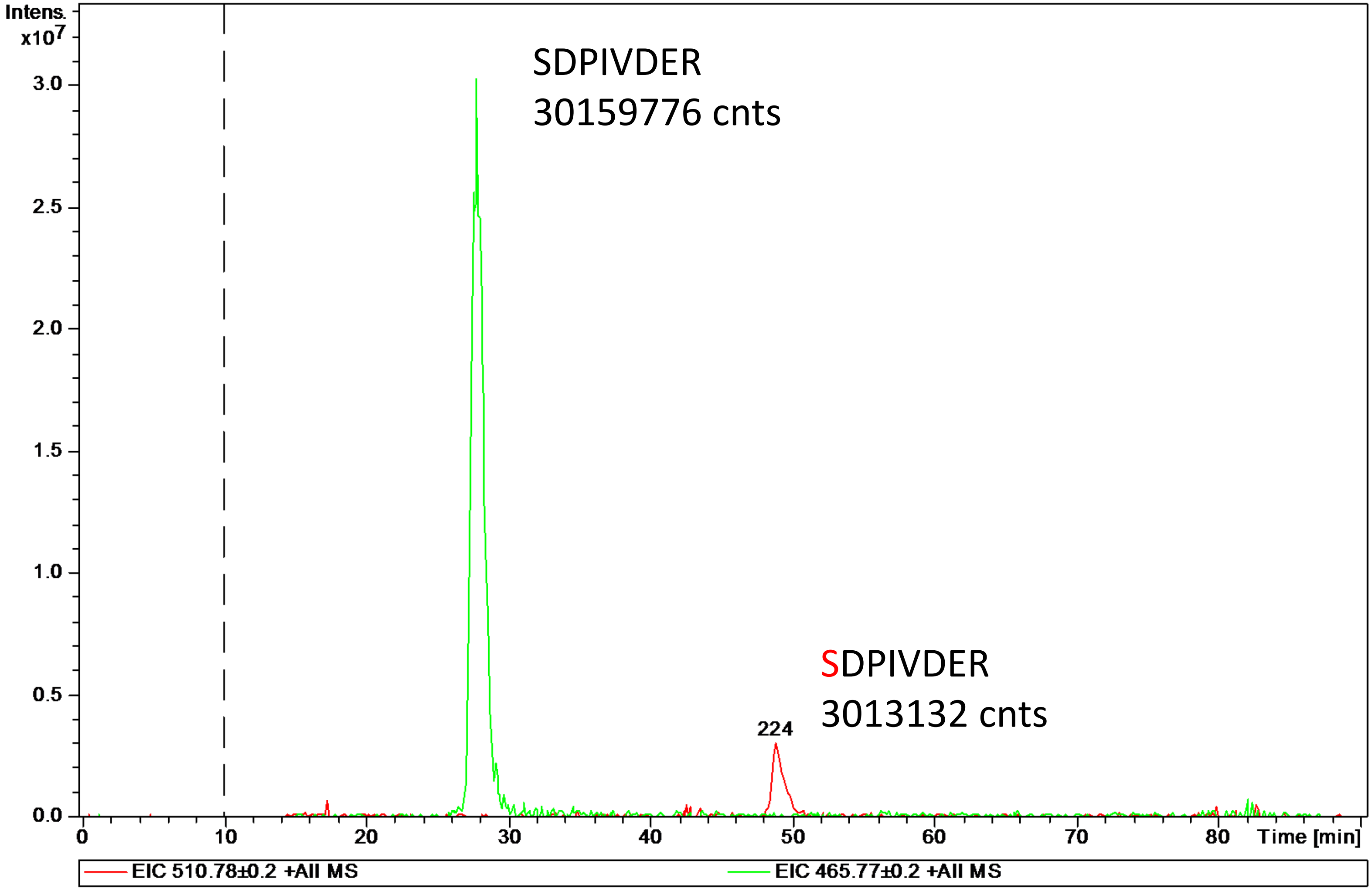

A

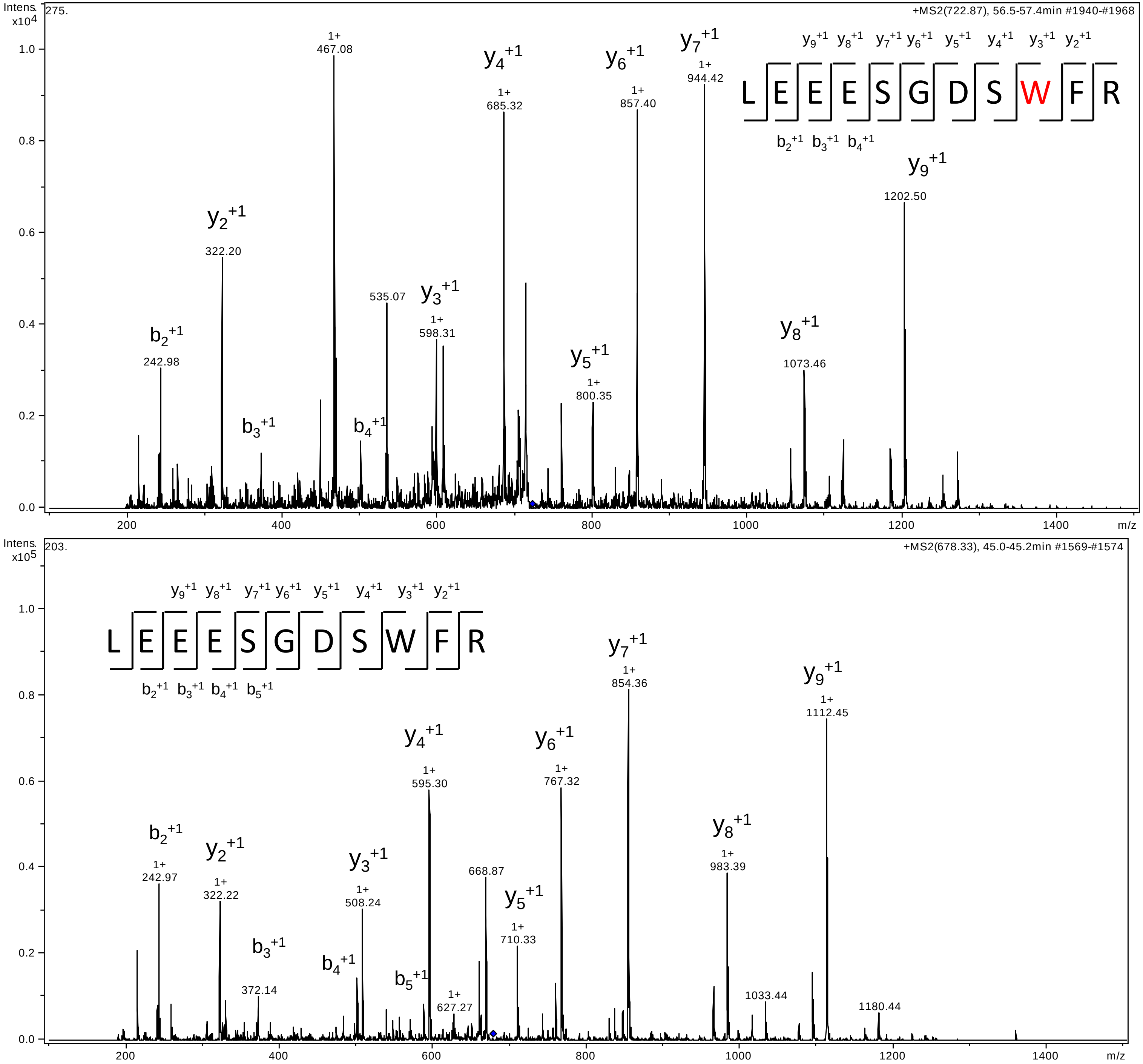
Intens. x10^4
275.
+MS2(722.87), 56.5-57.4min #1940-#1968
L E E E S G D S W F R
y9+1 y8+1 y7+1 y6+1 y5+1 y4+1 y3+1 y2+1
b2+1 b3+1 b4+1
b2+1 242.98
y2+1 322.20
b3+1
1+ 467.08
b4+1
535.07
y3+1 1+ 598.31
y4+1 1+ 685.32
y5+1 1+ 800.35
y6+1 1+ 857.40
y7+1 1+ 944.42
y8+1 1073.46
y9+1 1202.50
m/z
Intens. x10^5
203.
+MS2(678.33), 45.0-45.2min #1569-#1574
L E E E S G D S W F R
y9+1 y8+1 y7+1 y6+1 y5+1 y4+1 y3+1 y2+1
b2+1 b3+1 b4+1 b5+1
b2+1 1+ 242.97
y2+1 1+ 322.22
b3+1 372.14
b4+1
y3+1 1+ 508.24
y4+1 1+ 595.30
b5+1
1+ 627.27
668.87
y5+1 1+ 710.33
y6+1 1+ 767.32
y7+1 1+ 854.36
y8+1 1+ 983.39
1033.44
y9+1 1+ 1112.45
1180.44
m/z

B

Extracted ion-chromatogram of the MS signal (y-axis) corresponding to unmodified LEEESGDSWFR (black line) or LEEESGDSW(BzBr)FR (grey line) and their respective HPLC retention times

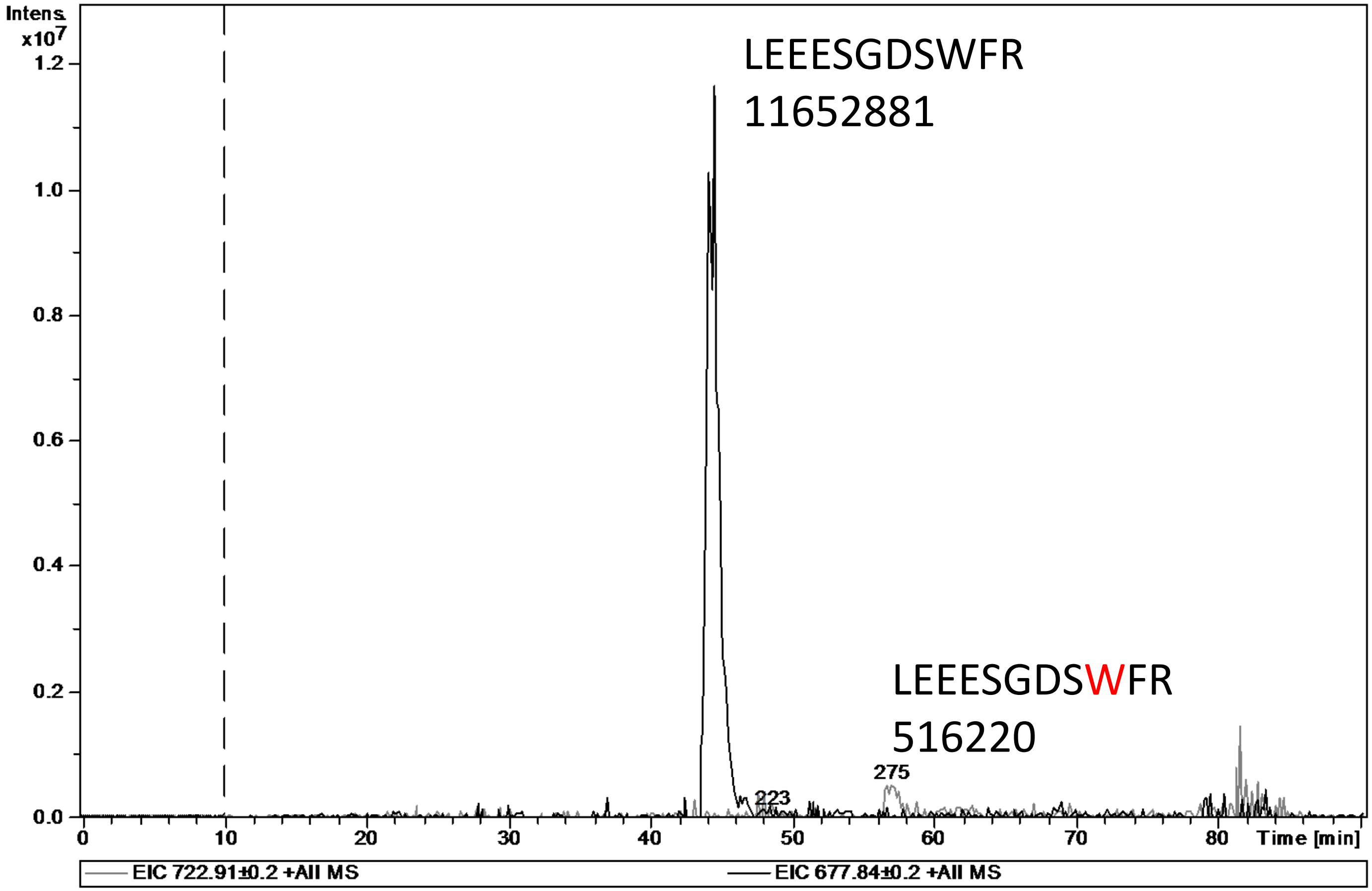

A

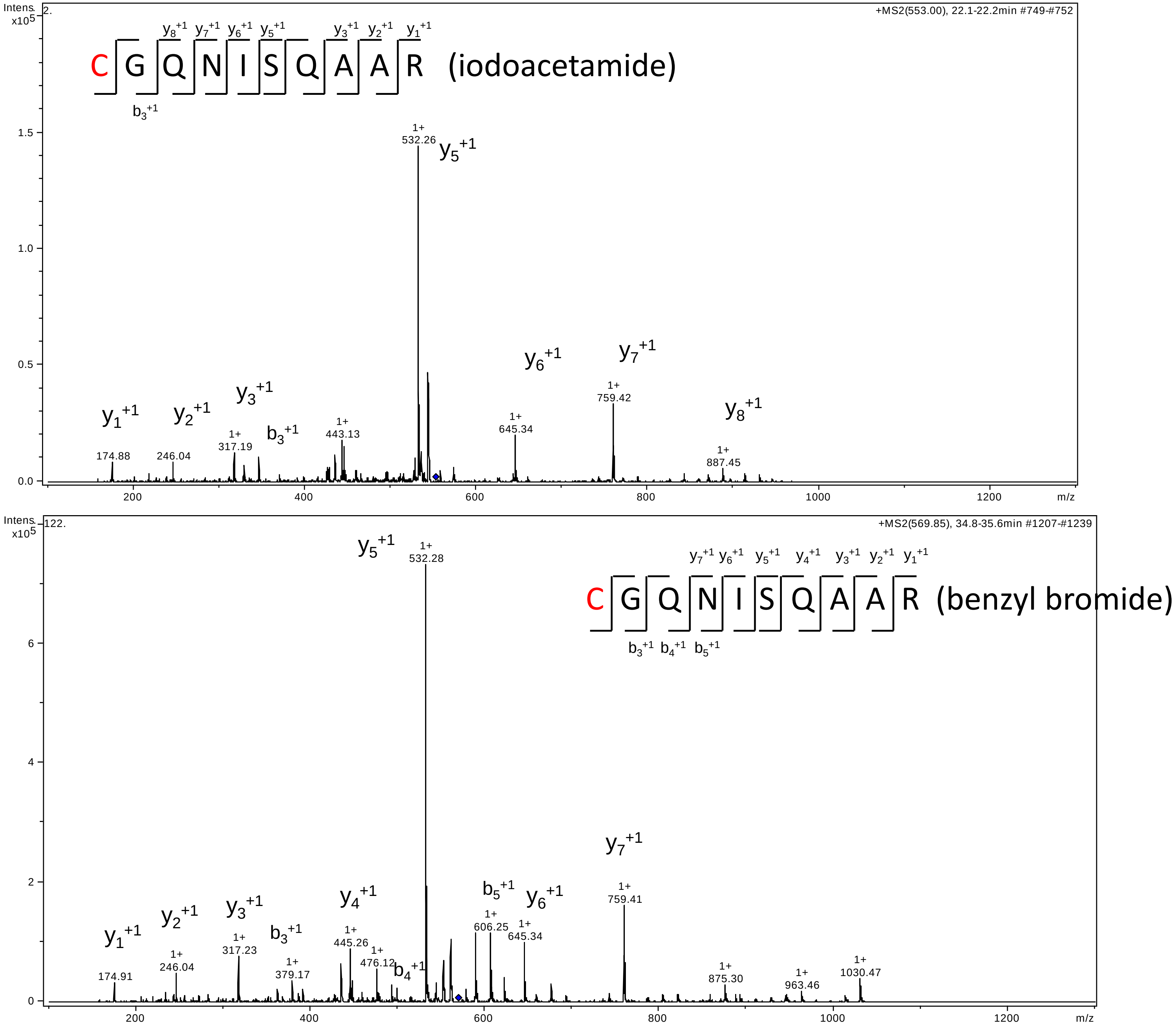

Intens. x10^5
2.
+MS2(553.00), 22.1-22.2min #749-#752
y8+1 y7+1 y6+1 y5+1 y3+1 y2+1 y1+1
C G Q N I S Q A A R (iodoacetamide)
b3+1
1+ 532.26 y5+1
y6+1
y7+1
1+ 759.42
1+ 645.34
y1+1
y2+1
y3+1
b3+1
y8+1
1+ 887.45
1+ 443.13
1+ 317.19
174.88
246.04
1.5
1.0
0.5
0.0
200
400
600
800
1000
1200
m/z
Intens. x10^5
122.
+MS2(569.85), 34.8-35.6min #1207-#1239
y5+1
1+ 532.28
y7+1 y6+1 y5+1 y4+1 y3+1 y2+1 y1+1
C G Q N I S Q A A R (benzyl bromide)
b3+1 b4+1 b5+1
y7+1
1+ 759.41
b5+1
y6+1
1+ 606.25
1+ 645.34
y4+1
1+ 445.26
1+ 476.12
b4+1
y3+1
b3+1
y2+1
y1+1
1+ 317.23
1+ 379.17
1+ 246.04
174.91
1+ 875.30
1+ 963.46
1+ 1030.47
6
4
2
0
200
400
600
800
1000
1200
m/z

B

Extracted ion-chromatogram of the MS signal (y-axis) corresponding to C(ia)GQNISQAAR (green line) or C(BrBz)GQNISQAAR (purple line) and their respective HPLC retention times

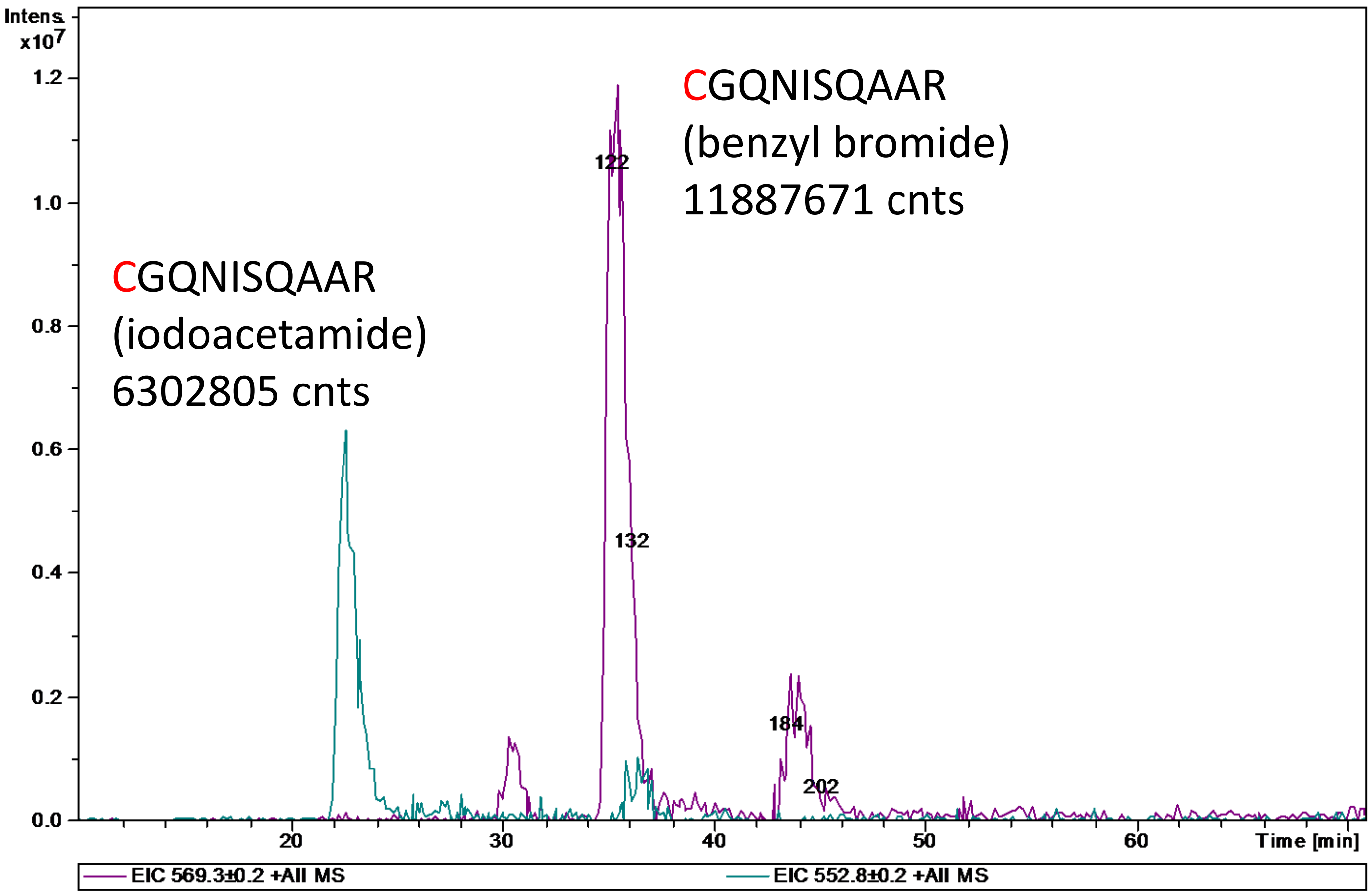
